# Linking human brain functional connectivity to underlying neurotransmitter systems

**DOI:** 10.64898/2026.04.28.721294

**Authors:** Leon D. Lotter, Golia Shafiei, Daouia Larabi, Abhay Koushik, Ottavia Dipasquale, Mitul Mehta, Mara Cercignani, Arjun Sethi, Neil Harrison, Štefan Holiga, Daniel Umbricht, Igor Yakushev, Suresh Muthukumaraswamy, Anna Forsyth, Joerg F. Hipp, Bratislav Misic, Svenja Caspers, Julian Koenig, Kaustubh R. Patil, Casey Paquola, Simon B. Eickhoff, Juergen Dukart

## Abstract

The human brain is organized into interacting functional systems. Their underlying neurobiological mechanisms remain difficult to study in vivo. Here, we adopt a topological framework to quantify the association between neurobiology and brain functional connectivity derived from both resting-state fMRI and MEG. Across six healthy adult cohorts (n = 19–112), regional variation in fMRI connectivity robustly aligns with the distribution of neurotransmitter receptors and transporters. These patterns are present in every single subject, replicate across all cohorts, and are mirrored in MEG. Most prominently, noradrenergic modulation of connectivity in a sensorimotor-insular network is consistently detected across individuals and linked to autonomic arousal. In pharmacological and clinical samples, associations are sensitive to neurotransmitter system manipulation and are altered in early psychosis, aligning with clinical symptomatology. These findings provide biological insight into typical and atypical functional organization of the human brain using a framework linking underlying neurobiology to the functional connectome (NEOFC).

## Introduction

Resting-state functional connectivity (rsFC)^1^ has become the primary framework for characterizing brain functional organization. Resting-state FC networks and, more recently, cortical gradients have been identified across various neuroimaging modalities and are now considered a core feature of functional brain architecture^2–5^. A central assumption in rsFC research is that the observed temporal synchrony between regions is driven by common neural activity^6^. However, despite three decades of study, the biological basis of rsFC – characterized through network, gradient, and related perspectives – remains only partly understood.

In humans, evidence linking macro-scale rsFC networks to specific cellular and molecular mechanisms is largely indirect, often extrapolated from non-human animal studies^7,8^, and constrained by the limited neurobiological specificity of in vivo connectivity measures: electroand magnetoencephalography (E/MEG) capture aggregated neural activity whose cellular and neurotransmitter origins remain difficult to dissociate^9^, blood-oxygenation-level-dependent (BOLD) functional magnetic resonance imaging (fMRI) reflects cumulative local metabolic demand shaped by neurovascular coupling and other non-neuronal physiological processes^6,7^. As a result, rsFC alone offers limited information on the neurobiological processes that may underlie functional variation across groups, and even more so, across individuals. Here, we introduce a framework that quantifies the spatial association between individual rsFC and reference atlases of neurobiological tissue properties. The revealed associations are robust, generalizable, related to other physiological processes, modified by intervention, and affected by clinical conditions, providing a biologically grounded perspective for the study of macro-scale functional organization of the human brain.

In recent years, we and others have shown that the spatial topology of resting-state brain activity and connectivity derived from haemodynamic and electrophysiological data is associated with specific neurobiological features^10–23^. For example, the magnitude of blood flow changes induced by dopaminergic medication follows the expected distributions of dopaminergic receptors in the brain^10^. Pharmacological modulation of regional BOLD signals can be understood through the expected distributions of the respective drug targets^22,24^. BOLD alterations in neurological and neurodevelopmental disorders colocalize with affected neurotransmitter systems^13–16^. Normative connectome organization is related to inter-regional similarity in receptor and transporter expression^17^, and neurophysiological network centrality colocalizes with major transmitter receptors^20^.

The cross-modal colocalization approaches employed by prior studies aggregated information within regions, across neurotransmitter systems, or across subjects, disregarding information on individual brain network organization. We argue that the assumption underlying the common region-level approaches – if a receptor’s activity contributes to the measured neural signal, then modulating receptor activity should modulate the signal in brain regions with higher receptor availability – extends to functional connectivity: if a receptor is active across multiple brain regions contributing to the measured signal, its activity should introduce a characteristic pattern to the regional signals, which may result in increased synchronization between regions with high receptor abundance. We hypothesize that the inverse may hold for receptors supporting local, short-range signaling, whose relative absence may facilitate interregional synchronization.

Building on these theoretical considerations, here we introduce a framework for evaluating the Neurobiologically Enriched Organization of Functional Connectivity (NEOFC). We quantify changes in rsFC as a function of regional receptor and transporter density estimates, obtained from nuclear imaging in healthy volunteers and spanning all major neurotransmitter systems^25,26^. A derived subject-level index aggregates the topological relationship between an individual rsFC connectome and specific neurobiological tissue properties. In multiple independent resting-state fMRI (rsfMRI) datasets, we show that these indices of neurobiology-dependent functional synchronization capture a robust and reliable component of brain network organization at both group and individual levels. Extended effects in MEG connectomes demonstrate that this network-organizational component generalizes across modalities. Close links between rsFC in a noradrenaline-rich sensorimotor-insular network and autonomic arousal support the neurobiological relevance of the framework. Sensitivity to pharmacological manipulation and to rsFC alterations in early psychosis further illustrate its clinical potential. All methods are openly available for the community to build upon.

## Results

### A framework linking typical neurobiological signals to individual connectomes

The NEOFC framework (Fig. 1, Animation S1) quantifies the topological association between the spatial distribution of a neurobiological marker across brain regions (reference atlas; here, a group-average nuclear imaging map; Fig. 1a) and the functional network defined over the same regions (connectome; here, an individual’s 200-region-by-region Pearson rsFC matrix; Fig. 1b). We obtain this association by (i) converting the reference atlas to percentiles, (ii) masking the connectome according to atlas percentile thresholds ranging from 0 to 95, (iii) calculating the mean rsFC between regions remaining after each threshold step, and (iv) condensing the result in a curve of percentile thresholds versus mean rsFC (Fig. 1c). This relationship is then summarized as a single score by calculating the area under the curve (AUC) after subtracting the global connectivity (mean rsFC at percentile 0, i.e. of all connections; Fig. 1d). These AUC scores capture the overall association between each individual connectome and reference atlas and are used for all main and follow-up analyses (Fig. 1e). The AUC serves as the primary aggregate metric because it captures the overall connectome-reference relationship independent of curve shape. Using this approach, we test for positive associations, such that synchronization is stronger between regions with higher density of a given neurobiological atlas (AUC+). We then repeat the analysis using the spatial inverse of each reference atlas to test for stronger synchronization between low-density regions (AUC−; Fig. 1a–d). To guard against spurious effects driven by shared spatial structure^27^, each association is evaluated against null reference atlases preserving spatial autocorrelation and interhemispheric symmetry^25^. This allows us to calculate non-parametric exact p values on group and individual levels, which we correct for the effective number of tests across atlases within each main analysis (p_Meff_, see Methods). In sensitivity analyses, we evaluated both alternative brain parcellations and an alternative aggregation score focused on the quadratic-like shape of some percentile-rsFC curves (second-degree polynomial fit; Supplementary Methods).

**Fig. 1:**
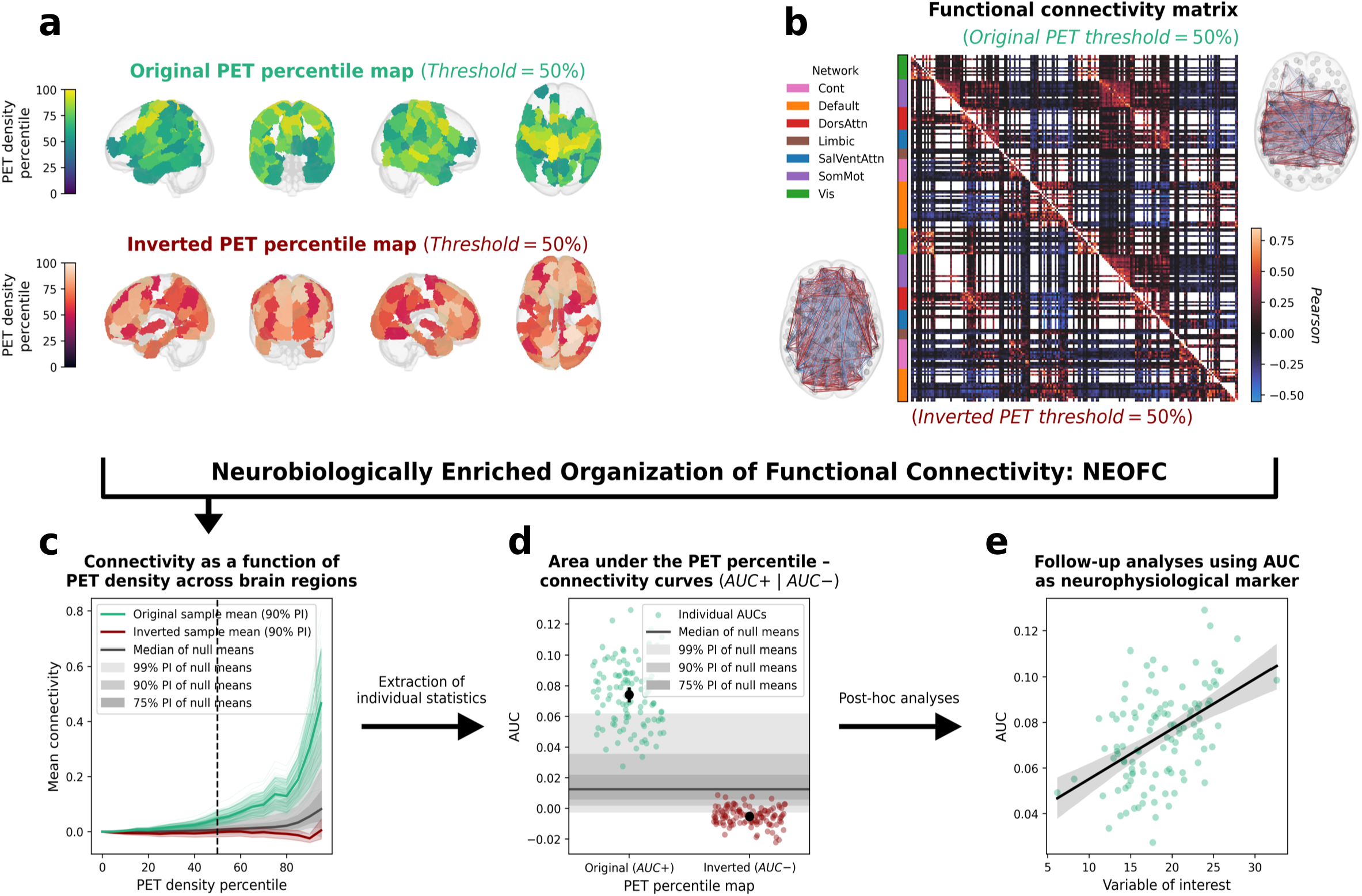
Neurobiologically enriched organization of functional connectivity (NEOFC) **a**: Cortical parcels (Schaefer200 parcellation, n = 200) color-coded by percentile rank of a given reference atlas (here: the noradrenaline transporter); percentile threshold 50% shown as an example. **b**: An rsFC matrix (here: group average correlation strength) thresholded by the 50th percentile of the reference atlas (top 50% densest parcels: top right; bottom 50%: bottom left); brain plots: thresholded connectomes projected on a standard glass brain. **c**: NEOFC curve: mean rsFC of included parcels as a function of the density percentile threshold, computed for the original reference map (green) and the spatially inverted reference map (dark red). Colored shaded area: group mean ± 90% PI across individuals. Grey shaded area: null distribution based on null models of the reference map. **d**: Individual AUC+ and AUC− scores, quantifying the area under the original and inverted NEOFC curves, respectively (each dot: one participant). **e**: Illustrative downstream application: AUC+ scores associated with a continuous variable of interest via linear regression (scatter: individual participants; line: regression fit). Abbreviations: rsFC: resting-state functional connectivity; PI: percentile interval; AUC: area under the curve.

We assembled a database of 25 reference atlases derived from independent healthy adult nuclear imaging studies^25,26^, using a harmonized registration procedure to ensure comparability and spanning all major neurotransmitter systems as well as metabolism, synaptic density, and transcriptomic activity (Tab. S1). For a positive control analysis, we additionally obtained probabilistic atlases of 12 established resting-state networks^28^. Individual connectomes were derived from rsfMRI and source-reconstructed MEG data from a subset of the Human Connectome Project Young Adult sample (HCP-YA)^29^, alongside rsfMRI data from one physiology study, three pharmacological studies, and one clinical case-control study (Tab. S2). Note that reference atlases are group-averaged maps derived from independent cohorts and are not subject-matched to individual connectomes. The HCP-YA rsfMRI dataset served as the discovery sample for the main analyses (n = 112, 64 female), and findings were tested for replication in healthy, untreated, adult subsamples from the other five rsfMRI datasets.

### NEOFC is sensitive to canonical resting state network structure

The canonical organization of the resting functional connectome is described by the well-established resting-state networks^2,6^. Probabilistic atlases map the probability of a region to belong to a given network, i.e., regions with high probability for a given network are likely to be connected to each other. In the NEOFC framework, such atlases should yield strong effects, acting as a positive control. In line with this, we found significant AUC+ effects for 8 out of 12 tested resting-state networks in the HCP-YA sample (p_Meff_ < 0.05; Figs. 2a/b; Tab. S2), indicating increased rsFC between regions with high probability of a given network. Interestingly, 5 networks, including 3 of the 4 which did not show AUC+ effects, also showed significant AUC− effects, indicating stronger synchronization between regions with low network probability and likely reflecting the known spatial anticorrelation of some of the networks (p_Meff_ < 0.05; Fig. 2a/b). This pattern was generally robust to the alternative parcellation schemes and aggregation metric (Fig. S1; Tab. S3).

**Fig. 2:**
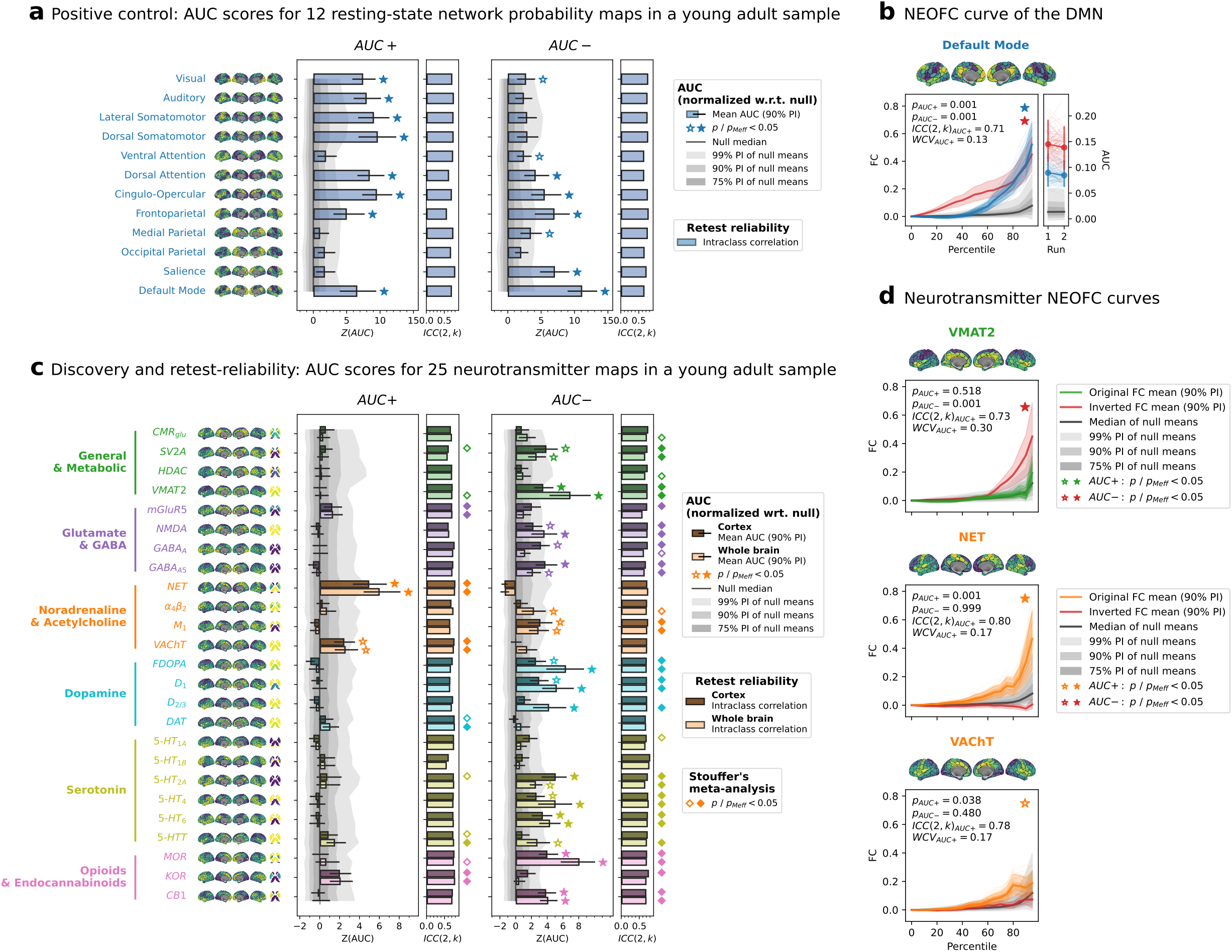
Resting-state fMRI functional connectivity organizes along neurobiology. **a**: AUC+ and AUC− scores (z-scored against a null distribution of spatially matched reference maps) for 12 resting-state network atlases used as a positive control, together with test-retest reliability (ICC(2,k); Schaefer200 parcellation, first scanning run). Bars: group mean; error bars: 90% PI across individuals; grey shaded areas null distributions of the AUC group means, z-scored similarly to the observed scores for visualization. Brain maps show the parcellated reference atlases converted to percentiles. **b**: Exemplary NEOFC curve for the DMN probability atlas during the first scanning run (original percentile-FC curve: blue; inverted test: dark red; colored shaded area: 90% PI across individuals; grey shaded area: null distribution). Right side: comparison of AUC estimates from run 1 and 2 (thin lines: individual subjects; thick line, dots, and errorbars: group mean with 90% PI). **c**: AUC+ and AUC− scores for 25 nuclear imaging reference maps, displayed separately for Schaefer200 (darker bars) and Schaefer200 + 16 subcortical parcels (brighter bars); shown as in **a**. **d**: NEOFC curves for three selected monoaminergic transporter maps (VMAT2, NET, VAChT); displayed as in **b**. Abbreviations: AUC: area under the curve; ICC: intraclass correlation coefficient; WCV: within-subject coefficient of variability; PI: percentile interval; pMeff: p-value corrected for effective number of comparisons; DMN: default mode network; CMR_glu_: cerebral metabolic rate of glucose; SV2A: synaptic vesicle protein 2A; HDAC: histone deacetylase; VMAT2: vesicular monoamine transporter 2; mGluR5: metabotropic glutamate receptor 5; NMDA: N-methyl-D-aspartate receptor; GABA_A, A5_: gamma-aminobutyric acid type A, A5 receptor; NET: noradrenaline/norepinephrine transporter; α4β2: alpha-4 beta-2 nicotinic receptor; M_1_: muscarinic acetylcholine receptor M1; VAChT: vesicular acetylcholine transporter; FDOPA: fluorodopa; D_1, 2/3_: dopamine receptor 1, 2/3; DAT: dopamine transporter; 5-HT_1A, 1B, 2A, 4, 6_: serotonin receptor 1A, 1B, 2A, 4, 6; MOR/KOR: mu/kappa-opioid receptor; CB1: cannabinoid receptor type 1.

### The fMRI resting-state connectome is robustly associated with neurobiological signals

Having established that NEOFC can capture known topological relationships, we next tested for the neurobiological signals underlying rsFC organization in the HCP-YA fMRI dataset. We observed the strongest AUC+ effect for the noradrenaline transporter (NET; p_Meff_ < 0.05; Fig. 2c/d; Fig. S2; Tab. S4), arising from higher connectivity between regions with stronger NET signals. The effect was significant in each of 112 discovery subjects (p < 0.05; Fig. S3) and in all five replication samples (p_Meff_ < 0.05; Fig. S4). A meta-analysis across all six healthy datasets, suited to also detect weaker effects, additionally revealed significant AUC+ effects of acetylcholine, dopamine, and serotonin transporters (VAChT, DAT, 5-HTT) as well as glutamate and opioid receptors (mGluR5, KOR; p_Meff_ < 0.05; Fig. 2c; Fig. S5; Tab. S5). In contrast to the few AUC+ effects, AUC–effects were observed across all studied neurotransmitter systems (p_Meff_ < 0.05; Fig. 2c/d; Tab. S4). This pattern of results was supported by analysis of the delta between AUC+ and AUC–effects, which should diminish influences of shared spatial patterns not captured by the null models (Fig. S6; Tab. S4). The main effects were robust to different parcellation schemes, removal of interhemispheric connections, as well as the alternative aggregation metric (Supplementary Results; Fig. S6/7; Tab. S4).

### Neurobiological rsFC organization is reliable and reproducible

One-day test-retest data from the HCP discovery dataset served to evaluate the reliability of the NEOFC framework. All significant AUC outcomes showed a good to excellent retest reliability, supporting robustness of the respective findings [ICC(2,k) > 0.7; within-subject coefficient of variation < 0.25; Fig. 2a; Tab. S6]. Comparison of the AUC+ and AUC–profiles between six independent datasets showed very good to excellent reproducibility (Fig. 3a; Fig. S4; Tabs. S7/8).

**Fig. 3:**
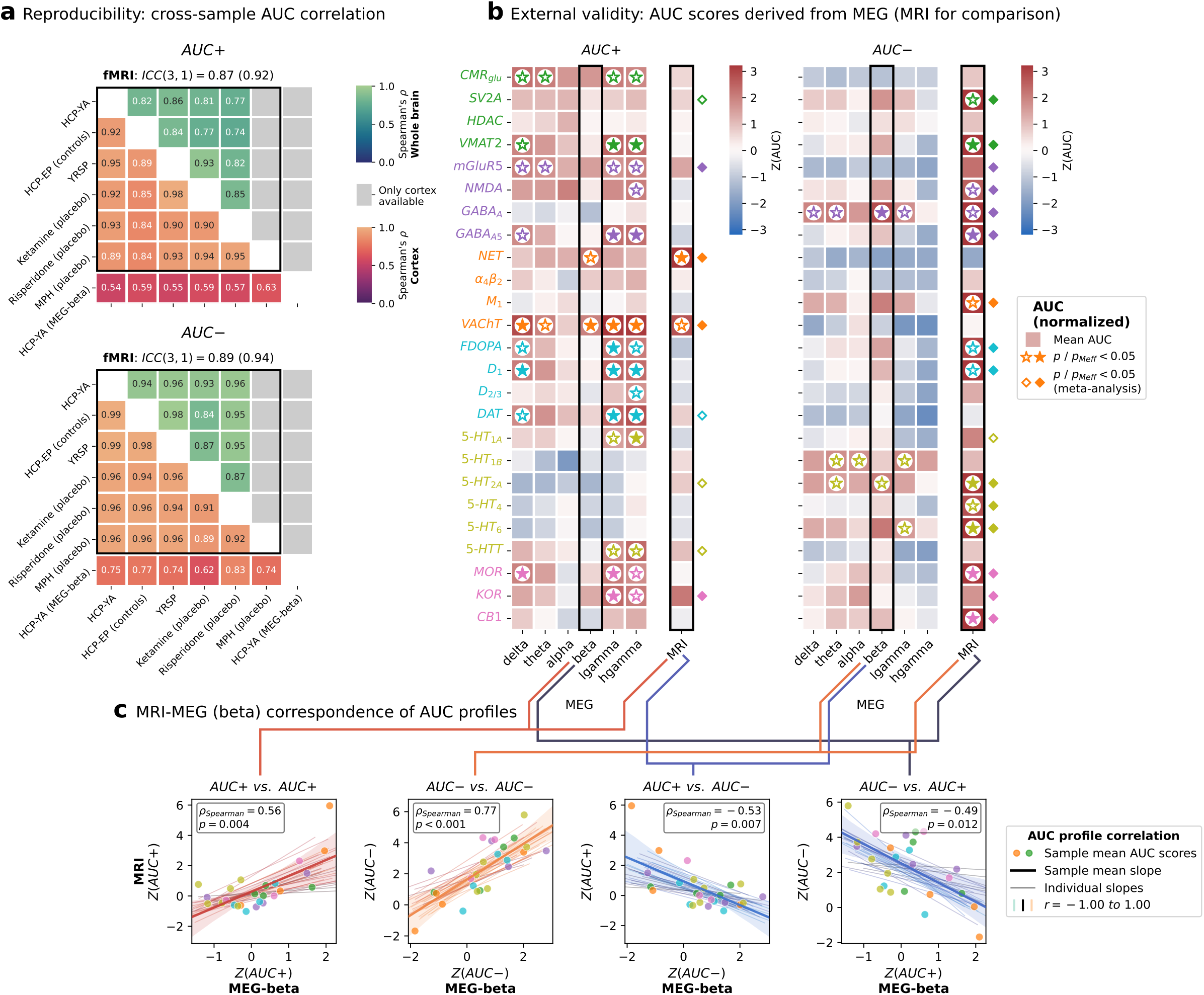
Neurobiological rsFC organization is reproducible and extends to MEG. **a**: Pairwise Spearman’s rho between mean AUC+ (top) and AUC− (bottom) profiles across 6 fMRI cohorts and one MEG cohort (25 nuclear imaging reference maps; Schaefer200 and Schaefer200 + subcortex parcellation). Lower triangle: cortex; upper triangle: whole-brain. ICC(3,1) reported for fMRI cohorts (first value: cortex; value in brackets: whole-brain). **b**: AUC+ and AUC− scores for 25 nuclear imaging reference maps derived from MEG AEC FC matrices, displayed as a heatmap (Schaefer200, z-scored against null distribution); right column: fMRI results for reference. **c**: Correspondence between fMRI and MEG-beta AUC profiles (z-scored against null) across 25 reference maps (each dot: sample-mean AUC value of one reference atlas; thick line: corresponding regression fit with 95% bootstrapped CI; thin lines: individual regression fits without single values; color: Spearman’s rho). Four combinations shown: fMRI AUC+ vs. MEG AUC+, fMRI AUC− vs. MEG AUC−, fMRI AUC+ vs. MEG AUC−, fMRI AUC− vs. MEG AUC+. Abbreviations: AUC: area under the curve; fMRI: functional magnetic resonance imaging; MEG: magnetoencephalography; ICC: intraclass correlation coefficient; FC: functional connectivity; AEC: amplitude envelope correlation; CI: confidence interval; pMeff: p-value corrected for effective number of comparisons.

### Electrophysiology replicates and extends hemodynamic findings

Electrophysiological measures are conceptually more closely related to neuronal activity than the hemodynamic signals measured by rsfMRI and may thus contain more specific information on interregional connectivity and its underlying biology. We extended our analysis of the HCP-YA dataset by studying source-reconstructed resting-state MEG connectomes across six frequency bands (amplitude envelope correlations; n = 33, 16 female)^21,30^. Strictly distance-dependent leakage is controlled by our spatial null models by construction; we applied additional pairwise orthogonalization^3^ in sensitivity analyses, expecting a general effect size reduction due to the unspecific removal of zero-lag signals (see Methods). Compared to rsfMRI, AUC profiles derived from MEG showed a shift towards higher sensitivity to AUC+ effects and reduced sensitivity to AUC–effects (Fig. 3b; Fig. S8; Tab. S9). Most AUC+ effects were observed in lowand highgamma frequency bands, spanning several neurotransmitter systems, while the beta band showed more sparse effects dominated by VAChT (p_Meff_ < 0.05) and NET AUC+ (nominal p < 0.05). In contrast to rsfMRI, AUC–results largely did not reach significance levels with the notable exception of GABA_A_ (p_Meff_ < 0.05). The results remained stable with different parcellation resolutions (Fig. S9; Tab. S9). Orthogonalization of connectomes led to reduced AUC+ effect sizes. AUC profiles remained significantly correlated with non-orthogonalized results. The NET beta-band effect specifically stayed nominally significant while VAChT’s did not, arguing against a leakage-driven origin of NET (Supplementary Results; Fig. S10; Tab. S9/S10).

Group- and within-subject level AUC+ and AUC–profiles displayed strong similarity between MEG and rsfMRI, with the strongest alignment observed for the MEG beta band (Fig. 3c; Fig. S11; Tab. S10). Between-subject correlations of individual AUC estimates per neurobiological atlas indicated only limited associations, possibly reflecting state-dependent variability between the non-concurrent MEG and rsfMRI sessions (n = 29; Fig. S12; Tab. S11).

### Region-specific influences on neurobiological rsFC organization

Next, we assessed how the general connectome-neurobiology associations captured by the AUC indices were influenced by rsFC patterns of individual brain regions. We employed a leaveone-out approach identifying regions/connections whose exclusion most strongly influenced the aggregated AUC metric as identified in the meta-analytic approach^31^. Second, we evaluated AUC scores calculated at systematically lowered upper percentile bounds to determine if the observed effects were driven by only a few regions with extreme expression or a continuous topological association across a larger subnetwork.

Both, AUC+ and AUC–displayed distinct topological associations (Figs. 4a and S13a/b). We found the NET AUC+ pattern to be most strongly influenced by posterior insula, lateral and medial sensorimotor, and supplementary motor areas, regions with notably high levels of NET (Figs. 4a and S13a). We furthermore found that the NET AUC+ effect emerged even when considering only the lowest 40% of the regional NET distribution. This indicates a widespread topo-logical brain network association, peaking in specific high-NET regions. AUC–scores derived from serotonergic and opioidergic atlases displayed even more extended patterns (Fig. S14; Tab. S12).

**Fig. 4:**
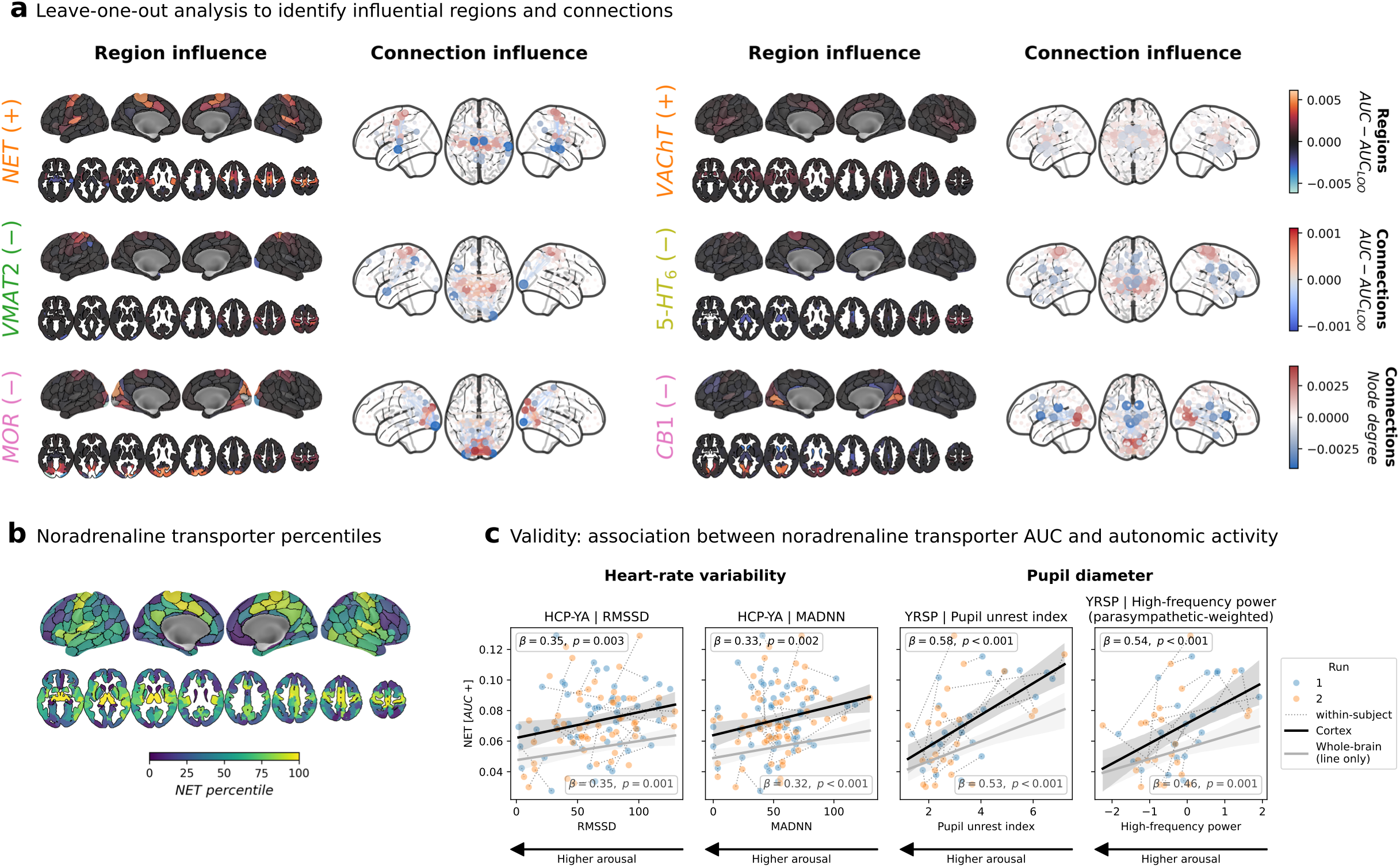
Neurobiological rsFC organization is driven by regional subnetworks and reveals a sensorimotor-insular component related to autonomic physiology. **a**: Regional (left, leave-one-region-out) and connection (right, leave-one-connection-out) influence on AUC for 6 selected reference maps and directions: NET (AUC+), VAChT (AUC+), VMAT2 (AUC−), 5-HT6A (AUC−), MOR (AUC−), and CB1 (AUC−) (Schaefer200 + subcortex parcellation). Color indicates direction and magnitude of influence. **b**: NET density percentile map projected onto the Schaefer200 + subcortex parcellation, shown for spatial reference. **c**: Association between NET AUC+ and autonomic physiological measures across two cohorts: HRV (RMSSD and MadNN; HCP-YA) and pupil-based arousal indices (PUI and HF-P power; YRSP). Scatter: individual observations, repeated measures are connected; black: Schaefer200 parcellation; grey: Schaefer200 + subcortex parcellation; line with shaded area: regression fit across all data with 95% bootstrapped CI. LMM used for statistical inference, controlling for mean FD, global connectivity, and demographic covariates. Abbreviations: rsFC: resting-state functional connectivity; AUC: area under the curve; HRV: heart rate variability; RMSSD: root mean square of successive differences; MadNN: median absolute deviation of NN intervals; PUI: pupil unrest index; HF-P: high-frequency power of the arousal-related pupil signal range (0.25–0.50 Hz); LMM: linear mixed model; FD: framewise displacement; CI: confidence interval; see Fig. 2 for reference map abbreviations.

So far, we introduced the NEOFC framework as a method to study topological associations between human brain connectome organization and neurobiological signals. The framework rests on the assumption that the spatial pattern exhibited by a neurobiological reference atlas is reflected in individual brain network topology. From the extent of this cross-modal spatial association, we derived interpretable markers of individual connectome-neurobiology relationships. These markers proved robust, reliable, and reproducible in rsfMRI data, and their spatial pattern was recovered in electrophysiological data, supporting a neuronal origin. In the following, we will investigate the (patho-)physiological relevance of our approach by exploring the notable noradrenergic effect, demonstrating that derived indices are sensitive to pharmacological modulation, and conducting a clinical application experiment.

### A noradrenergic motor-insular component reflects autonomic arousal

AUC+ associations with NET were the strongest and most consistent signal detected in the above analyses. We next tested for their neurobiological relevance, thereby providing external evidence supporting the NEOFC framework. The NET is typically located presynaptically at cortical and subcortical projection sites of noradrenergic neurons^32^, a subpopulation of which form the locus coeruleus (LC). Accordingly, the NET reference atlas showed the highest density in lateral and medial sensorimotor cortices, insula, and medial subcortical regions (Fig. 4b), the cortical regions of which strongly influenced the AUC+ effects.

On the basis of this anatomical pattern, the well-established autonomic functions of the LC^33^, the consistent NET effect across subjects and samples, and the observed regional importance profile, we hypothesized that the signal reflects a general neurobiological process related to autonomic arousal. Considering this hypothesis, the similarity of the NET atlas to the sensorimotor-association axis, and prior reports on a global fMRI component associated with peripheral physiology^34^, we tested the robustness of this association to spatial regression of atlases capturing brain vasculature, functional, and microstructural organization (Fig. S15; Tab. S13). The NET association remained nominally significant throughout (p < 0.05), though attenuated for several covariates – most strongly by the sensory-association axis and two principal gradients (p_Meff_ > 0.10) – suggesting NET shares structure with broad cortical gradients rather than one specific confound. To furthermore minimize the risk of physiological artifacts, we implemented a strict 36-parameter confound regression scheme during rsfMRI preprocessing^35^.

Across two independent datasets employing different measurements of peripheral autonomic activity, NET AUC+ scores were robustly associated with autonomic arousal measures across subjects and sessions (Fig. 4c; Tab. S14). First, NET AUC+ was significantly associated with in-scanner heart rate variability in the HCP-YA subsample (n = 64, 103 observations; Fig. S16). Second, it was strongly associated with low-frequency pupil diameter fluctuations in the Yale Resting State fMRI/pupillometry sample^36^ (YRSP; n = 25, 49 observations; Tab. S2; Fig. S17). These associations were robust to regressing out the covariates most strongly attenuating the group-level NET effect (Tab. S13), arguing against a shared-gradient confound (Tab. S14). Both analyses indicate an inverse relationship between NET AUC+ and autonomic arousal, with higher arousal linked to lower synchronization between regions with high NET availability, an association possibly related to the reported interruption of functional networks following tonic LC activation^37^.

### Neurobiological rsFC organization is sensitive to pharmacological modulation

As demonstrated above, the NEOFC framework is sensitive to neurobiological properties that underlie brain functional network organization. NEOFC metrics should therefore be altered by interventions affecting the same neurobiological systems. We tested this hypothesis in three healthy adult datasets (Tab. S2) applying four psychotropic substances – methylphenidate (MPH), risperidone, ketamine, and midazolam – in placebo-controlled, within-subject, crossover trials^24,38,39^. We systematically evaluated drug effects for both AUC+ and AUC–scores calculated on subject-/session-level across all reference atlases in separate mixed linear models (controlled for mean motion and global connectivity).

A combined overview of the results is shown in Fig. 5 (Tab. S15). First, MPH is expected to affect noradrenaline and dopamine transporters. We could not confirm effects of MPH on NET or DAT AUC in the available dataset, but we observed negative associations with AUC+ scores for an acetylcholine transporter and opioid receptor (VAChT and KOR, p_Meff_ < 0.05), indicating reduced rsFC between high-density brain regions under MPH influence. Risperidone showed nominally significant reductions of AUC–scores for the 5-HT_2A_ and D_2/3_ receptors compared to placebo – while the effects were well in line with risperidone’s known dopaminergic and serotonergic receptor targets, neither effect survived correction. For ketamine, we observed widely distributed effects on AUC+ and AUC–. Ketamine’s main target, the NMDA receptor, showed a nominal reduction of AUC–scores, but the strongest effects, surviving correction, were reductions of 5-HTT and NET AUC+. Lastly, exposure to midazolam resulted in a nominal increase in AUC– of the expected target receptor, GABA_A_, as the only such effect. We then tested for potentially opposing effects of ketamine and midazolam in a 3-factor model. The strongest effects following a “midazolam > placebo > ketamine” pattern were observed for NET AUC+, GABA_A_ AUC–, and serotonergic AUC–scores.

**Fig. 5:**
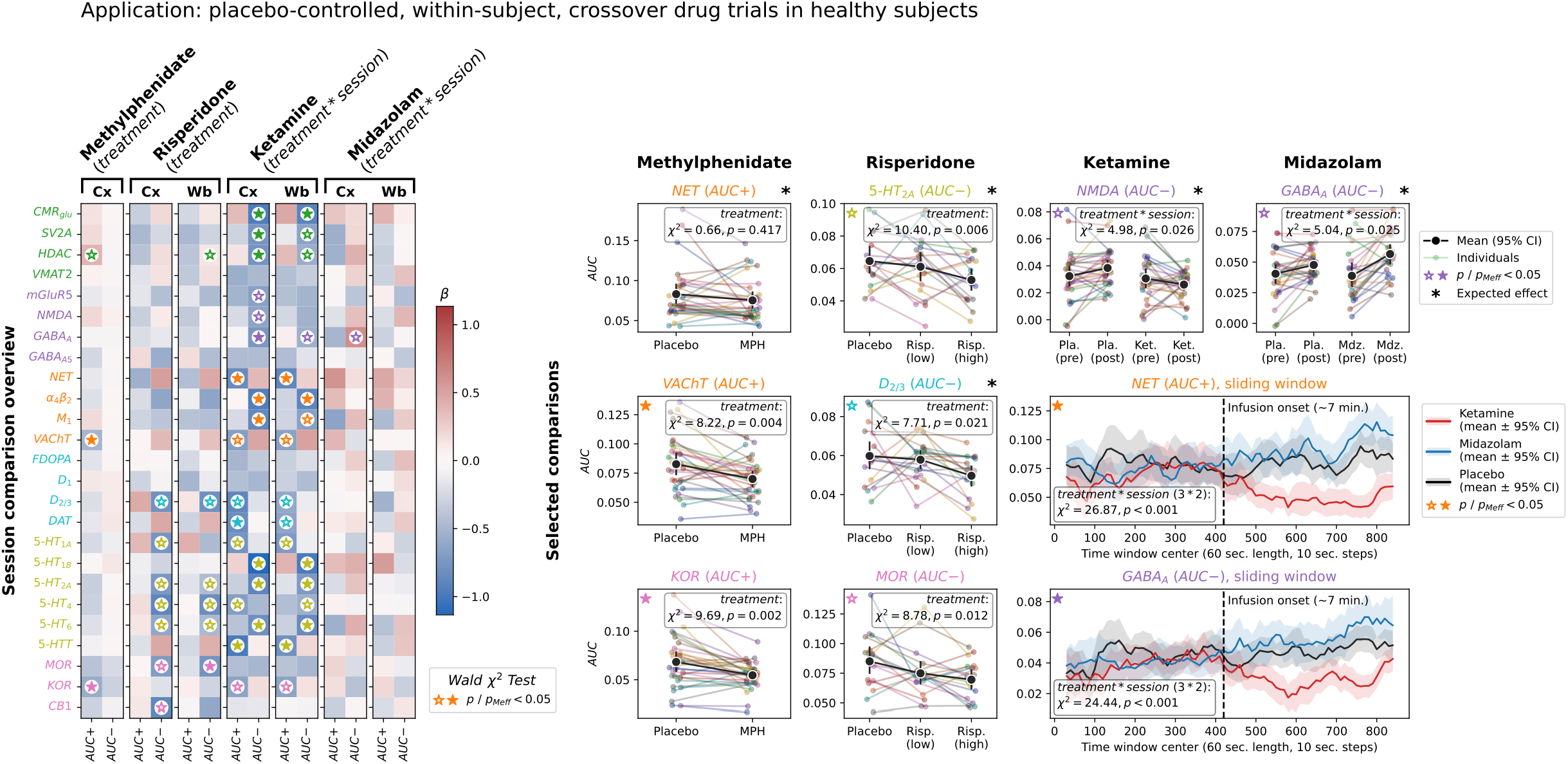
Neurobiological rsFC organization is modulated by pharmacological agents. Left, session comparison overview: LMM beta coefficients for treatment effects on AUC+ and AUC− across 25 nuclear imaging reference maps and 4 drug challenges for Schaefer200 and Schaefer200 + subcortex parcellations (methylphenidate and risperidone: treatment effect; ketamine and midazolam: treatment × session interaction). Color: standardized beta coefficient; significance markers indicate Wald χ² test p values. Right, selected comparisons, all for Schaefer200: small panels: individual AUC values per session for expected and strongest target effects (expected targets marked with an asterisk; methylphenidate: NET, VAChT, KOR; risperidone: 5-HT_2A_, D_2/3_, MOR; ketamine: NMDA; midazolam: GABA_A_). Colored dots with thin lines: individual subjects; Black dots with error bars: group mean with 95% bootstrapped CI. Lower right big panels: sliding window timecourses of mean AUC+ (NET) and AUC− (GABA_A_) across the infusion session (60-second windows, 10-second steps); ketamine: red; midazolam: blue; placebo: black; shaded area: 95% bootstrapped CI. Abbreviations: LMM: linear mixed model; AUC: area under the curve; FC: functional connectivity; CI: confidence interval; pMeff: p-value corrected for effective number of comparisons; see Fig. 2 for reference map abbreviations.

As ketamine and midazolam were applied intravenously during fMRI scanning, the data allowed us to further explore the acute pharmacological modulation of NEOFC by running a sliding-window analysis. For both drugs, we found a strong deviation of time-resolved AUCs from both placebo and pre-infusion data (Fig. 5, lower right), confirming the modulatory drug effects and the ability of the method to track rsFC patterns dynamically over time. Neurobiological rsFC organization is sensitive to network alterations in early psychosis

The pharmacological analyses indicate that the NEOFC framework carries potential for studying pathophysiological processes involved in psychiatric and neurological disorders. We explored this in a clinical experiment in the HCP Early Psychosis dataset (HCP-EP; n = 96 patients, n = 55 controls; Tab. S2). We systematically tested for group differences in both AUC+ and AUC– between the full patient and control cohorts, as well as after stratifying the psychosis group based on medication status (current and lifetime exposure; group differences in covariates: Tab. S16). Finally, we calculated partial Spearman associations of AUC indices with symptom and general cognitive function scales as well as medication intake in the psychosis group.

We found a strong reduction of primarily AUC+ scores in psychosis across various neurotransmitter systems (Fig. 6, left; Tab. S17). Serotonergic, followed by dopaminergic indices showed the strongest significant reductions (Fig. 6, center). These effects indicate reduced rsFC between serotonin and dopamine receptor-rich brain areas in psychosis. Notably, the serotonergic 5-HT_6_ AUC+ correlated negatively with current antipsychotics intake (Fig. 6, right). For the dopamine transporter, we found significant AUC–reduction and a nominal AUC+ reduction that was associated with higher symptom scores (Fig. 6, right). More generally, correlation patterns with clinical scales and medication intake were strongest with serotonergic and dopaminergic AUC+ indices, however, none survived correction for multiple comparisons (Fig. S18; Tab. S18). Medicated and unmedicated psychosis subgroups did not differ. The effects were largely similar when subcortical parcels were included (Fig. 6; Tab. S17).

**Fig. 6:**
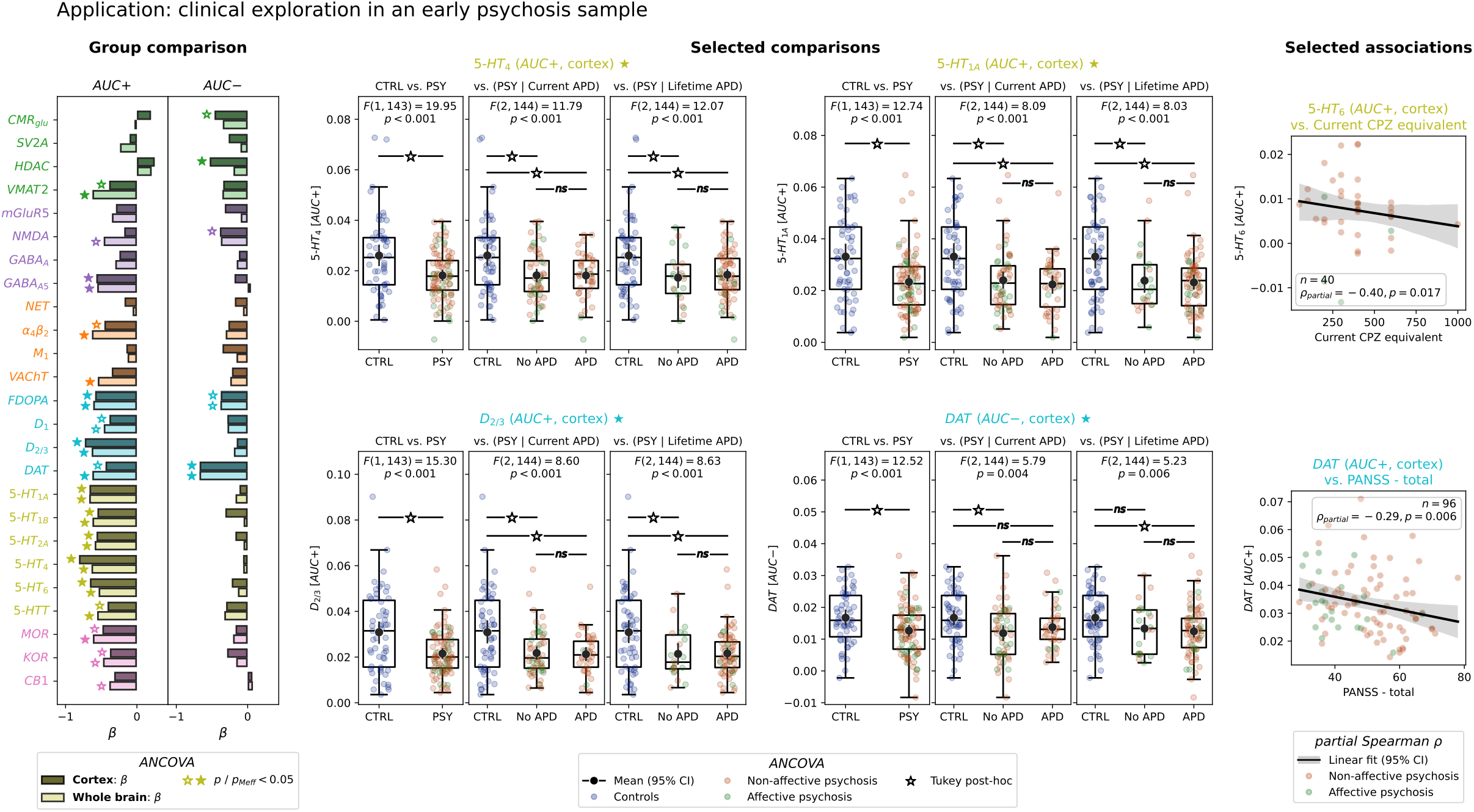
Neurobiological rsFC organization is altered in early psychosis. Left, group comparison: type II ANCOVA beta coefficients for CTRL vs. PSY group differences in AUC+ and AUC− across 25 nuclear imaging reference maps (site-harmonized data; covariates: age, sex, mean FD, global connectivity, current antipsychotic dose). Darker bars: Schaefer200; brighter bars: Schaefer200 + subcortex; significance markers indicate pMeff. Middle, selected comparisons: individual AUC values per diagnostic group for the 4 strongest effects (5-HT_4_, 5-HT_1A_, D_2/3_: AUC+; DAT: AUC−). Each panel shows three diagnostic contrasts: CTRL vs. PSY, CTRL vs. PSY stratified by current antipsychotic dose, and CTRL vs. PSY stratified by lifetime antipsychotic use. Subjects are additionally color-coded by controls (blue), and non-affective (orange) vs. affective psychosis (green). Dots: individual subjects; boxplots: median, quartiles and 1.5*interquartile range; black dots with errorbars: mean and 95% bootstrapped CI; brackets with significance markers: Tukey HSD post-hoc comparisons. Right, selected associations: Correlations between AUC+ and clinical variables within the PSY group (scatter: individual subjects; line: regression fit with 95% bootstrapped CI; partial Spearman’s rho reported). Abbreviations: AUC: area under the curve; FC: functional connectivity; ANCOVA: analysis of covariance; CTRL: healthy controls; PSY: psychosis; FD: framewise displacement; pMeff: p-value corrected for effective number of comparisons; see Fig. 2 for reference map abbreviations.

## Discussion

Here we examined how functional synchronization of the human brain relates to underlying neurobiology using an integrative framework based on brain network topology. We showed that functional connectivity measured with both haemodynamic and electrophysiological imaging modalities is robustly associated with neurotransmitter systems, with stronger connectivity observed within networks characterized by high or low expression of specific receptors or transporters. These effects were highly reliable and reproducible at group and individual levels. The close association between noradrenergic AUC scores and measures of physiological arousal provides external validation of the NEOFC framework in addition to evidence for the neurobiological relevance of the observed effects. The AUC scores were modulated by pharmacological interventions and altered in patients with psychotic symptoms, highlighting their potential for clinical research, which we facilitate by providing the underlying method as a Python package (mapconn).

We first validated the NEOFC approach using established resting-state networks as a positive control. For eight of the twelve networks, we observed the expected synchronization effects, indicating high sensitivity of the method to interregional rsFC organization. When applied to a comprehensive database of neurobiological atlases, most of these showed topological associations in terms of increased rsFC between regions with high or low density of the respective receptors or transporters. The strongest effects were consistently observed across all evaluated subjects and cohorts in both rsfMRI and MEG data, pointing to a general neurobiological basis. Individual AUCs, as well as group-level and single-subject AUC profiles across reference atlases, displayed good to excellent test-retest reliability.

The strongest evidence for a neurobiological source of the observed AUC effects comes from MEG, where the beta band showed the strongest correspondence to rsfMRI results, with an emphasis on VAChT (p_Meff_ < 0.05) and NET (p < 0.05) AUC+ effects. An orthogonalized (leakage-corrected) sensitivity analysis further supported a genuine, non-leakage-driven origin for the NET effect, whereas the VAChT effect was fully attenuated. As a more direct measure of neural activity, MEG is less likely to be biased by physiological confounds that are typically associated with rsfMRI, i.e., peripheral physiological signals. With the exception of GABA_A_, only AUC+ associations passed stringent significance thresholds, extending the recently reported neurotransmitter-related modulation of local intracranial EEG spectra to rsFC^40^. In line with the known association between gamma oscillation power and aggregated neural firing rates^41,42^, we observed the strongest MEG-neurobiology associations in gamma frequency bands. While gamma-band activity is typically generated locally, long-range network organization is enabled via cross-frequency coupling^43^, whereby fast local gamma activity is modulated by slower rhythms^41,44^.

With rsfMRI, we observe only a few AUC+ associations. The most consistent effects were found for noradrenaline, followed by acetylcholine transporter availability. The noradrenaline effects were primarily driven by a sensorimotor-insular network, overlapping with known interoceptive and sensory-integrative brain networks^45^. Two independent datasets provide convergent evidence for an association between these noradrenaline effects and arousal, indexed by heart rate variability and pupil unrest, and robust to regression of cortical gradients that attenuated the group-level NET effect. The tonic noradrenaline release by the LC is thought to interrupt the activity of functional networks to promote rapid behavioral adaptation^37^. This process reconfigures network activity from a tightly coupled, low-arousal state toward cross-network integration^37,46^. Consistent with this mechanism of action, we find higher arousal to be associated with lower individual AUC+ scores. More generally, a meta-analysis across all rsfMRI cohorts revealed a tendency of all neurotransmitter transporters towards AUC+ effects. Transporters are mainly presynaptic and closely tied to long-range ascending projection systems, so their network signatures may be sufficiently strong and spatially coherent to be detected in hemodynamic rsFC^33,47^.

The majority of rsfMRI effects displayed increased rsFC between regions with low density of the neurobiological atlases as indicated by AUC–effects. Notably, these effects were largely absent in MEG data. Such associations may be expected for receptors or transporters carrying primarily local information, i.e. if a receptor’s activity rendered the local BOLD signal more unique, its absence could facilitate synchronization. A more integrative explanation may lie in frequency-specific modes of interregional communication^48^. Regions interact through neurotransmission often at higher frequencies, such that synchronization is found in fast dynamics and therefore captured by MEG (gamma)^41–43^. In contrast, the same neural sources are observed in rsfMRI through neurovascular coupling and after heavy temporal filtering, which can attenuate interregional information and leave a more local-appearing metabolic covariance pattern^7,49^. Only ultra-slow fluctuations in neural population activity, producing large-scale networks with high spatial coherence, are captured directly by rsfMRI, though faster dynamics may be reflected by neurovascular coupling. Such slow signals may be tracked especially by MEG beta band amplitude envelopes. The beta band showed the strongest rsfMRI-MEG cross-modal convergence in our study and is generally considered spatiotemporally robust with a favorable signal-to-noise ratio^3,50,51,30^. Strongly supporting this reasoning, Sastre-Yagüe et al.^52^ recently demonstrated that causal manipulation of cortical excitability in mice dissociably affected high frequency electrophysiological coherence versus ultralow frequency rsfMRI connectivity. In line with this notion of opposing effects, we find a negative association between AUC– profiles observed with rsfMRI and AUC+ profiles observed with MEG and vice versa. Despite clear dissociations in the individual significant AUC effects, significant cross-modal correlations between AUC+ and AUC– profiles support the notion that both imaging modalities actually capture information from high and ultra-slow frequency spectra, though clearly with preferential sensitivity.

Using pharmacological challenge data, we tested whether the AUC scores are modifiable by targeted interventions. Consistent with their mechanisms of action, we found that risperidone, ketamine, and midazolam modulated the AUC of receptors with high affinity of the respective compounds^53–56^. Risperidone nominally reduced AUC–estimates of multiple serotonergic and the dopamine D_2/3_ receptors (uncorrected p < 0.05), which may be expected from its receptor antagonism, with downregulation reducing the desynchronization. Ketamine, an NMDA receptor antagonist, is known for its widespread direct and downstream effects on a multitude of neurotransmitter signaling pathways^55,57,58^. Consistently, we found non-specific ketamine effects on AUC across most of the evaluated neurotransmitter systems. AUC effects of ketamine and midazolam displayed opposing patterns for GABA_A_ AUC– and NET AUC+. While ketamine generally increases sympathetic drive, a property also linked to its antidepressant effects^59^, midazolam generally shows inhibiting effects. Despite its GABA-mediated inhibitory properties and in contrast to the other compounds, midazolam generally increased rsFC, supporting the notion that deactivation rather than activation of a receptor may often support long-range rsfMRI network synchronization. Contrary to its mechanism of action as a noradrenaline and dopamine reuptake inhibitor, MPH showed AUC+ reductions only for a cholinergic transporter and the κ-opioid receptor. The limited rsfMRI sensitivity to its primary mechanism of action is consistent with previous studies reporting similar negative MPH findings for local activity measures^10^. While further investigation is necessary, possible explanations include its indirect mechanism of action as a reuptake inhibitor, direct or downstream effects of MPH on the opioid system^60^, and limited effects of MPH in healthy adults^61^.

AUC profiles in patients with psychotic symptoms differed from controls across most neurobiological systems, with the strongest alterations seen for dopaminergic and serotonergic neurotransmission, consistent with their established role in psychosis^62,63^. All of the observed effects indicate AUC reductions, with stronger effects for AUC+. This pattern, which was independent of antipsychotic medication and global connectivity in our early psychosis sample, suggests a general disruption of the typical coupling between neurobiology and large-scale rsFC organization in psychosis. Notably, tested in control subjects against the spatial null, the respective AUC+ scores were significant only in MEG but not in rsfMRI, which raises questions regarding the directionality of this disruption in psychosis and the relevance of high-frequency spectra to the observed findings. Future corroboration is needed, ideally with PET and electrophysiological imaging.

While all of these findings provide compelling evidence for the neurobiological relevance of NEOFC-derived measures, key limitations of the framework include (i) absence of individual neurobiological data naturally preventing concrete statements about individual molecule availability, (ii) correlative nature of the non-interventional findings limiting causal interpretation, (iii) dependence on the quality and representativeness of nuclear imaging reference maps, (iv) reduced specificity due to intercorrelated maps, and (v) current restriction to univariate analyses^64–66^. Validation was further limited by heart rate estimation based on pulse-oximetry (relative to electrocardiography) in HCP-YA and small, demographically narrow pharmacological cohorts applying drugs with partly broad binding profiles. For atlases without independent validation comparable to NET’s, pharmacological and clinical AUC differences should be interpreted as consistent with, rather than proof of, reference-specific biology. Future work should focus on multivariate modeling across reference atlases, systematic comparison of connectivity estimators and metrics, integration with simultaneous fMRI-EEG and fMRI-PET, and clinical out-of-sample prediction. The framework is readily extensible to other biological priors, electrophysiological connectomes (e.g., EEG), task-based or dynamic rsFC, and within-subject mapping (e.g., structure–function coupling).

## Method

### Data sources and general processing

#### Ethics statement

No new data were acquired for this study. Ethical approval for the use of publicly available and restricted-access datasets including human demographic, behavioral, and neuroimaging data has been granted by the Heinrich-Heine-University, Düsseldorf, Germany. For information on ethical approvals for each dataset as provided by the local committees, we refer to the original sources cited in the text. Informed consent was obtained from each participant and their parents in the case of underage subjects (the latter applying only to a small subset of the HCP Early Psychosis sample).

#### Software

For connectomes which were not obtained directly from other publications, preprocessing of rsfMRI data was performed using the HCP pipeline^67^ for the HCP-EP sample and with FreeSurfer^68^ (7.4.1) and fMRIprep^69^ (24.0.1) for the remaining datasets. Postprocessing and extraction of connectomes was done with XCP-D^70^. All further analyses were performed in a Python (3.10.15) environment, relying on the NiSpace^25^ (0.0.1-beta.4) toolbox and mapconn^71^ (0.0.1) packages, as well as on routines from numpy^72^ (1.26.4), pandas^73^ (2.2.3), scipy^74^ (1.12.0), statsmodels^75^ (0.14.4), pingouin^76^ (0.5.5), neuroHarmonize^77^ (2.4.5), Nilearn^78^ (0.10.4), neuromaps^26^ (0.0.5), NeuroKit2^79^ (0.2.11), matplotlib^80^ (3.9.2), and seaborn^81^ (0.13.2).

#### Reference atlases

All deployed reference atlases, i.e., nuclear imaging, resting-state network probability, and covariate maps, were obtained directly through NiSpace in parcellated, tabular format. Originally, these data stem from independent studies conducted in healthy adults, which then shared their group-level outputs for public use. Sources of the nuclear imaging reference atlases are listed in Table S1^10,11,17,26,82–115^. A known problem with the currently available collections of nuclear imaging atlases is the heterogeneity of spatial registration target spaces, which are often unknown. To approach this, we registered all group-average maps to 2 mm MNI152NLin6Asym space using a non-linear deep-learning based method (SynthMorph v4)^116^. The resulting maps were masked with a liberal grey matter mask generated from the Harvard-Oxford atlas and scaled from 1e-6 to 1. Resting-state network probability maps, i.e., the probability that a given voxel belongs to a given canonical resting-state network across multiple subjects, were obtained from Dworetsky et al.^28^ and registered to MNI152NLin6Asym space as described above. Atlases used as spatial covariates were provided by several separate publications: T1/T2^29^, sensory-association axis^117^, cytoarchitectural gradient maps derived from post-mortem human histology^118,119^, as well as probability maps of gray matter and cerebrospinal fluid^120^, cerebral veins^121^, and arteries^122^.

#### Brain parcellations

We employed the Schaefer parcellation^123^ at 200-parcel density with and without 16 FreeSurfer subcortical parcels (“aseg”, excluding the brainstem) as the main parcellation scheme. To evaluate effects of different parcellation densities, we additionally tested 100- and 400-parcel densities with and without subcortical parcels only in the rsfMRI discovery and retest, MEG replication, and rsfMRI-MEG association analyses. The parcellations were applied directly in the target image spaces, i.e. in MNI152NLin6Asym for volumetric reference data, fsaverage for reference data originally provided in surface format, and fsLR for functional data.

#### Study samples and subject exclusions

Full sample descriptions and demographic details are provided in Supplementary Methods and Table S2. All rsfMRI sessions with mean framewise displacement (FD) > 0.3 mm were excluded (except the MPH dataset, for which preprocessed connectomes were obtained directly from the original authors). Briefly, the primary discovery sample was drawn from the HCP-YA dataset^29^ (n = 112, two resting-state runs; MEG data available for a subset of n = 33). The YRSP dataset^36,124^ provided simultaneous rsfMRI and pupillometry recordings from n = 25 subjects (49 sessions) for the physiological association analyses. Three placebo-controlled, randomized, within-subject crossover studies were included for pharmacological analyses: an oral MPH study^24^ (placebo vs. MPH; n = 30), an oral risperidone study^38^ (placebo vs. low vs. high dose; n = 17–19 sessions per condition), and an intravenous ketamine/midazolam study^39^ with pre- and post-infusion rsfMRI (placebo vs. ketamine vs. midazolam; n = 24–28 sessions per condition). The HCP-EP dataset^125^ served as the clinical application case, comprising 151 subjects across 4 acquisition sites (n = 96 early psychosis, n = 55 controls).

#### Resting-state fMRI processing

Full preprocessing details are provided in the Supplementary Methods. Briefly. the HCP-YA and HCP-EP datasets were obtained in “minimally preprocessed” format as generated by the HCP-pipeline^67,126^; all remaining datasets (except MPH, for which connectomes were provided directly^24^) were preprocessed using FreeSurfer and fMRIprep. All data was registered to the fsLR surface space (91k density). Except for MPH, postprocessing and connectome extraction was performed with XCP-D, applying a 36-parameter nuisance regression strategy with despiking and bandpass filtering (0.01–0.08 Hz)^35^. Pearson FC matrices were computed between parcellated time-series and Fisher z-transformed. For HCP-YA and HCP-EP, connectomes were averaged across phase-encoding directions within each run.

#### MEG processing

Source-reconstructed and parcellated resting-state MEG connectomes were provided by Shafiei et al.^30^; the processing was described in detail in the cited paper. Preprocessing and connectome extraction followed Brainstorm^127^ recommendations and included cleaning of the sensor-level data, reconstruction of source-level data on the individual fsLR surfaces, parcellation of the vertex-level time series, and connectome calculation using amplitude envelope correlation^128^ at 6 canonical electrophysiological bands (delta: 2–4 Hz, theta: 5–7 Hz, alpha: 8–12 Hz, beta: 15–29 Hz, low gamma: 30–59 Hz, and high gamma: 60–90 Hz). As we were interested in the shared signal across regions, we did not apply pairwise signal orthogonalization in the primary analysis, preserving the original network topology. We test for statistical significance using spatial null models of the reference atlases while the individual input connectomes remain unchanged during the null runs. Because the dominant component of MEG source leakage is a smooth function of inter-regional distance, this component is controlled by construction: any resulting spurious signal affects observed and null runs equally, providing a control for the associated false positives. This approach does not, however, control for spatially inhomogeneous leakage arising from regional differences in source-reconstruction accuracy, which the spin test cannot capture. To address this remaining source of bias, we additionally computed AEC connectomes after pairwise orthogonalization^3^ (Brainstorm default settings, see Shafiei et al.^30^), repeated all main analyses and compared non-orthogonalized and orthogonalized results with Spearman correlations across AUC profiles. Because orthogonalization removes all shared zero-lag signals, it may attenuate genuine near-zero-lag neuronal interactions alongside leakage^129–131^, and its results should be interpreted as a conservative bound rather than a leakage-free ground truth.

### Linking reference atlases to connectomes

#### Implementation

To derive the association between a parcellated atlas and an individual connectome (i.e., a region-to-region FC matrix), we (i) converted the reference atlas to percentiles of its own distribution, (ii) masked the connectome to keep only connections between regions that exceeded a specified reference atlas percentile, (iii) systematically repeated this for all percentiles in {0, 5, 10, …, 95}, and (iv) calculated the average FC of the masked connectome for each percentile, resulting in (v) a curve showing average FC as a function of percentile threshold. Global connectivity (the value at the 0th percentile, i.e., average FC across all connections) was subtracted from each curve to account for interindividual, acquisition- and processing-related differences in global connectivity. The area under the curve (AUC), computed via the trapezoidal rule, served as the prime aggregative index of the map–connectome association per individual. The inverse test (i.e., producing AUC–scores) was implemented by inverting each original reference atlas map around its mean, such that the highest-ranked regions became the lowest-ranked, and recomputing the steps lined out above.

We conducted several sensitivity analyses reported at appropriate places in the main text. In the initial rsfMRI analyses, we evaluated the leading coefficient of a second-degree polynomial fitted to the FC-percentile curve as an alternative aggregate metric, focussing on the shape of the curve rather than on the overall effect beyond the global connectivity. In the rsfMRI discovery analyses, to additionally control for spatial autocorrelation, we calculated the delta between aggregate outcomes from the original and inverted tests for further evaluation only. In a sensitivity analysis to assess if observed effects on rsFC were driven by the highest-density atlas regions, we systematically varied the maximum percentile threshold in AUC calculation from 5 to 90 and reported the lowest threshold above which significance emerged continuously. To test if the NET AUC+ effect was driven by possibly confounding spatial factors, we individually regressed each of a set of covariate maps from the NET atlas and reran the AUC+ estimation on the residuals.

#### Null models

Spatial patterns that are shared across brain-wide signals, but convey no information of interest, can inflate associations measures between the same signals^27^. Therefore, we evaluated each association against null models of the reference atlases, retaining spatial autocorrelation and interhemispheric symmetry. Null models were generated independently for parcellated cortical and subcortical data before conversion to percentiles. For cortical data, we employed a symmetric “spin test” on the fsLR spherical projection with nearest-neighbor reassignment between original and rotated coordinates^26,132^. As this process is only applicable to cortical data, null models for subcortical data were generated independently using Moran Spectral Randomization^133,134^ based on a Euclidean distance matrix. Interhemispheric symmetry was ensured through centerline-mirrored rotations in the first, and a near-symmetric distance matrix in the second case. We generated 1,000 null maps for each input reference map and recalculated NEOFC metrics with each. Exact *p* values were determined based on the resulting null distribution on individual as well as on group levels (group level: sample mean of observed scores vs. sample mean of null scores).

To compare the derived scores akin to effect sizes while respecting non-normal null distributions, we calculated robust z-scores by subtracting the median of the null distribution from the observed statistic and dividing by the median absolute deviation scaled by 1.4826 (the consistency factor for a normal distribution). These z-scores were used to present NEOFC metrics across multiple reference atlases (i.e., Fig. 2a/b and 3b) and in all analyses that compared reference atlas profiles between cohorts or modalities (e.g., Fig. 3). This procedure minimized the influence of unwanted spatial influences on profile-level associations in which, e.g., different levels of autocorrelation across atlases could drive the results. Supplementary tables contain both original and zscored values.

#### Multiple comparison correction

We empirically adjusted for alpha-error accumulation within sets of tests within each analysis by calculating the effective number of tests (M_eff_) based on eigenvalues of the Pearson correlation matrix of the input data to each analysis^135^. We then used the estimated factor to correct p values with Šidák’s correction as: p_Meff_ = 1 – (1 – p)^Meff^. This approach was applied to each analysis as described in the respective sections.

#### Positive-control and discovery analysis

The NEOFC framework was first applied to the 12 resting-state network probability maps as a positive control, and subsequently to the full nuclear imaging atlas database as the primary discovery analysis, both in the HCP-YA rsfMRI sample and separately per run. For each analysis, group- and individual-level NEOFC scores and associated p values were derived from the null distributions as described above. To correct for the effective number of simultaneous tests, the M_eff_ was computed from the Pearson intercorrelation matrix of the respective atlas set (resting-state networks or nuclear imaging) separately for each tested parcellation and applied to group-level p values for both AUC+ and AUC−. Individual-level significance was assessed at uncorrected p < 0.05; the proportion of subjects exceeding this threshold was reported to characterize the consistency of effects across individuals. We reported results for the first rsfMRI run in the results section (all data in supplementary tables).

#### Test-retest analysis

Test-retest reliability of individual-level NEOFC scores was evaluated using the two available rsfMRI runs from the HCP-YA dataset, acquired approximately one day apart. For each reference atlas and each combination of parcellation, NEOFC metric, and testing direction (e.g., AUC+ vs. AUC–), the intraclass correlation coefficient ICC(2,k) (two-way random effects model, average measures, absolute agreement) was calculated using pingouin. As a complementary measure of within-subject variability, the within-subject coefficient of variation (WCV) was computed as the root mean square of individual coefficients of variation across runs. Statistical significance of the WCV was assessed against a permutation null distribution (n = 1,000) generated by randomly exchanging session labels within subjects.

#### Reproducibility analysis and p value meta-analysis

Reproducibility of the group mean AUC profiles across independent samples was assessed in two complementary ways; AUC profiles of HCP-YA and YRSP samples were averaged across runs before analyses. First, pairwise Spearman correlations between the group-mean profiles (robust z-scores) across the nuclear imaging reference atlases were computed for all dataset pairs. Profile reproducibility was additionally summarized as an intraclass correlation coefficient ICC(3,1), treating reference atlases as targets and datasets as raters, separately for AUC+ and AUC−. Second, to increase statistical power to detect weaker effects, a meta-analysis was performed across the six independent fMRI datasets using Stouffer’s method^136^, with sample sizes as weights. The M_eff_ for multiple comparison correction was calculated from the reference atlases as described above and applied to the combined p values.

### fMRI-MEG associations

#### Cross-modal associations across NEOFC profiles

Cross-modal profile associations were assessed in the subset of subjects with both quality-controlled rsfMRI and MEG data available (n = 29; rsfMRI data averaged across runs). We calculated Spearman correlations between rsfMRI and MEG AUC profiles (robust z-scores) across all nuclear imaging reference atlases on two levels: individually for each of the 29 subjects, and on the group-level for the mean AUC (rsfMRI averaged across runs).

#### Cross-modal associations per reference atlas across subjects

Cross-modal associations per reference atlas were assessed by computing, for each nuclear imaging reference atlas and each MEG frequency band separately, the Spearman correlation between individual rsfMRI (run-averaged) and MEG AUC scores across subjects.

### Brain-regional influence estimation

To identify brain regions and connections most influential to the aggregate NEOFC scores, we performed two complementary leave-one-out analyses. We focused on the main whole-brain parcellation (Schaefer-200 with subcortical parcels) to enable simultaneous inference on both structures. First, in a leave-one-region-out analysis, the AUC was recomputed after excluding all connections involving a given region. Second, in a leave-one-connection-out analysis, individual connections were excluded for a given region. In both cases, the importance score was defined as the difference between the full AUC and the leave-one-out AUC, averaged across subjects. Analyses were conducted separately for AUC+ and AUC−. In supplementary figures, we additionally show the spatial correlation pattern between leave-one-region-out maps and the original atlases after conversion to percentiles.

### Physiological associations

Details on physiological data processing are provided in the Supplementary Methods. Pupil area time series (YRSP, pre-downsampled to 1 Hz) were provided minimally preprocessed and further cleaned using amplitude and velocity filtering^137^. Multiple arousal indices were derived: the pupil unrest index (PUI; root mean squared of first-differences)^138,139^ and power in different frequency bands defined by Rizzuto et al.^140^. Photoplethysmography signals (HCP-YA, 400 Hz) were processed with NeuroKit2 (Elgendi method) to extract RMSSD^141^ and MadNN, averaged across phase-encoding directions within runs. Sessions were excluded if the NeuroKit2 quality index fell below a data-driven threshold or HRV exceeded 150 ms.

Associations between NET AUC+ and physiological measures were tested using linear mixed models (LMMs) with a random intercept per subject to account for repeated observations across runs. Both the NEOFC scores and the physiological variable were z-standardized prior to modeling. For HCP-YA, covariates included mean FD, sex, age, BMI, and global connectivity; for YRSP, mean FD and global connectivity (demographics not available). Models were fitted via maximum likelihood using the Powell optimizer separately for the two physiological outcomes per dataset and per parcellation. Prior analyses showed that the NET AUC+ effect may share variance with broad cortical gradients. To ensure that physiological associations arose from a NET-specific pattern, we repeated the LMMs using NET AUC+ scores calculated after regressing out covariate maps from the NET map (covariate maps: sensory-association axis, functional gradient, microstructural gradient).

### Pharmacological challenges

Drug effects on NEOFC scores were assessed using LMMs with a random intercept per subject, fitted via maximum likelihood with the Powell optimizer separately for each reference atlas, AUC+ and AUC–scores, and parcellation. Treatment effects were evaluated using Wald Chi² tests. Mean FD and global connectivity were generally included as covariates to account for potential drug-induced differences in motion and general FC strengths. Multiple testing correction was applied across reference atlases per analysis set, with M_eff_ estimated from the Pearson intercorrelation matrix of individual-level AUC scores within each drug sample. All NEOFC scores and continuous covariates were z-standardized prior to modeling.

For MPH (2-level: placebo/MPH) and risperidone (3-level: placebo/low dose/high dose), we evaluated the main effect of treatment. For MPH, only cortical data and no FD information was available. For ketamine and midazolam, each analyzed separately in a 2×2 within-subjects factorial design (treatment: placebo/drug × session: pre/post-infusion), we focused on the treatment × session interaction. A joint 3×2 model (placebo/ketamine/midazolam × session) was additionally fitted to directly compare the two compounds.

### Clinical comparison

Prior to group comparisons, AUC scores were harmonized across the 4 acquisition sites using ComBat as implemented in neuroHarmonize^77,142,143^. Harmonization was performed separately for each parcellation and testing direction, retaining sex, age, mean FD, and global connectivity as biological covariates. The harmonization model was learned on controls and subsequently applied to patients.

Group differences in NEOFC scores between patients and controls were tested using type II analyses of covariance with Tukey HSD post hoc tests separately per reference atlas, testing direction, and parcellation. All NEOFC scores and continuous covariates were z-standardized prior to modeling. Covariates included sex, age, mean FD, global connectivity, and current antipsychotic drug intake in chlorpromazine equivalents. The M_eff_ was estimated from the intercorrelation of individual-level AUC scores across reference atlases within each analysis set. Three sub-analyses were conducted: (i) restricting the patient group to currently unmedicated subjects vs. controls (no drug intake covariate); (ii) comparing controls, currently medicated, and currently unmedicated patients; and (iii) comparing controls, lifetime medicated, and lifetime unmedicated patients.

Association analyses were calculated in the psychosis group only and between NEOFC scores and six clinical and cognitive outcomes: PANSS total, positive, and negative subscales, current antipsychotics dose and duration of exposure, and the NIH cognitive composite score. For non-antipsychotics outcomes, Spearman correlation and partial Spearman correlation were computed in parallel, adjusting for sex, age, mean FD, global connectivity, and current antipsychotics dose. Correlations with antipsychotics outcomes were calculated without the antipsychotics covariate and while excluding subjects with zero exposure. The M_eff_ was estimated accordingly as the product of the effective number of reference atlases and the effective number of outcomes.

## Supporting information

Supplementary Methods and Results

Supplementary Animation S1

## Data and code availability

All reference atlases used in this study are distributed as part of the NiSpace toolbox (https://github.com/leondlotter/nispace). HCP-YA resting-state fMRI data are available through the HCP data portal (https://db.humanconnectome.org). The HCP-EP data are available via the NDA data portal (https://nda.nih.gov/edit_collection.html?id=2914). Both are subject to data use agreements. Resting-state fMRI and pupillometry data from the YRSP dataset are publicly available via OpenNeuro (https://openneuro.org/datasets/ds003673/versions/2.0.1). The pharmacological datasets are available from the respective principal investigators upon reasonable request. Individual-level data cannot be shared publicly due to data governance restrictions of the source datasets.

The NEOFC framework is implemented in the open-source mapconn Python toolbox, available at https://github.com/leondlotter/mapconn (DOI: 10.5281/zenodo.19859027). All analysis code and sharable (including all group-level) data are available in the GitHub repository associated with this publication; https://github.com/leondlotter/neofc (DOI: 10.5281/zenodo.19858671). Note that full reproduction of all reported analyses requires access to restricted datasets and is therefore largely not directly possible from the shared code alone. All data can be shared upon proof of data access rights to the corresponding authors.

## Acknowledgements

We thank all study participants and scientists who agreed to share their data, making this research possible. Data were provided in part by the Human Connectome Project, WU-Minn Consortium (Principal Investigators: David Van Essen and Kamil Ugurbil; 1U54MH091657) funded by the 16 NIH Institutes and Centers that support the NIH Blueprint for Neuroscience Research; and by the McDonnell Center for Systems Neuroscience at Washington University. Research using Human Connectome Project for Early Psychosis (HCP-EP) data reported in this publication was supported by the National Institute of Mental Health of the National Institutes of Health under Award Number U01MH109977. The HCP-EP 1.1 Release data used in this report came from DOI: 10.15154/1522899.

LDL and AK were supported by the Federal Ministry of Education and Research (BMBF) and the Max Planck Society (MPG), Germany. LDL, SBE and JD received support from the German Research Foundation (DFG; DU 2027/7-1 project number 549186835). GS was supported by a postdoctoral fellowship from the Canadian Institutes of Health Research (CIHR). NH was supported by an Intermediate Clinical Fellowship from Wellcome. SM and AF were supported by F Hoffman La Roche Ltd. BM acknowledges support from the Natural Sciences and Engineering Research Council of Canada (RGPIN-2017-04265), Canadian Institutes of Health Research (PJT-180439), and Canada Research Chairs Program (CRC-2022-00169). SC acknowledges support through the European Union’s Horizon Europe research and innovation programme under grant agreement No 101147319 (EBRAINS 2.0 Project). JK acknowledges financial support for the Mapping Autonomic Neural Interaction and Control (MANIAC) Emerging Group by the University of Cologne Excellent Research Support Program. KRP and CP received support from the DFG (KRP: SSP 2205; CP: Emmy Noether Programme – 524408221).

## Author contributions

Conceptualization: JD, LDL; Data curation and Investigation: LDL, GS, DL, OD, MM, MC, AS, NH, SH, DU, IY, SM, AF, JH, BM, KP; Formal analysis: LDL, JD, AK; Funding acquisition: JD, LDL, SBE; Methodology, Project administration, Software, Validation: LDL, JD; Resources: JD, SBE; Supervision: JD, SBE, CP, JK, SC; Visualization: LDL; Writing – original draft: LDL, JD; Writing – review & editing: all authors.

## References

1. Biswal, B., Zerrin Yetkin, F., Haughton, V. M. & Hyde, J. S. Functional connectivity in the motor cortex of resting human brain using echo-planar mri. Magnetic Resonance in Medicine 34, 537–541 (1995).

2. Damoiseaux, J. S. et al. Consistent resting-state networks across healthy subjects. Proceedings of the National Academy of Sciences 103, 13848–13853 (2006).

3. Hipp, J. F., Hawellek, D. J., Corbetta, M., Siegel, M. & Engel, A. K. Large-scale cortical correlation structure of spontaneous oscillatory activity. Nat Neurosci 15, 884–890 (2012).

4. Huntenburg, J. M., Bazin, P.-L. & Margulies, D. S. Large-Scale Gradients in Human Cortical Organization. Trends in Cognitive Sciences 22, 21–31 (2018).

5. Taylor, H. P. et al. Functional hierarchy of the human neocortex across the lifespan. Nature 1–10 (2026) doi:10.1038/s41586-026-10219-x.

6. Biswal, B. B. & Uddin, L. Q. The history and future of resting-state functional magnetic resonance imaging. Nature 641, 1121–1131 (2025).

7. Logothetis, N. K. What we can do and what we cannot do with fMRI. Nature 453, 869–878 (2008).

8. Bazinet, V., Hansen, J. Y. & Misic, B. Towards a biologically annotated brain connectome. Nat. Rev. Neurosci. 24, 747–760 (2023).

9. Baillet, S. Magnetoencephalography for brain electrophysiology and imaging. Nat Neurosci 20, 327–339 (2017).

10. Dukart, J. et al. Cerebral blood flow predicts differential neurotransmitter activity. Sci Rep 8, 4074 (2018).

11. Dukart, J. et al. JuSpace: A tool for spatial correlation analyses of magnetic resonance imaging data with nuclear imaging derived neurotransmitter maps. Human Brain Mapping 42, 555–566 (2021).

12. Lotter, L. D., Nehls, S., Losse, E., Dukart, J. & Chechko, N. Temporal dissociation between local and global functional adaptations of the maternal brain to childbirth: A longitudinal assessment. 2023.08.15.553345 Preprint at 10.1101/2023.08.15.553345 (2023).

13. Kasper, J. et al. Resting state changes in aging and Parkinson’s disease are shaped by underlying neurotransmission – a normative modeling study. 2023.10.18.562677 Preprint at 10.1101/2023.10.18.562677 (2023).

14. Kasper, J. et al. Local synchronicity in dopamine-rich caudate nucleus influences Huntington’s disease motor phenotype. Brain 146, 3319–3330 (2023).

15. Hahn, L. et al. Resting-state alterations in behavioral variant frontotemporal dementia are related to the distribution of monoamine and GABA neurotransmitter systems. eLife 13, e86085 (2024).

16. Grumbach, P. et al. Local activity alterations in individuals with autism correlate with neurotransmitter properties and ketamine-induced brain changes. Nat Commun 16, 8248 (2025).

17. Hansen, J. Y. et al. Mapping neurotransmitter systems to the structural and functional organization of the human neocortex. Nat Neurosci 25, 1569–1581 (2022).

18. Hansen, J. Y. et al. Integrating multimodal and multiscale connectivity blueprints of the human cerebral cortex in health and disease. PLoS Biol 21, e3002314 (2023).

19. Saiz-Masvidal, C. et al. Mapping cross-modal functional connectivity of major neurotransmitter systems in the human brain. Brain Struct Funct 230, 137 (2025).

20. Siebenhühner, F., Palva, J. M. & Palva, S. Linking the microarchitecture of neurotransmitter systems to large-scale MEG resting state networks. iScience 27, (2024).

21. Shafiei, G. et al. Neurophysiological signatures of cortical micro-architecture. Nat Commun 14, 6000 (2023).

22. Dipasquale, O. et al. Receptor-Enriched Analysis of functional connectivity by targets (REACT): A novel, multimodal analytical approach informed by PET to study the pharmacodynamic response of the brain under MDMA. NeuroImage 195, 252–260 (2019).

23. Cao, Z. et al. Unraveling the molecular relevance of brain phenotypes: A comparative analysis of null models and test statistics. NeuroImage 293, 120622 (2024).

24. Dipasquale, O. et al. Unravelling the effects of methylphenidate on the dopaminergic and noradrenergic functional circuits. Neuropsychopharmacol. 45, 1482–1489 (2020).

25. Lotter, L. D. & Dukart, J. Nispace: Neuroimaging Spatial Colocalization Environment. Zenodo 10.5281/zenodo.12514622 (2024).

26. Markello, R. D. et al. neuromaps: structural and functional interpretation of brain maps. Nat Methods 1–8 (2022) doi:10.1038/s41592-022-01625-w.

27. Markello, R. D. & Misic, B. Comparing spatial null models for brain maps. NeuroImage 236, 118052 (2021).

28. Dworetsky, A. et al. Probabilistic mapping of human functional brain networks identifies regions of high group consensus. NeuroImage 237, 118164 (2021).

29. Van Essen, D. C. et al. The WU-Minn Human Connectome Project: An overview. NeuroImage 80, 62–79 (2013).

30. Shafiei, G., Baillet, S. & Misic, B. Human electromagnetic and haemodynamic networks systematically converge in unimodal cortex and diverge in transmodal cortex. PLOS Biology 20, e3001735 (2022).

31. Lotter, L. D. et al. Regional patterns of human cortex development correlate with underlying neurobiology. Nat Commun 15, 7987 (2024).

32. Schroeter, S. et al. Immunolocalization of the cocaine- and antidepressant-sensitive l-norepinephrine transporter. Journal of Comparative Neurology 420, 211–232 (2000).

33. Samuels, E. R. & Szabadi, E. Functional Neuroanatomy of the Noradrenergic Locus Coeruleus: Its Roles in the Regulation of Arousal and Autonomic Function Part I: Principles of Functional Organisation. Current Neuropharmacology 6, 235–253 (2008).

34. Bolt, T. et al. Autonomic physiological coupling of the global fMRI signal. Nat Neurosci 28, 1327–1335 (2025).

35. Ciric, R. et al. Benchmarking of participant-level confound regression strategies for the control of motion artifact in studies of functional connectivity. NeuroImage 154, 174–187 (2017).

36. Lee, K. et al. Arousal impacts distributed hubs modulating the integration of brain functional connectivity. NeuroImage 258, 119364 (2022).

37. Aston-Jones, G. & Cohen, J. D. An integrative theory of locus coeruleus-norepinephrine function: adaptive gain and optimal performance. Annu Rev Neurosci 28, 403–450 (2005).

38. Hawkins, P. C. T. et al. The effect of risperidone on reward-related brain activity is robust to drug-induced vascular changes. Human Brain Mapping 42, 2766–2777 (2021).

39. Forsyth, A. et al. Comparison of local spectral modulation, and temporal correlation, of simultaneously recorded EEG/fMRI signals during ketamine and midazolam sedation. Psychopharmacology 235, 3479–3493 (2018).

40. Stoof, U. M., Friston, K. J., Tisdall, M., Cooray, G. K. & Rosch, R. E. Topographic Variation in Human Neurotransmitter Receptor Densities Explains Differences in Intracranial EEG Spectra. Hum Brain Mapp 46, e70393 (2025).

41. Buzsáki, G. & Wang, X.-J. Mechanisms of Gamma Oscillations. Annu Rev Neurosci 35, 203–225 (2012).

42. Manning, J. R., Jacobs, J., Fried, I. & Kahana, M. J. Broadband Shifts in Local Field Potential Power Spectra Are Correlated with Single-Neuron Spiking in Humans. J. Neurosci. 29, 13613–13620 (2009).

43. Canolty, R. T. & Knight, R. T. The functional role of cross-frequency coupling. Trends in Cognitive Sciences 14, 506–515 (2010).

44. Engel, A. K., Gerloff, C., Hilgetag, C. C. & Nolte, G. Intrinsic Coupling Modes: Multiscale Interactions in Ongoing Brain Activity. Neuron 80, 867–886 (2013).

45. Uddin, L. Q. Salience processing and insular cortical function and dysfunction. Nat Rev Neurosci 16, 55–61 (2015).

46. Shine, J. M. Neuromodulatory Influences on Integration and Segregation in the Brain. Trends in Cognitive Sciences 23, 572–583 (2019).

47. Kristensen, A. S. et al. SLC6 neurotransmitter transporters: structure, function, and regulation. Pharmacol Rev 63, 585–640 (2011).

48. Siegel, M., Donner, T. H. & Engel, A. K. Spectral fingerprints of large-scale neuronal interactions. Nat Rev Neurosci 13, 121–134 (2012).

49. Logothetis, N. K., Pauls, J., Augath, M., Trinath, T. & Oeltermann, A. Neurophysiological investigation of the basis of the fMRI signal. Nature 412, 150–157 (2001).

50. Hipp, J. F. & Siegel, M. BOLD fMRI Correlation Reflects Frequency-Specific Neuronal Correlation. Current Biology 25, 1368–1374 (2015).

51. Brookes, M. J. et al. Investigating the electrophysiological basis of resting state networks using magnetoencephalography. Proceedings of the National Academy of Sciences 108, 16783–16788 (2011).

52. Sastre-Yagüe, D. et al. Cortical excitability inversely modulates fMRI connectivity via low-frequency neuronal coupling. 2026.03.12.710517 Preprint at 10.64898/2026.03.12.710517 (2026).

53. Schotte, A. et al. Risperidone compared with new and reference antipsychotic drugs: in vitro and in vivo receptor binding. Psychopharmacology 124, 57–73 (1996).

54. Nyberg, S., Farde, L., Eriksson, L., Halldin, C. & Eriksson, B. 5-HT2 and D2 dopamine receptor occupancy in the living human brain. Psychopharmacology 110, 265–272 (1993).

55. Mion, G. & Villevieille, T. Ketamine Pharmacology: An Update (Pharmacodynamics and Molecular Aspects, Recent Findings). CNS Neuroscience & Therapeutics 19, 370–380 (2013).

56. Wang, J., Sun, P. & Liang, P. Neuropsychopharmacological effects of midazolam on the human brain. Brain Inf. 7, 15 (2020).

57. Keiser, M., Hasan, M. & Oswald, S. Affinity of Ketamine to Clinically Relevant Transporters. Mol. Pharmaceutics 15, 326–331 (2018).

58. Kegeles, L. S. et al. Modulation of amphetamine-induced striatal dopamine release by ketamine in humans: implications for schizophrenia. Biological Psychiatry 48, 627–640 (2000).

59. Meyer, T. et al. Predictive value of heart rate in treatment of major depression with ketamine in two controlled trials. Clin Neurophysiol 132, 1339–1346 (2021).

60. Zhu, J., Spencer, T. J., Liu-Chen, L.-Y., Biederman, J. & Bhide, P. G. Methylphenidate and μ opioid receptor interactions: A pharmacological target for prevention of stimulant abuse. Neuropharmacology 61, 283–292 (2011).

61. Batistela, S., Bueno, O. F. A., Vaz, L. J. & Galduróz, J. C. F. Methylphenidate as a cognitive enhancer in healthy young people. Dement Neuropsychol 10, 134–142 (2016).

62. Pourhamzeh, M. et al. The Roles of Serotonin in Neuropsychiatric Disorders. Cell Mol Neurobiol 42, 1671–1692 (2021).

63. Tost, H., Alam, T. & Meyer-Lindenberg, A. Dopamine and Psychosis: Theory, Pathomechanisms and Intermediate Phenotypes. Neurosci Biobehav Rev 34, 689–700 (2010).

64. Royer, J. et al. Opportunities and pitfalls of data contextualization in neuroimaging. Nat. Rev. Neurosci. 1–19 (2026) doi:10.1038/s41583-026-01038-0.

65. Hansen, J. Y. & Misic, B. Integrating and interpreting brain maps. Trends in Neurosciences 48, 594–607 (2025).

66. Hansen, J. Y. et al. Inter-individual variability of neurotransmitter receptor and transporter density in the human brain. Brain Struct Funct 231, 13 (2026).

67. Glasser, M. F. et al. The minimal preprocessing pipelines for the Human Connectome Project. Neuroimage 80, 105–124 (2013).

68. Fischl, B. FreeSurfer. Neuroimage 62, 774–781 (2012).

69. Esteban, O. et al. fMRIPrep: a robust preprocessing pipeline for functional MRI. Nat Methods 16, 111–116 (2019).

70. Mehta, K. et al. XCP-D: A robust pipeline for the post-processing of fMRI data. Imaging Neuroscience 2, imag–2–00257 (2024).

71. Leon Lotter. mapconn. Zenodo 10.5281/ZENODO.19859027 (2026).

72. Harris, C. R. et al. Array programming with NumPy. Nature 585, 357–362 (2020).

73. McKinney, W. Data Structures for Statistical Computing in Python. in 56–61 (Austin, Texas, 2010). doi:10.25080/Majora-92bf1922-00a.

74. Virtanen, P. et al. SciPy 1.0: fundamental algorithms for scientific computing in Python. Nat Methods 17, 261–272 (2020).

75. Seabold, S. & Perktold, J. Statsmodels: Econometric and Statistical Modeling with Python. SciPy 2010 10.25080/Majora-92bf1922-011 (2010) doi:10.25080/Majora-92bf1922-011.

76. Vallat, R. Pingouin: statistics in Python. Journal of Open Source Software 3, 1026 (2018).

77. Pomponio, R. et al. Harmonization of large MRI datasets for the analysis of brain imaging patterns throughout the lifespan. NeuroImage 208, 116450 (2020).

78. Abraham, A. et al. Machine learning for neuroimaging with scikit-learn. Frontiers in Neuroinformatics 8, (2014).

79. Makowski, D. et al. NeuroKit2: A Python toolbox for neurophysiological signal processing. Behav Res 53, 1689–1696 (2021).

80. Hunter, J. D. Matplotlib: A 2D Graphics Environment. Computing in Science Engineering 9, 90–95 (2007).

81. Waskom, M. L. seaborn: statistical data visualization. Journal of Open Source Software 6, 3021 (2021).

82. Castrillon, G. et al. An energy costly architecture of neuromodulators for human brain evolution and cognition. Science Advances 9, eadi7632 (2023).

83. Finnema, S. J. et al. Imaging synaptic density in the living human brain. Sci Transl Med 8, 348ra96 (2016).

84. Finnema, S. J. et al. Kinetic evaluation and test-retest reproducibility of [11C]UCB-J, a novel radioligand for positron emission tomography imaging of synaptic vesicle glycoprotein 2A in humans. J Cereb Blood Flow Metab 38, 2041–2052 (2018).

85. Andersson, J. D., Matuskey, D. & Finnema, S. J. Positron emission tomography imaging of the γ-aminobutyric acid system. Neuroscience Letters 691, 35–43 (2019).

86. Chen, M.-K. et al. Comparison of [11C]UCB-J and [18F]FDG PET in Alzheimer’s disease: A tracer kinetic modeling study. J Cereb Blood Flow Metab 41, 2395–2409 (2021).

87. Chen, M.-K. et al. Assessing Synaptic Density in Alzheimer Disease With Synaptic Vesicle Glycoprotein 2A Positron Emission Tomographic Imaging. JAMA Neurol 75, 1215 (2018).

88. Holmes, S. E. et al. Lower synaptic density is associated with depression severity and network alterations. Nat Commun 10, 1529 (2019).

89. Wey, H.-Y. et al. Insights into neuroepigenetics through human histone deacetylase PET imaging. Sci. Transl. Med. 8, (2016).

90. Larsen, B. et al. Maturation of the human striatal dopamine system revealed by PET and quantitative MRI. Nat Commun 11, 846 (2020).

91. Luna, B. PETfrog. Openneuro 10.18112/OPENNEURO.DS002385.V1.1.0 (2021).

92. Smart, K. et al. Sex differences in [11C]ABP688 binding: a positron emission tomography study of mGlu5 receptors. Eur J Nucl Med Mol Imaging 46, 1179–1183 (2019).

93. Galovic, M. et al. Validation of a combined image derived input function and venous sampling approach for the quantification of [18F]GE-179 PET binding in the brain. NeuroImage 237, 118194 (2021).

94. Galovic, M. et al. In Vivo *N* -Methyl- D -Aspartate Receptor (NMDAR) Density as Assessed Using Positron Emission Tomography During Recovery From NMDAR-Antibody Encephalitis. JAMA Neurol 80, 211 (2023).

95. Nørgaard, M. et al. A high-resolution in vivo atlas of the human brain’s benzodiazepine binding site of GABAA receptors. Neuroimage 232, 117878 (2021).

96. Lukow, P. B. et al. Cellular and molecular signatures of in vivo imaging measures of GABAergic neurotransmission in the human brain. Commun Biol 5, 1–11 (2022).

97. Hesse, S. et al. Central noradrenaline transporter availability in highly obese, non-depressed individuals. Eur J Nucl Med Mol Imaging 44, 1056–1064 (2017).

98. Hillmer, A. T. et al. Imaging of cerebral α4β2* nicotinic acetylcholine receptors with (-)- [(18)F]Flubatine PET: Implementation of bolus plus constant infusion and sensitivity to acetylcholine in human brain. Neuroimage 141, 71–80 (2016).

99. Baldassarri, S. R. et al. Use of Electronic Cigarettes Leads to Significant Beta2-Nicotinic Acetylcholine Receptor Occupancy: Evidence From a PET Imaging Study. Nicotine & Tobacco Research 20, 425–433 (2018).

100. Naganawa, M. et al. First in Human Assessment of the Novel M1 Muscarinic Acetylcholine Receptor PET Radiotracer 11C-LSN3172176. Journal of Nuclear Medicine 10.2967/jnumed.120.246967 (2020) doi:10.2967/jnumed.120.246967.

101. Aghourian, M. et al. Quantification of brain cholinergic denervation in Alzheimer’s disease using PET imaging with [18F]-FEOBV. Mol Psychiatry 22, 1531–1538 (2017).

102. García Gómez, F., Huertas, I., Lojo Ramírez, J. & García Solís, D. Elaboración de una plantilla de SPM para la normalización de imágenes de PET con 18F-DOPA. ImagenDiagnostica 9, 23–25 (2018).

103. Kaller, S. et al. Test-retest measurements of dopamine D1-type receptors using simultaneous PET/MRI imaging. Eur J Nucl Med Mol Imaging 44, 1025–1032 (2017).

104. Sandiego, C. M. et al. Reference region modeling approaches for amphetamine challenge studies with [11C]FLB 457 and PET. J Cereb Blood Flow Metab 35, 623–629 (2015).

105. Sandiego, C. M. et al. Imaging robust microglial activation after lipopolysaccharide administration in humans with PET. Proceedings of the National Academy of Sciences 112, 12468–12473 (2015).

106. Sandiego, C. M. et al. The Effect of Treatment with Guanfacine, an Alpha2 Adrenergic Agonist, on Dopaminergic Tone in Tobacco Smokers: An [11C]FLB457 PET Study. Neuropsychopharmacol. 43, 1052–1058 (2018).

107. Beliveau, V. et al. A High-Resolution In Vivo Atlas of the Human Brain’s Serotonin System. J Neurosci 37, 120–128 (2017).

108. Radhakrishnan, R. et al. Age-Related Change in 5-HT6 Receptor Availability in Healthy Male Volunteers Measured with 11C-GSK215083 PET. J Nucl Med 59, 1445–1450 (2018).

109. Radhakrishnan, R. et al. In vivo 5-HT6 and 5-HT2A receptor availability in antipsychotic treated schizophrenia patients vs. unmedicated healthy humans measured with [11C]GSK215083 PET. Psychiatry Research: Neuroimaging 295, 111007 (2020).

110. Kantonen, T. et al. Interindividual variability and lateralization of μ-opioid receptors in the human brain. NeuroImage 217, 116922 (2020).

111. Vijay, A. et al. PET imaging reveals lower kappa opioid receptor availability in alcoholics but no effect of age. Neuropsychopharmacol 43, 2539–2547 (2018).

112. Normandin, M. D. et al. Imaging the cannabinoid CB1 receptor in humans with [11C]OMAR: assessment of kinetic analysis methods, test-retest reproducibility, and gender differences. J Cereb Blood Flow Metab 35, 1313–1322 (2015).

113. D’Souza, D. C. et al. Rapid Changes in Cannabinoid 1 Receptor Availability in Cannabis-Dependent Male Subjects After Abstinence From Cannabis. Biological Psychiatry: Cognitive Neuroscience and Neuroimaging 1, 60–67 (2016).

114. Ranganathan, M. et al. Reduced Brain Cannabinoid Receptor Availability in Schizophrenia. Biological Psychiatry 79, 997–1005 (2016).

115. Neumeister, A. et al. Positron Emission Tomography Shows Elevated Cannabinoid CB_1_ Receptor Binding in Men with Alcohol Dependence. Alcoholism Clin & Exp Res 36, 2104–2109 (2012).

116. Hoffmann, M., Hoopes, A., Greve, D. N., Fischl, B. & Dalca, A. V. Anatomy-aware and acquisition-agnostic joint registration with SynthMorph. Imaging Neuroscience 2, imag–2–00197 (2024).

117. Sydnor, V. J. et al. Intrinsic Activity Develops Along a Sensorimotor-Association Cortical Axis in Youth. 2022.08.15.503994 Preprint at 10.1101/2022.08.15.503994 (2022).

118. Paquola, C. et al. The BigBrainWarp toolbox for integration of BigBrain 3D histology with multimodal neuroimaging. eLife 10, e70119 (2021).

119. Amunts, K. et al. BigBrain: An Ultrahigh-Resolution 3D Human Brain Model. Science 340, 1472–1475 (2013).

120. Lorio, S. et al. New tissue priors for improved automated classification of subcortical brain structures on MRI. NeuroImage 130, 157–166 (2016).

121. Huck, J. et al. High resolution atlas of the venous brain vasculature from 7 T quantitative susceptibility maps. Brain Struct Funct 224, 2467–2485 (2019).

122. Mouches, P. & Forkert, N. D. A statistical atlas of cerebral arteries generated using multi-center MRA datasets from healthy subjects. Sci Data 6, 29 (2019).

123. Schaefer, A. et al. Local-Global Parcellation of the Human Cerebral Cortex from Intrinsic Functional Connectivity MRI. Cereb Cortex 28, 3095–3114 (2018).

124. Kangjoo Lee et al. Yale Resting State fMRI/Pupillometry: Arousal Study. Openneuro 10.18112/OPENNEURO.DS003673.V2.0.1 (2022).

125. Jacobs, G. R. et al. An Introduction to the Human Connectome Project for Early Psychosis. Schizophrenia Bulletin sbae123 (2024) doi:10.1093/schbul/sbae123.

126. Larabi, D. I., Gell, M., Amico, E., Eickhoff, S. B. & Patil, K. R. Spatially localized fMRI metrics as predictive and highly distinct state-independent fingerprints. 2021.08.03.454862 Preprint at 10.1101/2021.08.03.454862 (2022).

127. Tadel, F., Baillet, S., Mosher, J. C., Pantazis, D. & Leahy, R. M. Brainstorm: A User-Friendly Application for MEG/EEG Analysis. Computational Intelligence and Neuroscience 2011, 879716 (2011).

128. Bruns, A., Eckhorn, R., Jokeit, H. & Ebner, A. Amplitude envelope correlation detects coupling among incoherent brain signals. Neuroreport 11, 1509–1514 (2000).

129. Wens, V. et al. A geometric correction scheme for spatial leakage effects in MEG/EEG seed-based functional connectivity mapping. Hum Brain Mapp 36, 4604–4621 (2015).

130. Colclough, G. L. et al. How reliable are MEG resting-state connectivity metrics? Neuroimage 138, 284–293 (2016).

131. Palva, J. M. et al. Ghost interactions in MEG/EEG source space: A note of caution on inter-areal coupling measures. Neuroimage 173, 632–643 (2018).

132. Alexander-Bloch, A. F. et al. On testing for spatial correspondence between maps of human brain structure and function. Neuroimage 178, 540–551 (2018).

133. Vos de Wael, R., et al. BrainSpace: a toolbox for the analysis of macroscale gradients in neuroimaging and connectomics datasets. Commun Biol 3, 1–10 (2020).

134. Wagner, H. H. & Dray, S. Generating spatially constrained null models for irregularly spaced data using Moran spectral randomization methods. Methods in Ecology and Evolution 6, 1169–1178 (2015).

135. Li, J. & Ji, L. Adjusting multiple testing in multilocus analyses using the eigenvalues of a correlation matrix. Heredity 95, 221–227 (2005).

136. Whitlock, M. C. Combining probability from independent tests: the weighted Z-method is superior to Fisher’s approach. Journal of Evolutionary Biology 18, 1368–1373 (2005).

137. Kret, M. E. & Sjak-Shie, E. E. Preprocessing pupil size data: Guidelines and code. Behav Res 51, 1336–1342 (2019).

138. Lüdtke, H., Wilhelm, B., Adler, M., Schaeffel, F. & Wilhelm, H. Mathematical procedures in data recording and processing of pupillary fatigue waves. Vision Research 38, 2889–2896 (1998).

139. Wilhelm, B., Wilhelm, H., Lüdtke, H., Streicher, P. & Adler, M. Pupillographic assessment of sleepiness in sleep-deprived healthy subjects. Sleep 21, 258–265 (1998).

140. Rizzuto, V. et al. Pupillary Hippus as a Biomarker: Spectral Signatures and Complexity Approaches in Autonomic and Clinical Contexts. Bioengineering (Basel) 12, 1376 (2025).

141. Heart rate variability: standards of measurement, physiological interpretation and clinical use. Task Force of the European Society of Cardiology and the North American Society of Pacing and Electrophysiology. Circulation 93, 1043–1065 (1996).

142. Johnson, W. E., Li, C. & Rabinovic, A. Adjusting batch effects in microarray expression data using empirical Bayes methods. Biostatistics 8, 118–127 (2007).

143. Fortin, J.-P. et al. Harmonization of cortical thickness measurements across scanners and sites. NeuroImage 167, 104–120 (2018).

