## Supplementary Methods and Results for "Linking human brain functional connectivity to underlying neurotransmitter systems"

**Author Note**

*Correspondence to:

**Supplementary Figures**

See separate PDF document.

**Supplementary Tables**

See separate Excel file.

**Animation S1**

See separate GIF file.

### Supplementary Text

#### Supplementary Results

##### *Discovery analyses robustness checks*

We performed several analyses to test for the stability of our main results. Inclusion of subcortical parcels often increased effect estimates compared to null effects (Fig. 2c), while different parcellation resolutions had effects comparable to those observed for resting-state networks (Fig. S6; Tab. S4). Analyses without interhemispheric connections resulted in only slightly decreased effect sizes (Fig. S7a; Tab. S4), proving that the null models correctly accounted for interhemispheric symmetry effects. Using the fit of a polynomial as an alternative aggregation method, we observed related but not fully similar AUC profiles (Fig. S7b; Tab. S4), indicating potential for future exploration of different aggregate metrics.

##### *MEG leakage-correction (orthogonalized AEC) sensitivity analysis*

To assess whether the observed MEG AUC effects reflect residual, spatially inhomogeneous signal leakage not captured by our spatial null models, we repeated all MEG analyses using connectomes (amplitude envelope correlation) computed after pairwise orthogonalization[1]. As expected, given the unspecific removal of shared zero-lag signal by orthogonalization, mean AUC+ effect sizes were substantially reduced across all frequency bands (61–113% reduction relative to the non-orthogonalized analysis, all $p_{Meff} > 0.05$; Tab. S9). Despite this overall attenuation, AUC profiles across the 25 reference atlases remained significantly correlated between the orthogonalized and non-orthogonalized analyses in several bands. This correlation was most consistent for AUC– profiles, which were significantly correlated in every frequency band (rho = 0.48–0.90, all $p < 0.05$). AUC+ profile correlations were more variable, reaching significance in the delta, alpha, and beta bands (rho = 0.40–0.71, $p < 0.05$; Tab. S10). Notably, the two dominant AUC+ effects across all analyses, NET and VAChT, showed markedly different sensitivity to orthogonalization. The VAChT effect, significant in the non-orthogonalized analysis ($p = 0.005$, $p_{Meff} = 0.043$), was rendered insignificant by orthogonalization ($p = 0.52$, $p_{Meff} = 0.998$), consistent with this effect being substantially affected by signal leakage. In contrast, the NET effect remained nominally significant and virtually unchanged in its uncorrected p value ($p = 0.039$ vs. 0.037, $p_{Meff} > 0.05$ in both conditions). This dissociation suggests that the noradrenergic AUC+ effect is not primarily attributable to spatially inhomogeneous MEG signal leakage, whereas the cholinergic (VAChT) effect should be interpreted with greater caution.

#### Supplementary Methods

##### *Study samples and subject exclusions*

Study samples were obtained from open data repositories and multiple individual studies. For all rsfMRI datasets except for the MPH study, for which we obtained connectomes directly, scan sessions were consistently excluded if they exhibited too much motion (mean frame-wise displacement > 0.3).

To assemble the rsfMRI discovery sample, we selected 132 subjects from the full HCP-YA dataset. These were the union of the “100 unrelated subjects” and those for which we had MEG data available. We retained only subjects that had a full dataset of two runs (mostly one day apart) with two scans each according to phase encoding directions (“LR” and “RL”). After exclusion of subjects without full rsfMRI data (n = 8) and high motion in at least scan (n = 12), a final sample of 112 subjects remained. We used 3 Tesla fMRI data obtained as described by Van-Essen et al.[2]. The HCP-YA MEG subdataset consisted of data from 33 subjects, which were selected, processed, and provided by Shafiei et al.[3]. Finally, of 112 fMRI subjects, 64 participants with 103 sessions had high-quality heart-rate variability data for post-hoc association analyses (see below).

The YRSP rsfMRI/pupillometry dataset was downloaded from openneuro. Scanning parameters (3 Tesla) and pupillometry setups were described by Lee et al.[4,5]. Pupillometry data was obtained simultaneously with MRI scanning using an infrared eye-tracking system. Of 27 subjects with two runs each, a final sample of 26 subjects passed motion thresholds for both runs. After quality control of the pupillometry data (see below), 25 subjects with 49 sessions went into the rsfMRI-pupil diameter association analyses.

The MPH data was provided and processed by Dipasquale et al.[6]; scanning parameters (1.5 Tesla) and preprocessing is described in the original publication. The 30 included subjects received 20 mg of oral MPH or placebo 90 minutes before MRI scanning in a double-blind, randomized, within-subjects design with two sessions at least one week apart.

The risperidone study (3 Tesla) was described in detail by Hawkins et al.[7]. Twenty-one participants received oral doses of either 0.5 mg of risperidone, 2 mg of risperidone, or placebo 120 minutes before MRI scanning in a double-blind, randomized, within-subjects design with three sessions at least one week apart. The final sample, after motion criteria were applied, consisted of 19, 17, and 17 sessions for the low-dose, high-dose, and placebo conditions, respectively.

Details on the ketamine/midazolam study (3 Tesla) were provided by Forsyth et al.[8]. Thirty subjects received either ketamine (0.25 mg/kg bolus plus 0.25 mg/kg/h), midazolam (0.03 mg/kg bolus plus 0.03 mg/kg/h), or placebo intravenously at subanesthetic doses starting after minute 7 within a 17 minute rsfMRI scan. The study was conducted in a single-blind, randomized, within-subjects design with three sessions at least 48 hours apart. For the main analyses, the rsfMRI scan was split into a pre- and a post-treatment run (the first and last ~7 minutes, i.e., 190 slices with 2.2 seconds repetition time). Subject numbers after exclusions were 28, 28, and 24 for the placebo, ketamine, and midazolam sessions, respectively.

The HCP-EP study[9] sampled individuals across 4 sites within the first five years of onset of psychotic symptoms, along with a control group. The psychosis group is cross-diagnostic along the psychosis spectrum including individuals with diagnoses of schizophrenia, schizophreniform, schizoaffective, delusional, and brief psychotic disorder, as well as non-specified psychosis and affective disorders with psychosis. We used the first run of each subject within two scans with different phase encoding directions ("AP" and "PA"). Of the full sample, 159 subjects had MRI and the required demographic and clinical data available; of these, 151 passed motion thresholds for both runs (n = 96 cases, n = 55 controls).

***Resting-state fMRI processing***

Preprocessing, denoising, and Pearson connectome extraction of the MPH data was performed by Dipasquale et al.[6]. The HCP-YA data was obtained directly in "minimally preprocessed" format (without FIX denoising). The HCP-EP data was preprocessed using the HCP pipeline ("GenericfMRIVolumeProcessingPipeline")[10] as described by Larabi et al.[11]. All other datasets were preprocessed in-house using FreeSurfer and fMRIprep. Following separately conducted FreeSurfer "recon-all" runs, processing of the structural images included intensity non-uniformity correction, skull-stripping, tissue segmentation, spatial normalization to MNI152NLin6Asym space, and resampling onto the fsLR greyordinates (91k density). Per run of the functional preprocessing stream, head motion was corrected relative to a reference volume, this reference was coregistered to the structural image, and slice timing-correction was performed if the timing information was available. Using the derived transforms, the BOLD time-series were then resampled onto the left/right-symmetric fsLR 91k surface and the associated subcortical volume in 2mm MNI152NLin6Asym space.

Except for the MPH dataset, postprocessing and connectome extraction was performed using XCP-D based on fsLR greyordinate data from the HCP- or fMRIprep pipeline outputs. Framewise displacement[12] was calculated from the estimated motion parameters (50 mm head radius) to derive the scan-level estimate used for motion outlier identification and as a session/group-level covariate in post-hoc analyses. To ensure solid control of confounding physiological signals while retaining as much neural signal as possible, we employed the "36P" nuisance regression strategy with despiking of the BOLD signals[13]. No censoring of high-motion volumes was performed, to avoid unnecessary interpolation of BOLD signals. The 36 nuisance regressors included six motion and three tissue signals (mean global, white matter, and cerebrospinal fluid) together with their temporal derivatives, and quadratic expansion of the original parameters and their derivatives. The time-series and the confounds were band-pass filtered (second-order Butterworth; 0.01–0.08 Hz), and the BOLD time-series were denoised using linear regression. Average time-series were then extracted from the residual BOLD signals using the Schaefer[14] and the HCP CIFTI subcortical[10] parcellations, the latter of which corresponds to the FreeSurfer subcortical parcellation. If a parcel had > 50% of vertices uncovered (values of all zeros or NaNs), these vertices were ignored. If it had < 50% coverage, the parcel time-series was set to zero. Finally, pair-wise Pearson functional connectivity matrices were calculated between all parcel time-series and Fisher z-transformed. All-zero time-series due to uncovered vertices resulted in all-NaN rows and columns in the connectome matrices. For HCP-YA and HCP-EP, connectomes were computed separately per phase-encoding direction and then averaged within runs.

#### Physiological data processing and quality control

##### *Pupilometry data*

Pupil area time series from the YRSP dataset were provided in minimally preprocessed form (blink interpolation, downsampled to 1 Hz). We additionally excluded samples outside a lenient plausible amplitude range based on visual inspection of the data (300, 10,000 arbitrary units), applied a square-root transform to approximate a linear diameter scale, and removed samples whose first-difference exceeded the median velocity plus five times the MAD (scaled by 1.4826)[15]. Resulting gaps were linearly interpolated up to 5 data points; longer gaps were replaced by NaN values. Sessions with fewer than 80% valid data points after cleaning were excluded. The following measures were derived: the pupil unrest index (PUI)[16,17], defined as the root mean square of the first-differences of the filtered signal, and Welch spectral power (Hann window, 256-sample segments, constant detrending) in three frequency bands estimated from the longest artifact-free segment of each run: sympathetic-linked low-frequency slow oscillations (LF-S: 0.04–0.10 Hz), respiration-linked low-frequency (LF-R: 0.10–0.24 Hz), and parasympathetic-linked high-frequency (HF-P: 0.25–0.50 Hz; upper limit set to Nyquist)[18]. All three spectral measures were log-transformed prior to statistical analyses.

##### *Heart rate data*

Heart rate variability was derived from photoplethysmography (PPG) signals recorded at 400 Hz during HCP-YA rsfMRI scanning. PPG signals were processed using NeuroKit2 (Elgendi method, robust estimation), and two complementary HRV estimates, RMSSD[19] (square root of the mean of squared successive differences between adjacent RR intervals) and MadNN (median absolute deviation of RR intervals), were computed per phase-encoding direction and run. To align with the connectome data, values were then averaged across directions within each run. Quality control was applied per metric: sessions were included if the mean NeuroKit2 PPG signal quality index exceeded a data-driven threshold (knee point of its empirical cumulative distribution) and if HRV values did not exceed a realistic threshold of 150 ms.

#### Supplementary References


1. Hipp, J. F., Hawellek, D. J., Corbetta, M., Siegel, M. & Engel, A. K. Large-scale cortical correlation structure of spontaneous oscillatory activity. *Nat Neurosci* **15**, 884–890 (2012).
2. Van Essen, D. C. *et al.* The WU-Minn Human Connectome Project: An overview. *NeuroImage* **80**, 62–79 (2013).
3. Shafiei, G., Baillet, S. & Misic, B. Human electromagnetic and haemodynamic networks systematically converge in unimodal cortex and diverge in transmodal cortex. *PLOS Biology* **20**, e3001735 (2022).
4. Lee, K. *et al.* Arousal impacts distributed hubs modulating the integration of brain functional connectivity. *NeuroImage* **258**, 119364 (2022).
5. Kangjoo Lee *et al.* Yale Resting State fMRI/Pupillometry: Arousal Study. Openneuro https://doi.org/10.18112/OPENNEURO.DS003673.V2.0.1 (2022).
6. Dipasquale, O. *et al.* Unravelling the effects of methylphenidate on the dopaminergic and noradrenergic functional circuits. *Neuropsychopharmacol.* **45**, 1482–1489 (2020).
7. Hawkins, P. C. T. *et al.* The effect of risperidone on reward-related brain activity is robust to drug-induced vascular changes. *Human Brain Mapping* **42**, 2766–2777 (2021).
8. Forsyth, A. *et al.* Comparison of local spectral modulation, and temporal correlation, of simultaneously recorded EEG/fMRI signals during ketamine and midazolam sedation. *Psychopharmacology* **235**, 3479–3493 (2018).
9. Jacobs, G. R. *et al.* An Introduction to the Human Connectome Project for Early Psychosis. *Schizophrenia Bulletin* sbae123 (2024) doi:10.1093/schbul/sbae123.
10. Glasser, M. F. *et al.* The minimal preprocessing pipelines for the Human Connectome Project. *Neuroimage* **80**, 105–124 (2013).
11. Larabi, D. I., Gell, M., Amico, E., Eickhoff, S. B. & Patil, K. R. Spatially localized fMRI metrics as predictive and highly distinct state-independent fingerprints. 2021.08.03.454862 Preprint at https://doi.org/10.1101/2021.08.03.454862 (2022).
12. Power, J. D. *et al.* Methods to detect, characterize, and remove motion artifact in resting state fMRI. *NeuroImage* **84**, 320–341 (2014).
13. Ciric, R. *et al.* Benchmarking of participant-level confound regression strategies for the control of motion artifact in studies of functional connectivity. *NeuroImage* **154**, 174–187 (2017).
14. Schaefer, A. *et al.* Local-Global Parcellation of the Human Cerebral Cortex from Intrinsic Functional Connectivity MRI. *Cereb Cortex* **28**, 3095–3114 (2018).
15. Kret, M. E. & Sjak-Shie, E. E. Preprocessing pupil size data: Guidelines and code. *Behav Res* **51**, 1336–1342 (2019).
16. Wilhelm, B., Wilhelm, H., Lüdtke, H., Streicher, P. & Adler, M. Pupillographic assessment of sleepiness in sleep-deprived healthy subjects. *Sleep* **21**, 258–265 (1998).
17. Lüdtke, H., Wilhelm, B., Adler, M., Schaeffel, F. & Wilhelm, H. Mathematical procedures in data recording and processing of pupillary fatigue waves. *Vision Research* **38**, 2889–2896 (1998).
18. Rizzuto, V. *et al.* Pupillary Hippus as a Biomarker: Spectral Signatures and Complexity Approaches in Autonomic and Clinical Contexts. *Bioengineering (Basel)* **12**, 1376 (2025).
19. Heart rate variability: standards of measurement, physiological interpretation and clinical use. Task Force of the European Society of Cardiology and the North American Society of Pacing and Electrophysiology. *Circulation* **93**, 1043–1065 (1996).
