## Supplementary Figures for "Linking human brain functional connectivity to underlying neurotransmitter systems"


**Fig. S1**: NEOFC curves derived from resting-state network probability

**Fig. S2**: NEOFC curves derived from neurobiological atlases

**Fig. S3**: Individual-level p values associated to AUC scores

**Fig. S3a**: Schaefer200

**Fig. S3b**: Schaefer200+Subcortical

**Fig. S4**: Replication of AUC profiles in adult control samples

**Fig. S5**: Meta-Analysis of AUC p values from six MRI datasets

**Fig. S6**: AUC results including AUC-delta for different parcellation resolutions

**Fig. S6a**: Schaefer100(+Subcortical)

**Fig. S6b**: Schaefer200(+Subcortical)

**Fig. S6c**: Schaefer400(+Subcortical)

**Fig. S7**: AUC results without interhemispheric connections and an alternative aggregate metric

**Fig. S7a**: No interhemispheric connections

**Fig. S7b**: Polynomial fit aggregation

**Fig. S8**: NEOFC curves across MEG frequency bands

**Fig. S8a**: delta

**Fig. S8b**: theta

**Fig. S8c**: alpha

**Fig. S8d**: beta

**Fig. S8e**: lgamma

**Fig. S8f**: hgamma

**Fig. S9**: MEG AUC results with different parcellation resolutions

**Fig. S9a**: Schaefer100

**Fig. S9b**: Schaefer400

**Fig. S10**: MEG AUC results after orthogonalization

**Fig. S10a**: AUC results

**Fig. S10b**: Correspondence of AUC profiles

**Fig. S11**: Correspondence of fMRI and MEG AUC profiles across all frequency bands

**Fig. S12**: Correlation of atlas-wise AUC scores between MRI and MEG

**Fig. S13**: Leave-one-out analysis to identify influential regions and connections

**Fig. S13a**: AUC+

**Fig. S13b**: AUC–

**Fig. S14**: NEOFC curves with AUC estimation restricted to lowered percentiles

**Fig. S14a**: Schaefer200

**Fig. S14b**: Schaefer200+Subcortical

**Fig. S15**: Effects of covariate regression on NET AUC results

**Fig. S16**: Quality control of HCP-YA physiological data (heart-rate variability)

**Fig. S17**: Quality control of YRSP physiological data (pupil diameter over time)

**Fig. S18**: Associations of AUC scores with clinical variables

### Fig. S1

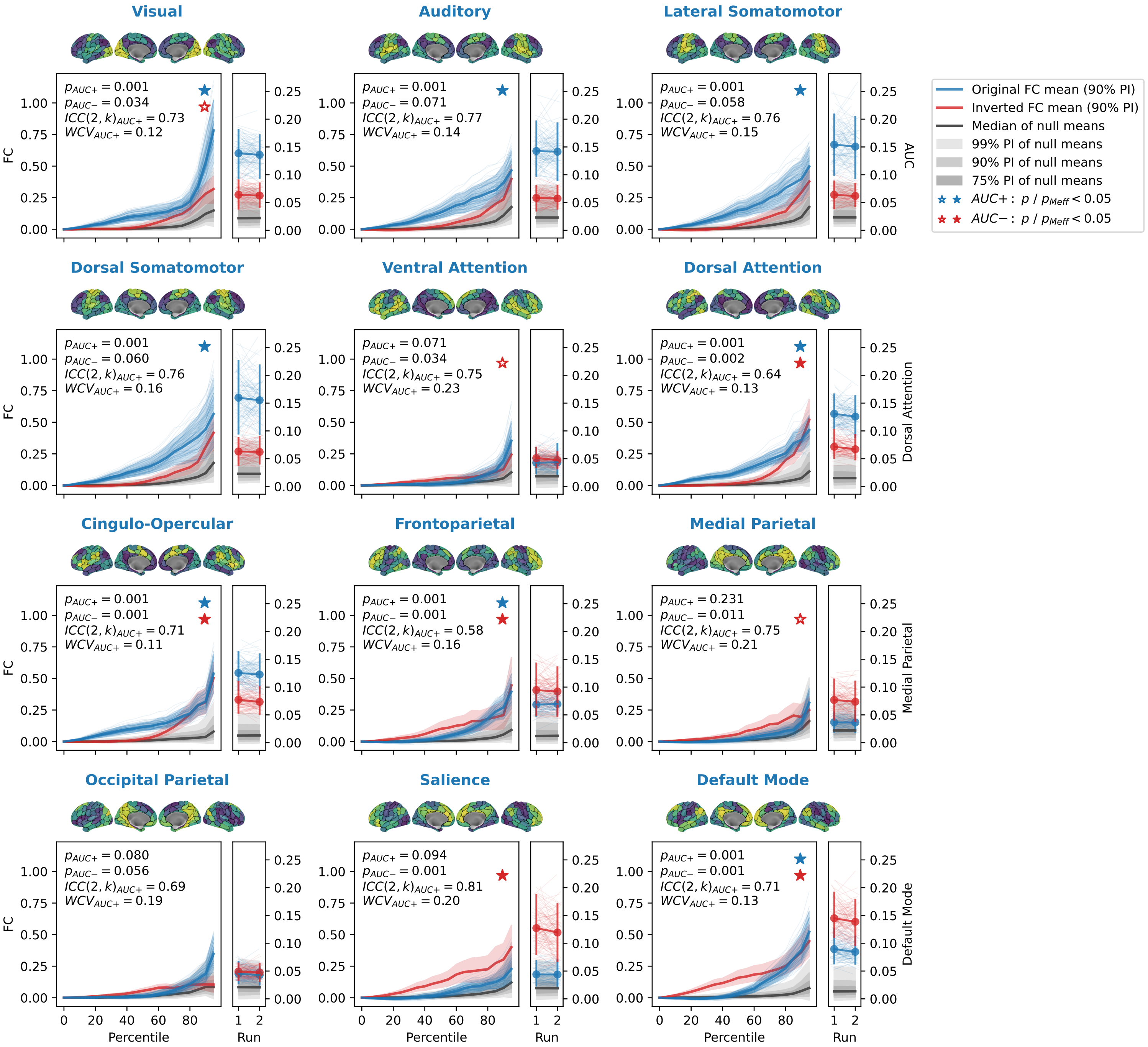

Visual
Auditory
Lateral Somatomotor
Dorsal Somatomotor
Ventral Attention
Dorsal Attention
Cingulo-Opercular
Frontoparietal
Medial Parietal
Occipital Parietal
Salience
Default Mode
FC
AUC
Percentile
Run
Original FC mean (90% PI)
Inverted FC mean (90% PI)
Median of null means
99% PI of null means
90% PI of null means
75% PI of null means
AUC+ : p / p_Meff < 0.05
AUC− : p / p_Meff < 0.05

#### **Fig. S1**: NEOFC curves derived from resting-state network probability

NEOFC curves for all 12 RSN probability atlases used as positive controls (Schaefer200 parcellation). Each panel shows mean rsFC as a function of the density percentile threshold for the original atlas (blue) and its spatially inverted reference (dark red). Colored shaded area: 90% PI across individuals; grey shaded area: null distribution. Right inset: individual AUC+ and AUC– scores (each dot: one participant). Displayed as in **Fig. 2b**. Brain maps indicate the parcellated RSN probability atlas, converted to percentiles. Abbreviations: rsFC: resting-state functional connectivity; AUC: area under the curve; RSN: resting-state network; PI: percentile interval.

**Fig. S2**

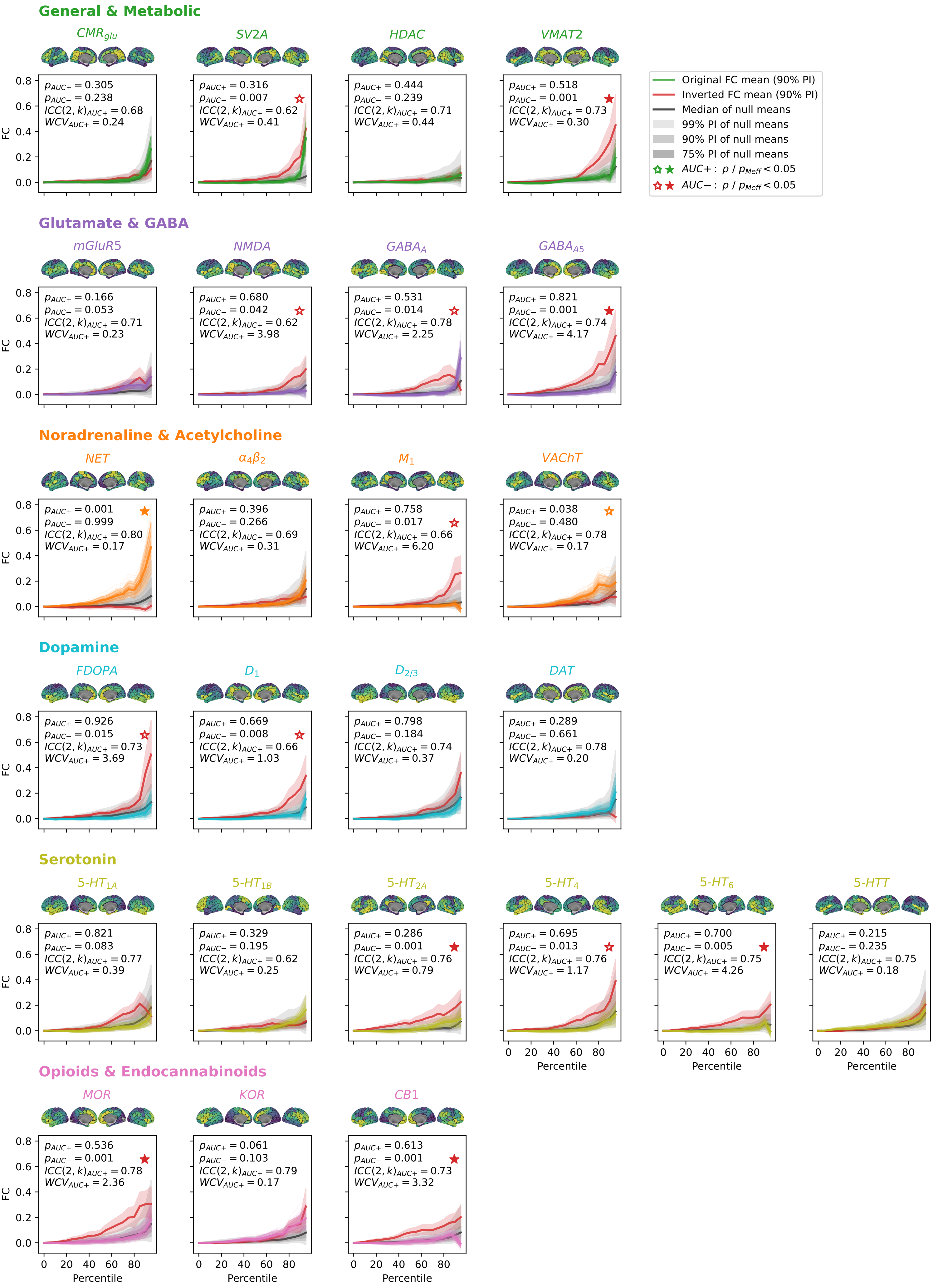
General & Metabolic
CMR_glu
p_AUC+ = 0.305
p_AUC− = 0.238
ICC(2,k)_AUC+ = 0.68
WCV_AUC+ = 0.24
SV2A
p_AUC+ = 0.316
p_AUC− = 0.007
ICC(2,k)_AUC+ = 0.62
WCV_AUC+ = 0.41
HDAC
p_AUC+ = 0.444
p_AUC− = 0.239
ICC(2,k)_AUC+ = 0.71
WCV_AUC+ = 0.44
VMAT2
p_AUC+ = 0.518
p_AUC− = 0.001
ICC(2,k)_AUC+ = 0.73
WCV_AUC+ = 0.30
Original FC mean (90% PI)
Inverted FC mean (90% PI)
Median of null means
99% PI of null means
90% PI of null means
75% PI of null means
AUC+ : p / p_Meff < 0.05
AUC− : p / p_Meff < 0.05
Glutamate & GABA
mGluR5
p_AUC+ = 0.166
p_AUC− = 0.053
ICC(2,k)_AUC+ = 0.71
WCV_AUC+ = 0.23
NMDA
p_AUC+ = 0.680
p_AUC− = 0.042
ICC(2,k)_AUC+ = 0.62
WCV_AUC+ = 3.98
GABA_A
p_AUC+ = 0.531
p_AUC− = 0.014
ICC(2,k)_AUC+ = 0.78
WCV_AUC+ = 2.25
GABA_A5
p_AUC+ = 0.821
p_AUC− = 0.001
ICC(2,k)_AUC+ = 0.74
WCV_AUC+ = 4.17
Noradrenaline & Acetylcholine
NET
p_AUC+ = 0.001
p_AUC− = 0.999
ICC(2,k)_AUC+ = 0.80
WCV_AUC+ = 0.17
α4β2
p_AUC+ = 0.396
p_AUC− = 0.266
ICC(2,k)_AUC+ = 0.69
WCV_AUC+ = 0.31
M1
p_AUC+ = 0.758
p_AUC− = 0.017
ICC(2,k)_AUC+ = 0.66
WCV_AUC+ = 6.20
VAChT
p_AUC+ = 0.038
p_AUC− = 0.480
ICC(2,k)_AUC+ = 0.78
WCV_AUC+ = 0.17
Dopamine
FDOPA
p_AUC+ = 0.926
p_AUC− = 0.015
ICC(2,k)_AUC+ = 0.73
WCV_AUC+ = 3.69
D1
p_AUC+ = 0.669
p_AUC− = 0.008
ICC(2,k)_AUC+ = 0.66
WCV_AUC+ = 1.03
D2/3
p_AUC+ = 0.798
p_AUC− = 0.184
ICC(2,k)_AUC+ = 0.74
WCV_AUC+ = 0.37
DAT
p_AUC+ = 0.289
p_AUC− = 0.661
ICC(2,k)_AUC+ = 0.78
WCV_AUC+ = 0.20
Serotonin
5-HT1A
p_AUC+ = 0.821
p_AUC− = 0.083
ICC(2,k)_AUC+ = 0.77
WCV_AUC+ = 0.39
5-HT1B
p_AUC+ = 0.329
p_AUC− = 0.195
ICC(2,k)_AUC+ = 0.62
WCV_AUC+ = 0.25
5-HT2A
p_AUC+ = 0.286
p_AUC− = 0.001
ICC(2,k)_AUC+ = 0.76
WCV_AUC+ = 0.79
5-HT4
p_AUC+ = 0.695
p_AUC− = 0.013
ICC(2,k)_AUC+ = 0.76
WCV_AUC+ = 1.17
5-HT6
p_AUC+ = 0.700
p_AUC− = 0.005
ICC(2,k)_AUC+ = 0.75
WCV_AUC+ = 4.26
5-HTT
p_AUC+ = 0.215
p_AUC− = 0.235
ICC(2,k)_AUC+ = 0.75
WCV_AUC+ = 0.18
Opioids & Endocannabinoids
MOR
p_AUC+ = 0.536
p_AUC− = 0.001
ICC(2,k)_AUC+ = 0.78
WCV_AUC+ = 2.36
KOR
p_AUC+ = 0.061
p_AUC− = 0.103
ICC(2,k)_AUC+ = 0.79
WCV_AUC+ = 0.17
CB1
p_AUC+ = 0.613
p_AUC− = 0.001
ICC(2,k)_AUC+ = 0.73
WCV_AUC+ = 3.32
FC
0.0 0.2 0.4 0.6 0.8
Percentile
0 20 40 60 80

### **Fig. S2**: NEOFC curves derived from neurobiological atlases

NEOFC curves for all 25 nuclear imaging reference maps, organized by broad neurotransmitter systems (Schaefer200 and Schaefer200 + subcortex parcellation). Each panel shows mean rsFC as a function of the density percentile threshold for the original atlas (colored by neurotransmitter system) and its spatially inverted reference (dark red). Colored shaded area: 90% PI across individuals; grey shaded area: null distribution. Brain maps indicate the parcellated reference atlas, converted to percentiles. Statistics printed in each plot include the raw p values for AUC+ and AUC–, the ICC(2,k), and the WCV for AUC+. Displayed as in **Fig. 2d**. Abbreviations: rsFC: resting-state functional connectivity; PI: percentile interval; ICC: intraclass correlation coefficient; WCV: within-subject coefficient of variability; see **Fig. 2** for reference map abbreviations.

**Fig. S3a**: Schaefer200

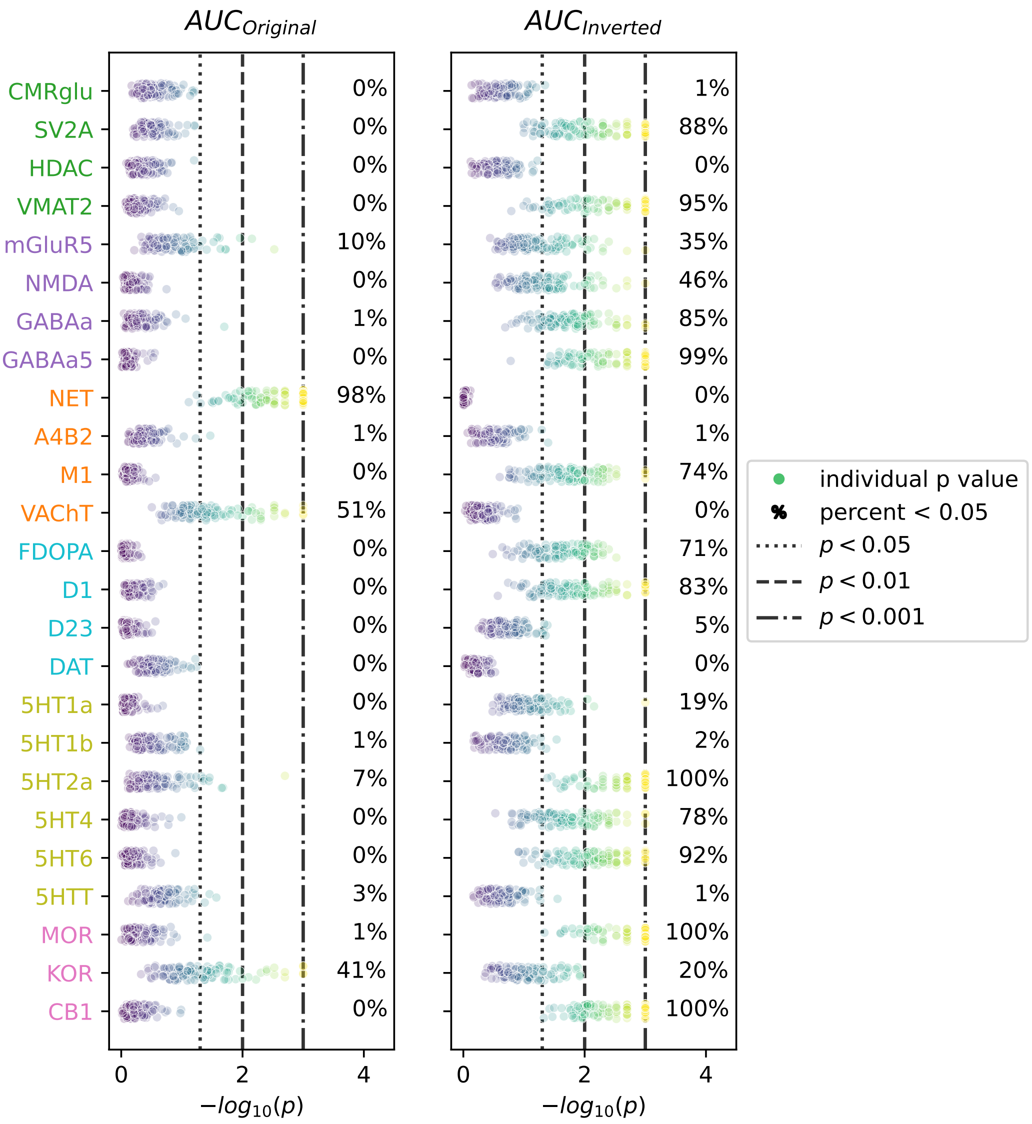

**Fig. S3b**: Schaefer200+Subcortical

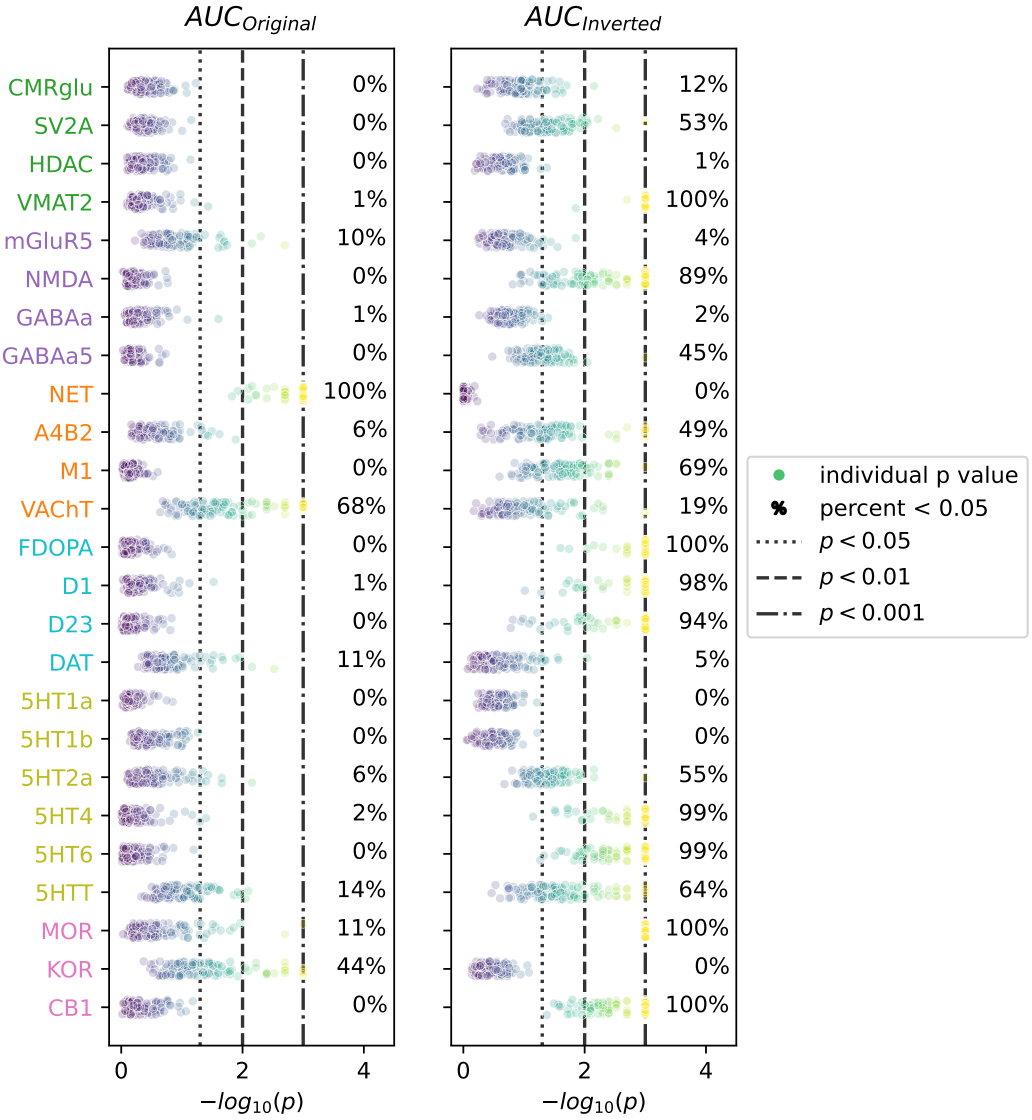

**Fig. S3**: Individual-level p values associated to AUC scores

Distribution of individual-level –log10(p) values for AUC+ (left) and AUC– (right) across 25 nuclear imaging reference maps. **a**: Schaefer200 parcellation; **b**: Schaefer200 + subcortex parcellation. Each dot represents one participant; color indicates –log10(p). Vertical lines indicate significance thresholds at $p < 0.05$ (dotted), $p < 0.01$ (dashed), and $p < 0.001$ (dot-dashed). Abbreviations: AUC: area under the curve; see **Fig. 2** for reference map abbreviations.

**Fig. S4**

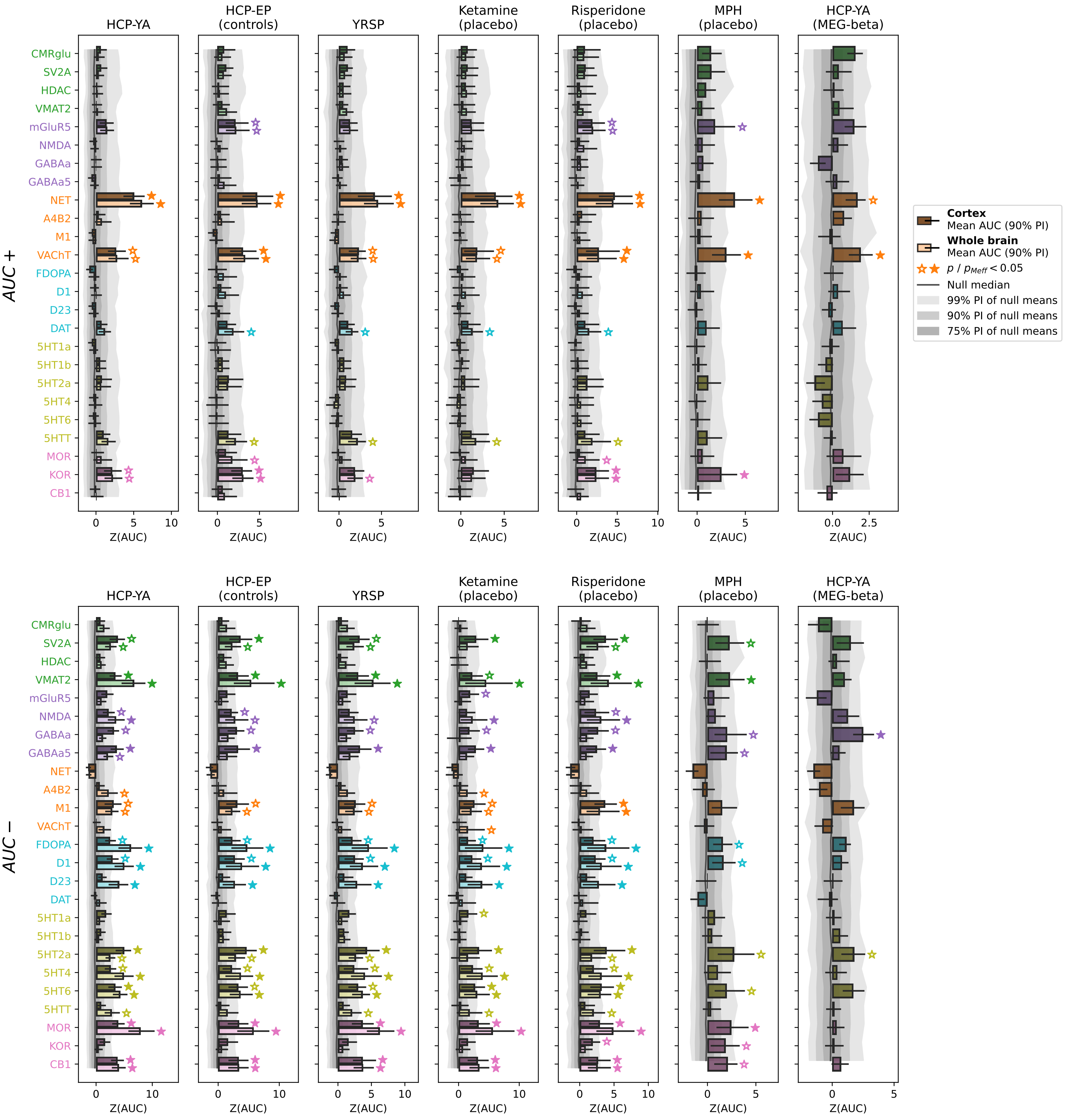
HCP-YA
HCP-EP (controls)
YRSP
Ketamine (placebo)
Risperidone (placebo)
MPH (placebo)
HCP-YA (MEG-beta)
AUC +
AUC −
CMRglu
SV2A
HDAC
VMAT2
mGluR5
NMDA
GABAa
GABAa5
NET
A4B2
M1
VAChT
FDOPA
D1
D23
DAT
5HT1a
5HT1b
5HT2a
5HT4
5HT6
5HTT
MOR
KOR
CB1
Z(AUC)
Cortex
Mean AUC (90% PI)
Whole brain
Mean AUC (90% PI)
$p$ / $p_{Meff} < 0.05$
Null median
99% PI of null means
90% PI of null means
75% PI of null means

#### **Fig. S4**: Replication of AUC profiles in adult control samples

AUC+ and AUC– scores (z-scored against a null distribution of spatially matched reference maps) for 25 nuclear imaging reference maps, shown separately for six independent adult control samples. Displayed as in **Fig. 2c** (Schaefer200 and Schaefer200 + subcortex parcellations). Abbreviations: AUC: area under the curve; PI: percentile interval; WCV: within-subject coefficient of variability; see **Fig. 2** for reference map abbreviations.

**Fig. S5**

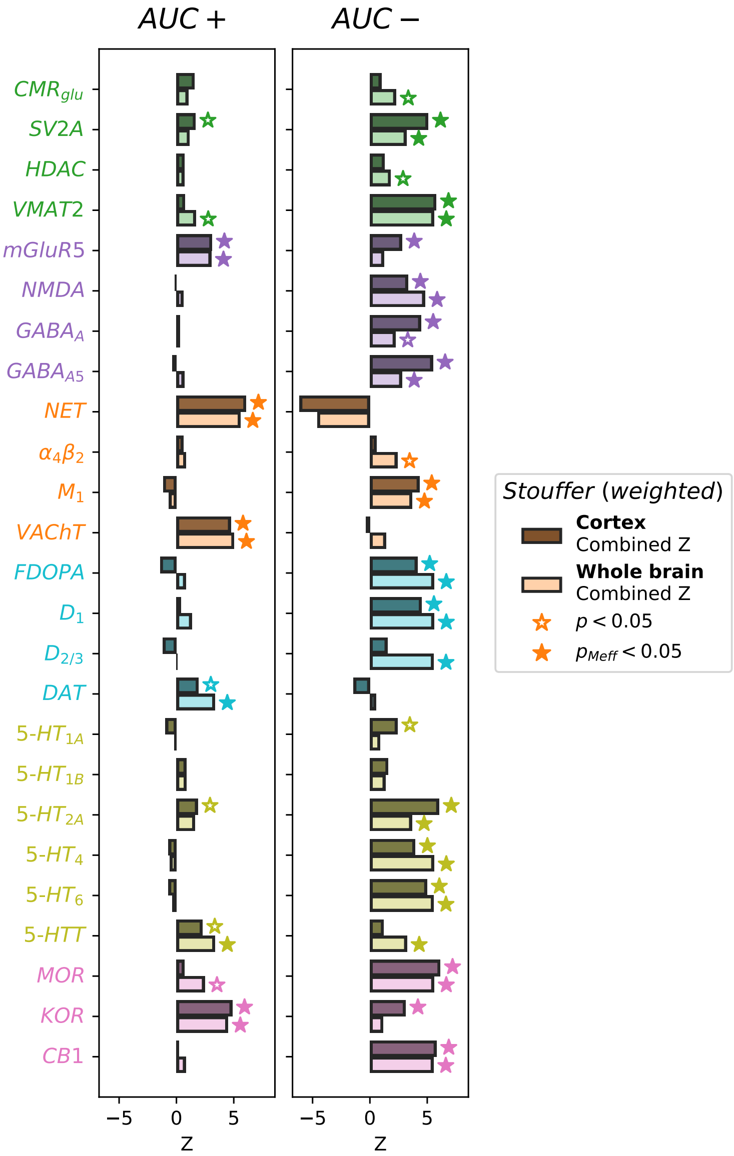
AUC +
AUC −
CMR_glu
SV2A
HDAC
VMAT2
mGluR5
NMDA
GABA_A
GABA_A5
NET
α4β2
M1
VAChT
FDOPA
D1
D2/3
DAT
5-HT1A
5-HT1B
5-HT2A
5-HT4
5-HT6
5-HTT
MOR
KOR
CB1
−5
0
5
Z
Stouffer (weighted)
Cortex
Combined Z
Whole brain
Combined Z
p < 0.05
p_Meff < 0.05

#### **Fig. S5**: Meta-Analysis of AUC p values from six MRI datasets

Meta-analytic Z scores (weighted Stouffer's method) for AUC+ and AUC– across 25 nuclear imaging reference maps, combining raw p values from six independent fMRI cohorts weighted by sample size. Darker bars: Schaefer200; brighter bars: Schaefer200 + subcortex; significance markers indicate pMeff. Abbreviations: AUC: area under the curve; pMeff: p-value corrected for effective number of comparisons; see **Fig. 2** for reference map abbreviations.

**Fig. S6a**: Schaefer100(+Subcortical)

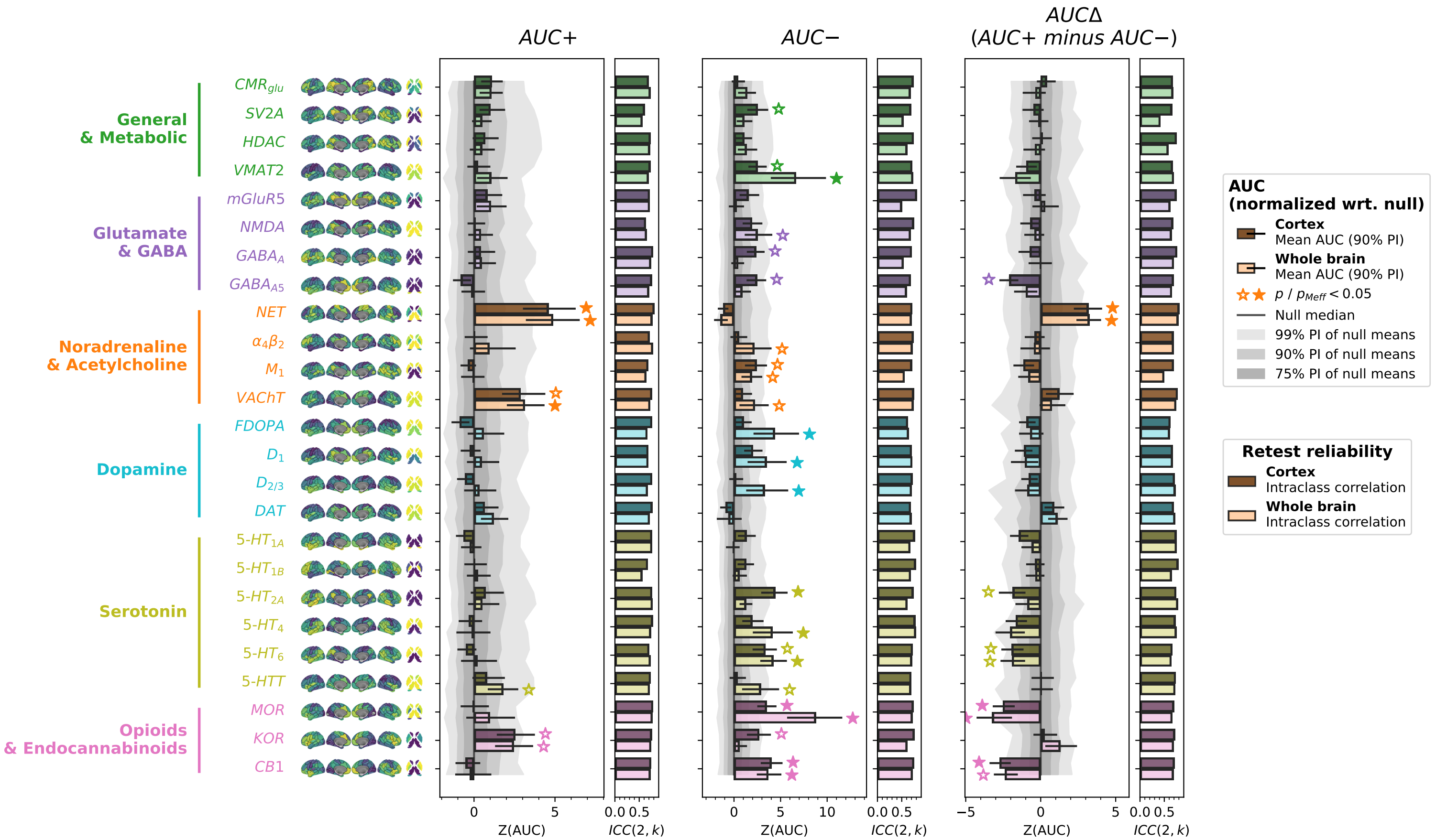

**Fig. S6b**: Schaefer200(+Subcortical)

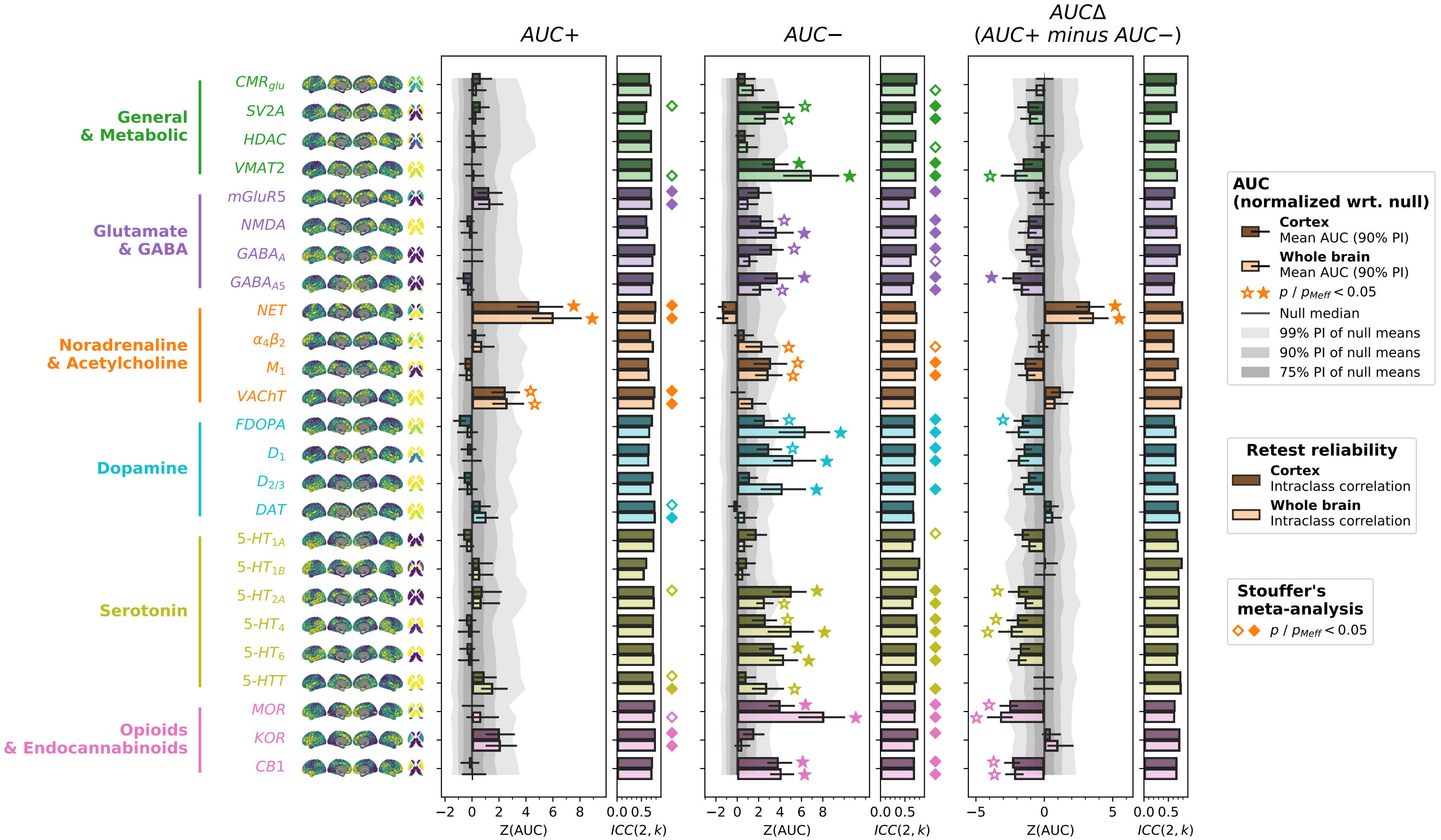

**Fig. S6c**: Schaefer400(+Subcortical)

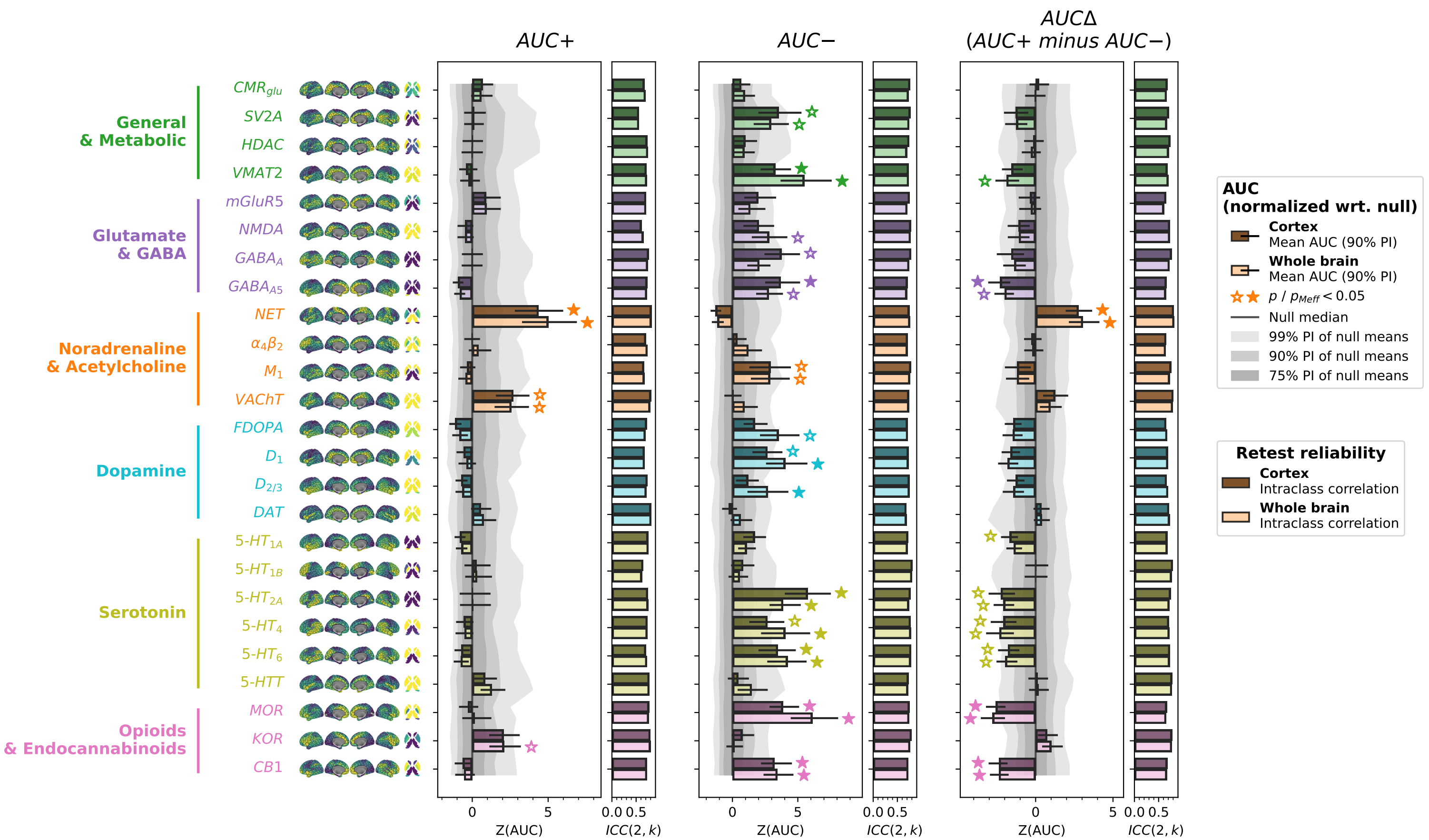

**Fig. S6**: AUC results including AUC-delta for different parcellation resolutions

AUC+, AUC–, and AUCΔ (AUC+ minus AUC–) scores for 25 nuclear imaging reference maps, shown for three parcellation resolutions: **a** Schaefer100 (+subcortex), **b** Schaefer200 (+subcortex), **c** Schaefer400 (+subcortex). Displayed as in **Fig. 2c**. Abbreviations: AUC: area under the curve; PI: percentile interval; ICC: intraclass correlation coefficient; see **Fig. 2** for reference map abbreviations.

**Fig. S7a**: No interhemispheric connections

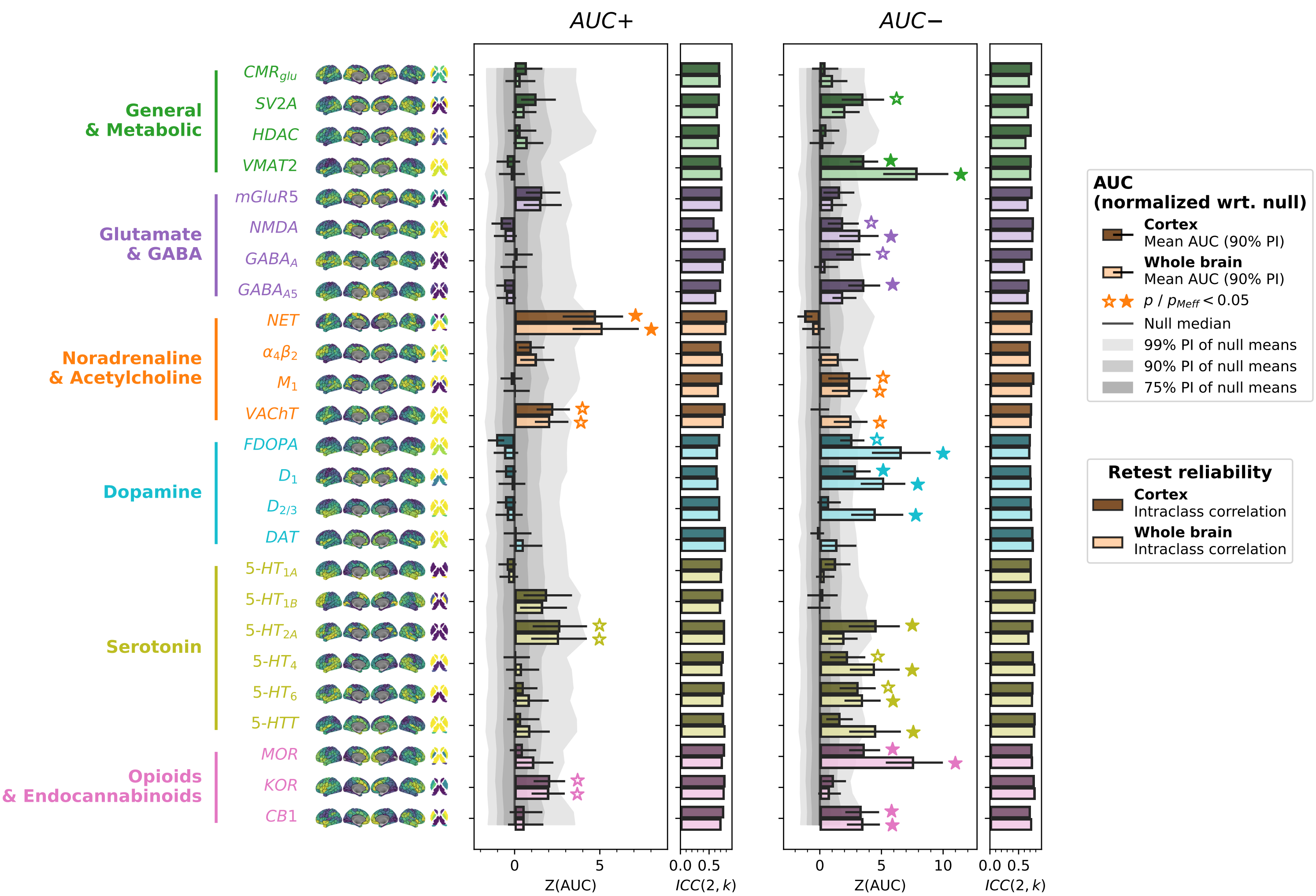

**Fig. S7b**: Polynomial fit aggregation

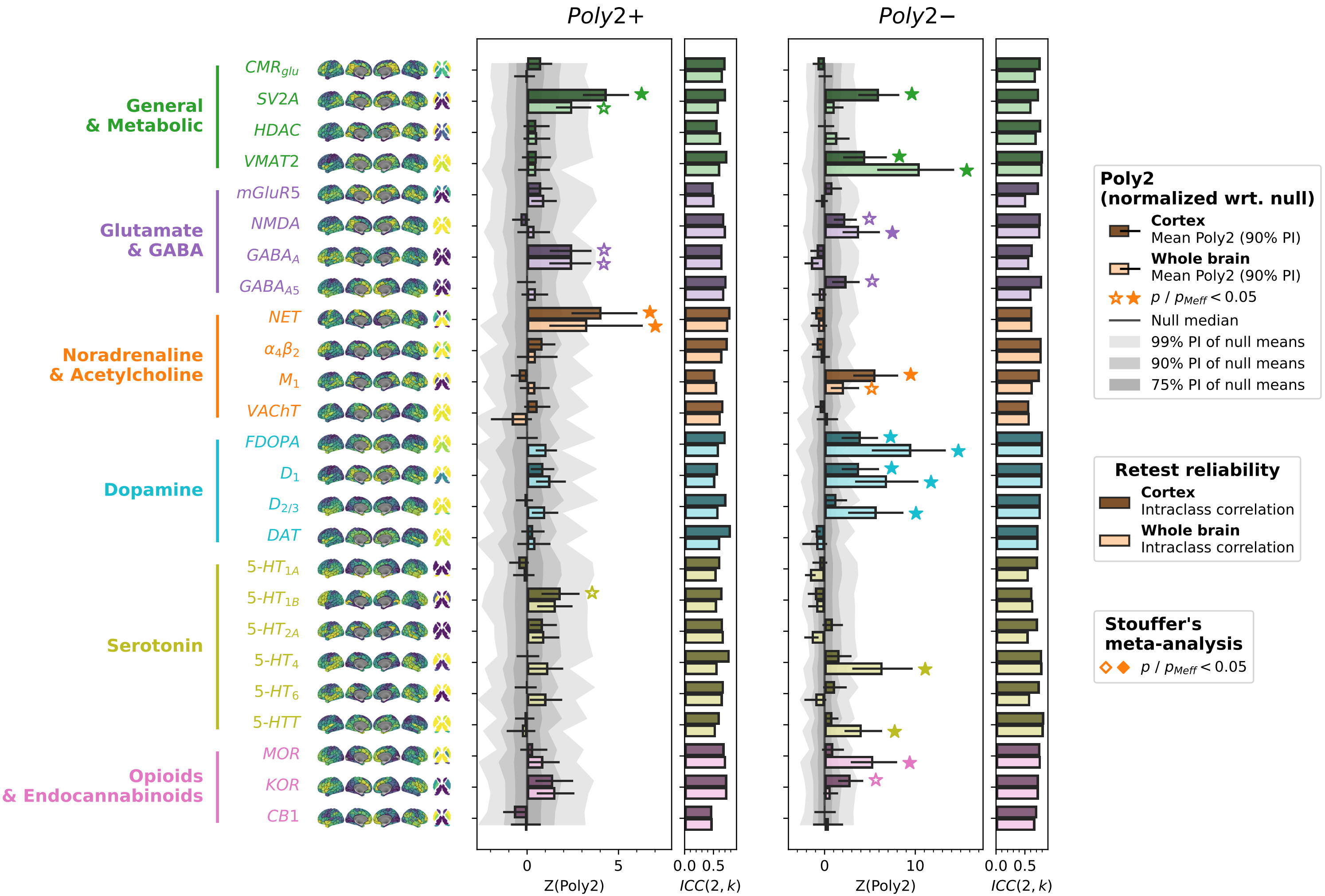

**Fig. S7**: AUC results without interhemispheric connections and an alternative aggregate metric

AUC+ and AUC– scores for 25 nuclear imaging reference maps under two methodological variants (Schaefer200 parcellation). **a** Replication of the main analysis excluding interhemispheric connections. **b** Replication using a 2nd-degree polynomial fit coefficient (Poly2) as an alternative to AUC as the aggregate metric. Displayed as in **Fig. 2c**. Abbreviations: AUC: area under the curve; Poly2: 2nd-degree polynomial fit coefficient; PI: percentile interval; ICC: intraclass correlation coefficient; see **Fig. 2** for reference map abbreviations.

**Fig. S8a**: delta

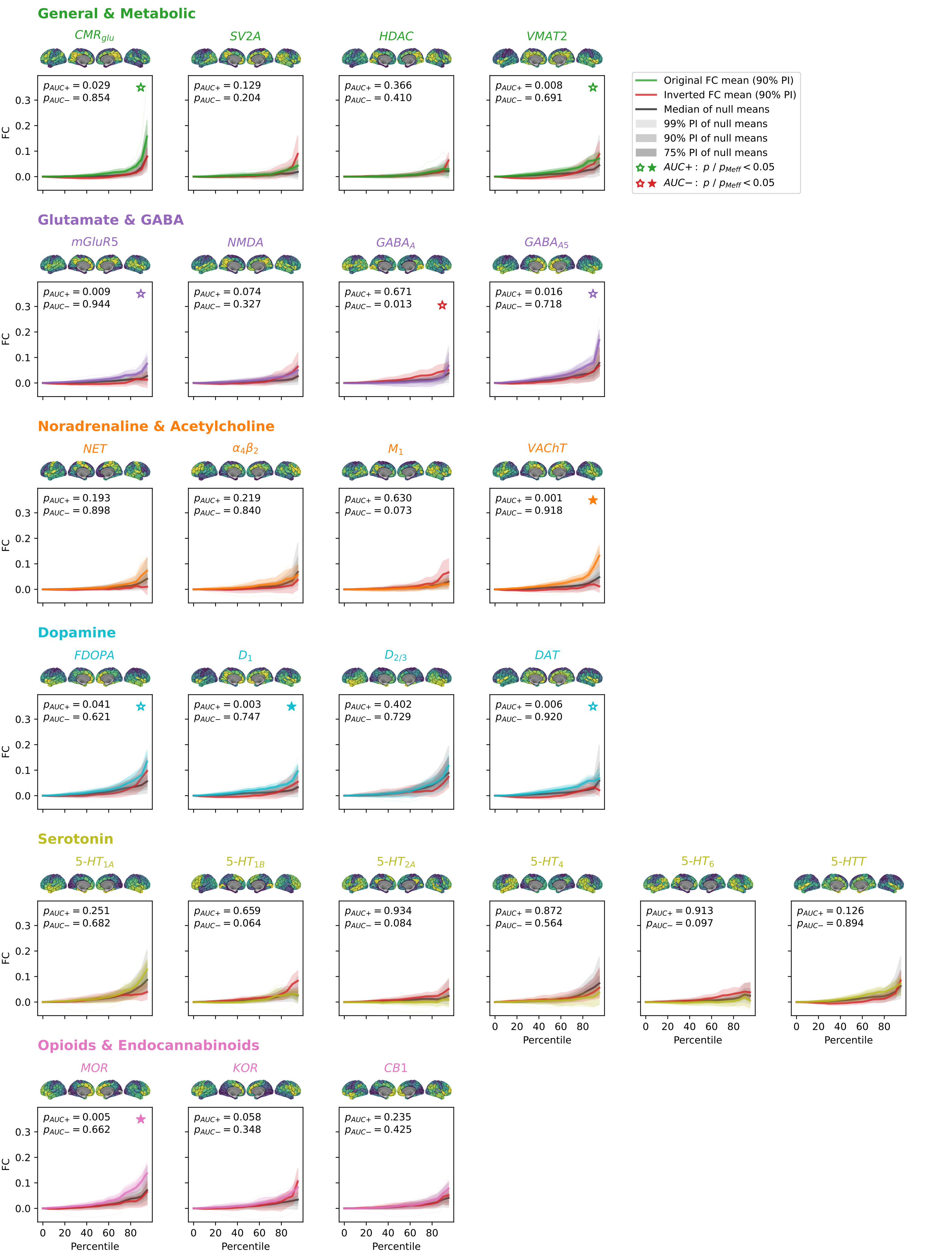

**Fig. S8b**: theta

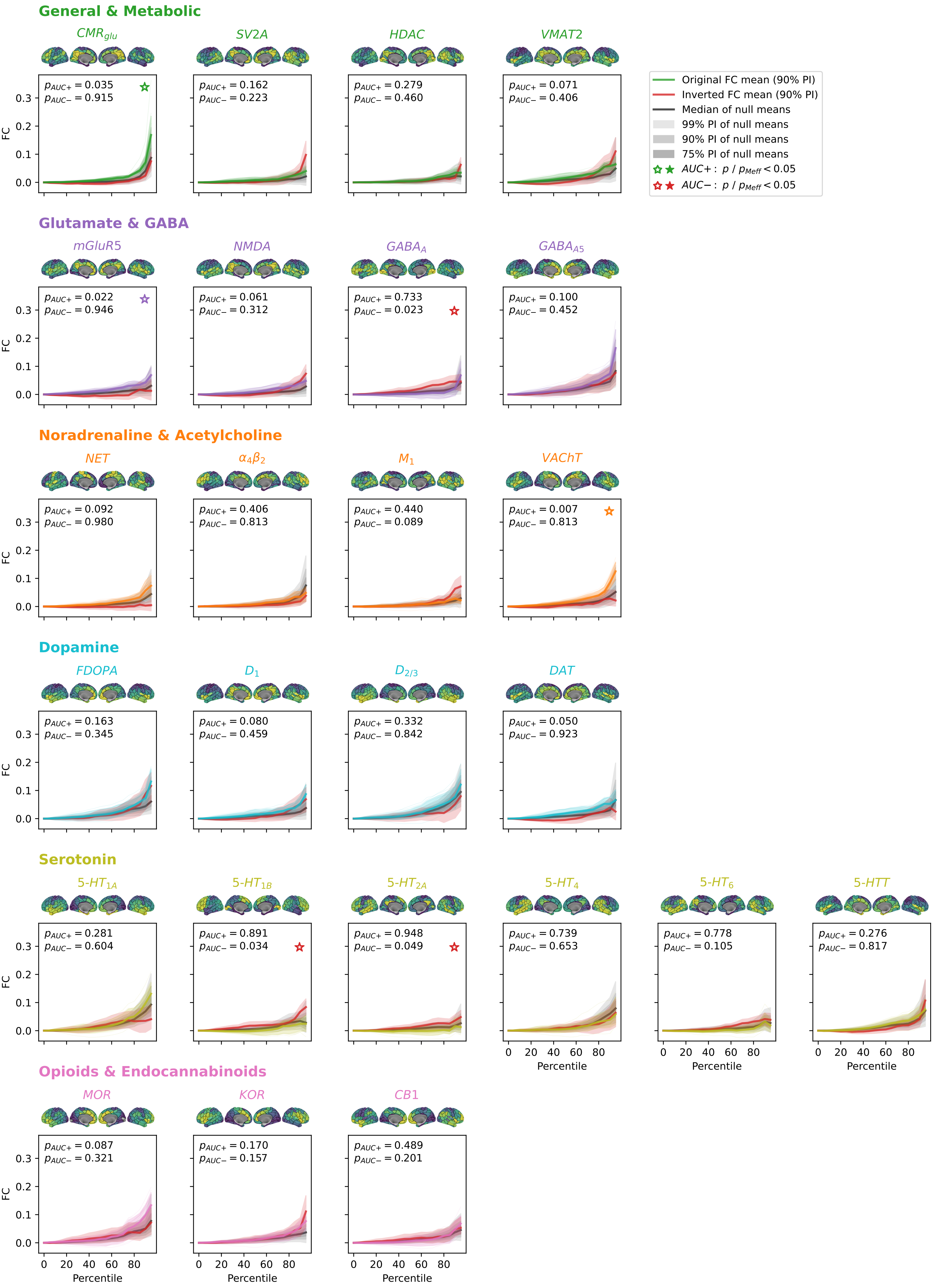

**Fig. S8c**: alpha

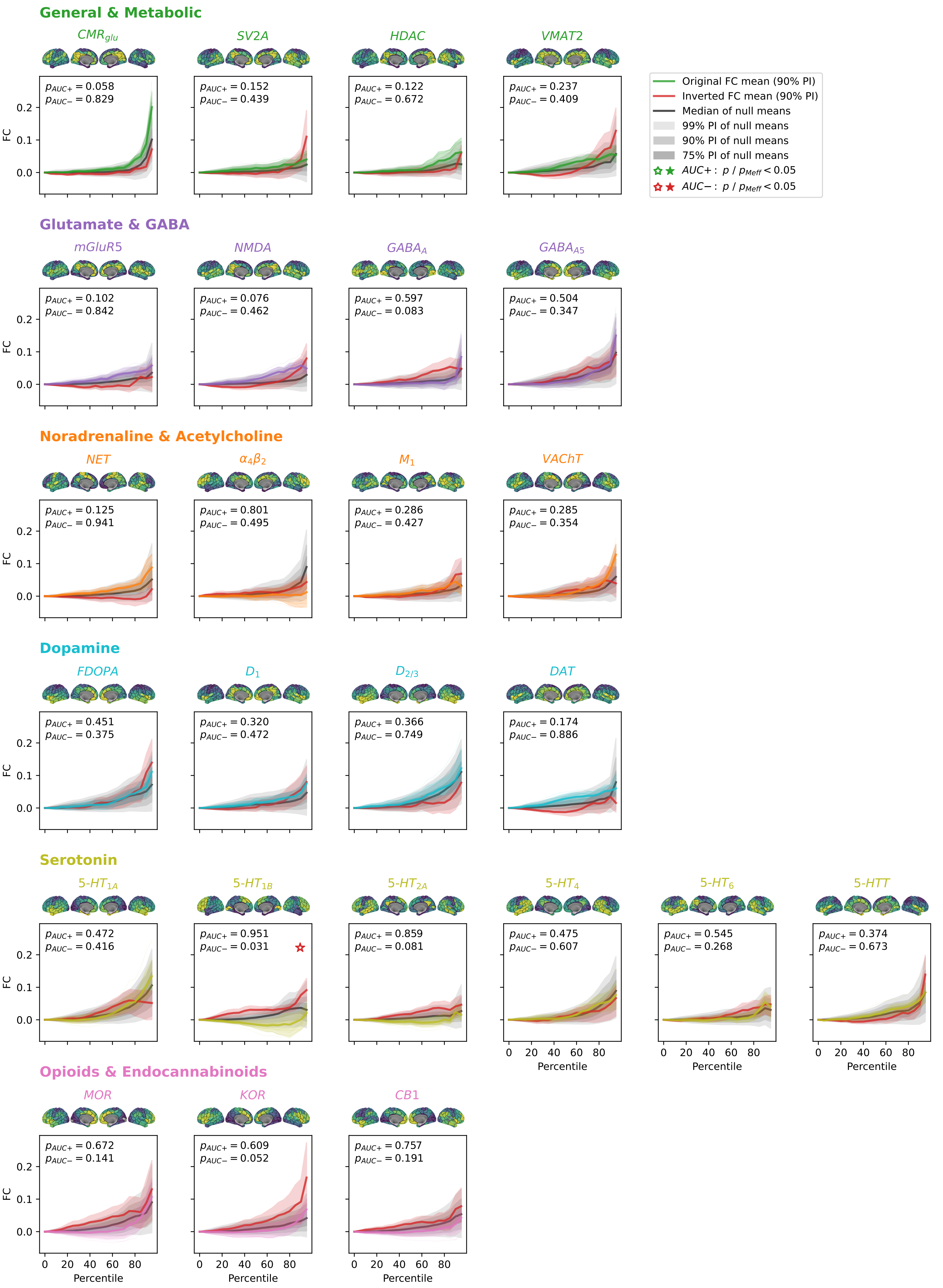

**Fig. S8d**: beta

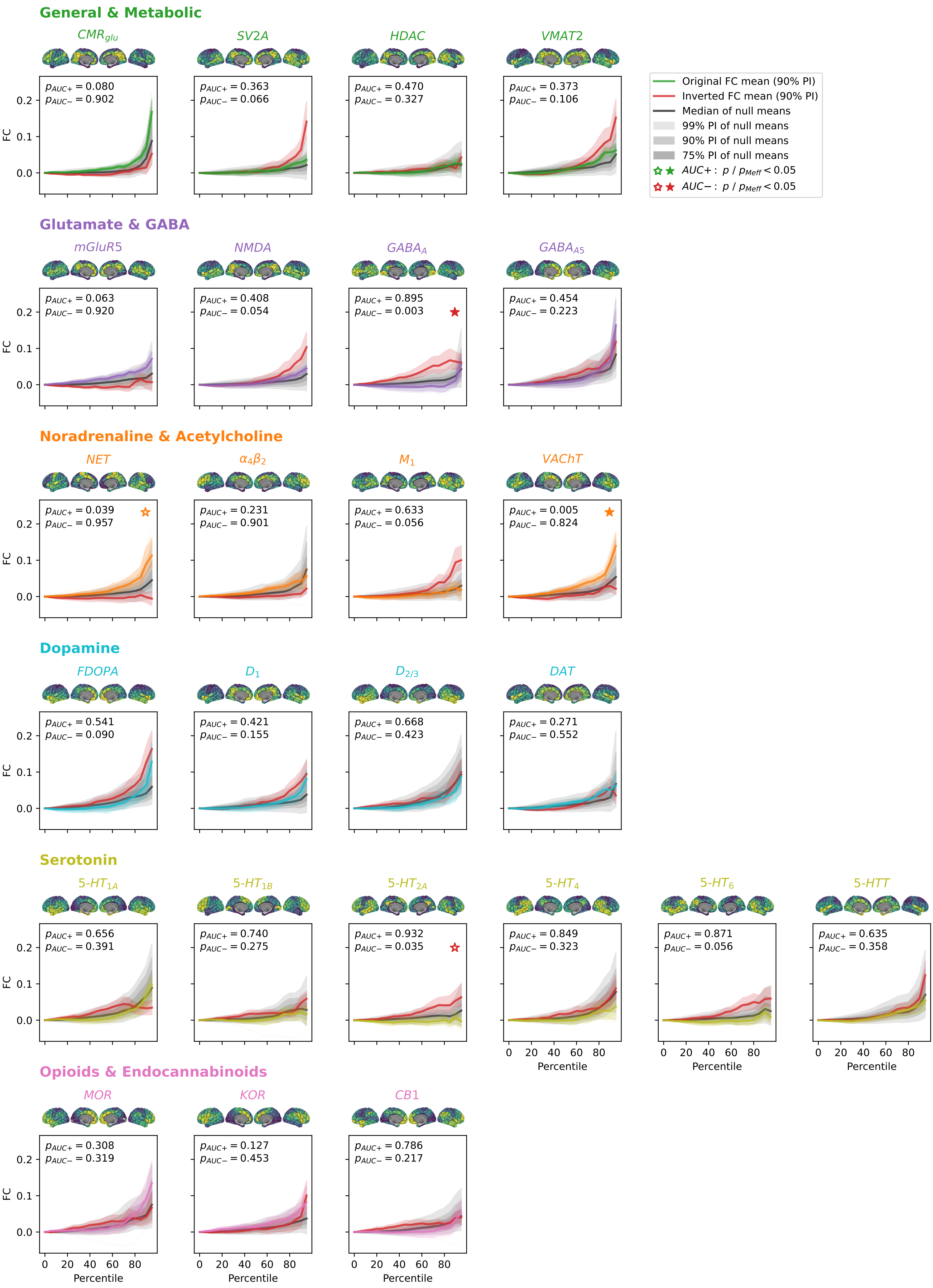

General & Metabolic
$CMR_{glu}$
$p_{AUC+} = 0.080$
$p_{AUC-} = 0.902$
SV2A
$p_{AUC+} = 0.363$
$p_{AUC-} = 0.066$
HDAC
$p_{AUC+} = 0.470$
$p_{AUC-} = 0.327$
VMAT2
$p_{AUC+} = 0.373$
$p_{AUC-} = 0.106$
Original FC mean (90% PI)
Inverted FC mean (90% PI)
Median of null means
99% PI of null means
90% PI of null means
75% PI of null means
$AUC+: p / p_{Meff} < 0.05$
$AUC-: p / p_{Meff} < 0.05$
Glutamate & GABA
mGluR5
$p_{AUC+} = 0.063$
$p_{AUC-} = 0.920$
NMDA
$p_{AUC+} = 0.408$
$p_{AUC-} = 0.054$
$GABA_A$
$p_{AUC+} = 0.895$
$p_{AUC-} = 0.003$
$GABA_{A5}$
$p_{AUC+} = 0.454$
$p_{AUC-} = 0.223$
Noradrenaline & Acetylcholine
NET
$p_{AUC+} = 0.039$
$p_{AUC-} = 0.957$
$\alpha_4\beta_2$
$p_{AUC+} = 0.231$
$p_{AUC-} = 0.901$
$M_1$
$p_{AUC+} = 0.633$
$p_{AUC-} = 0.056$
VAChT
$p_{AUC+} = 0.005$
$p_{AUC-} = 0.824$
Dopamine
FDOPA
$p_{AUC+} = 0.541$
$p_{AUC-} = 0.090$
$D_1$
$p_{AUC+} = 0.421$
$p_{AUC-} = 0.155$
$D_{2/3}$
$p_{AUC+} = 0.668$
$p_{AUC-} = 0.423$
DAT
$p_{AUC+} = 0.271$
$p_{AUC-} = 0.552$
Serotonin
$5\text{-}HT_{1A}$
$p_{AUC+} = 0.656$
$p_{AUC-} = 0.391$
$5\text{-}HT_{1B}$
$p_{AUC+} = 0.740$
$p_{AUC-} = 0.275$
$5\text{-}HT_{2A}$
$p_{AUC+} = 0.932$
$p_{AUC-} = 0.035$
$5\text{-}HT_4$
$p_{AUC+} = 0.849$
$p_{AUC-} = 0.323$
$5\text{-}HT_6$
$p_{AUC+} = 0.871$
$p_{AUC-} = 0.056$
5-HTT
$p_{AUC+} = 0.635$
$p_{AUC-} = 0.358$
Opioids & Endocannabinoids
MOR
$p_{AUC+} = 0.308$
$p_{AUC-} = 0.319$
KOR
$p_{AUC+} = 0.127$
$p_{AUC-} = 0.453$
CB1
$p_{AUC+} = 0.786$
$p_{AUC-} = 0.217$
FC
0.0
0.1
0.2
Percentile
0
20
40
60
80

**Fig. S8e**: lgamma

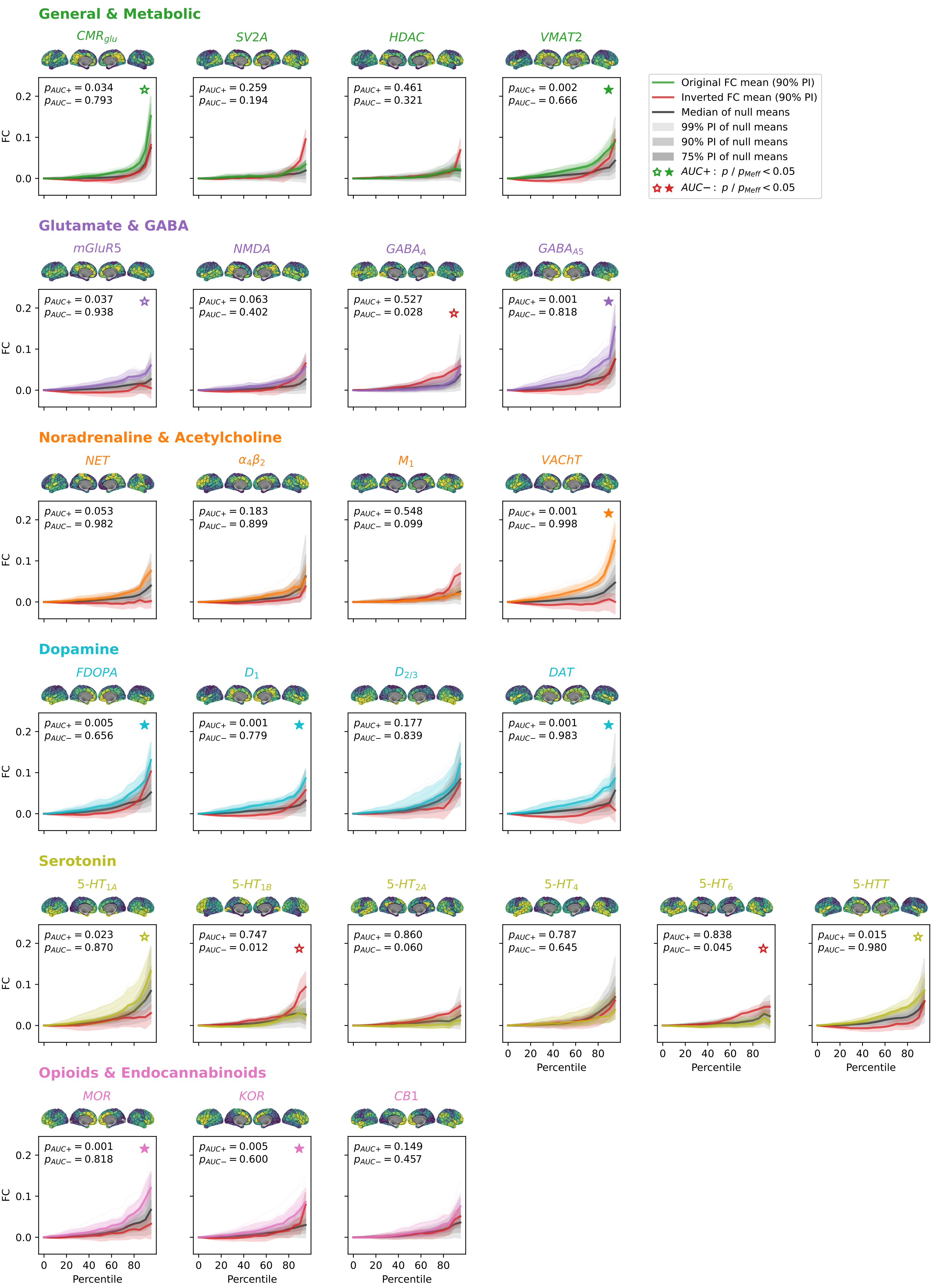

**Fig. S8f**: hgamma

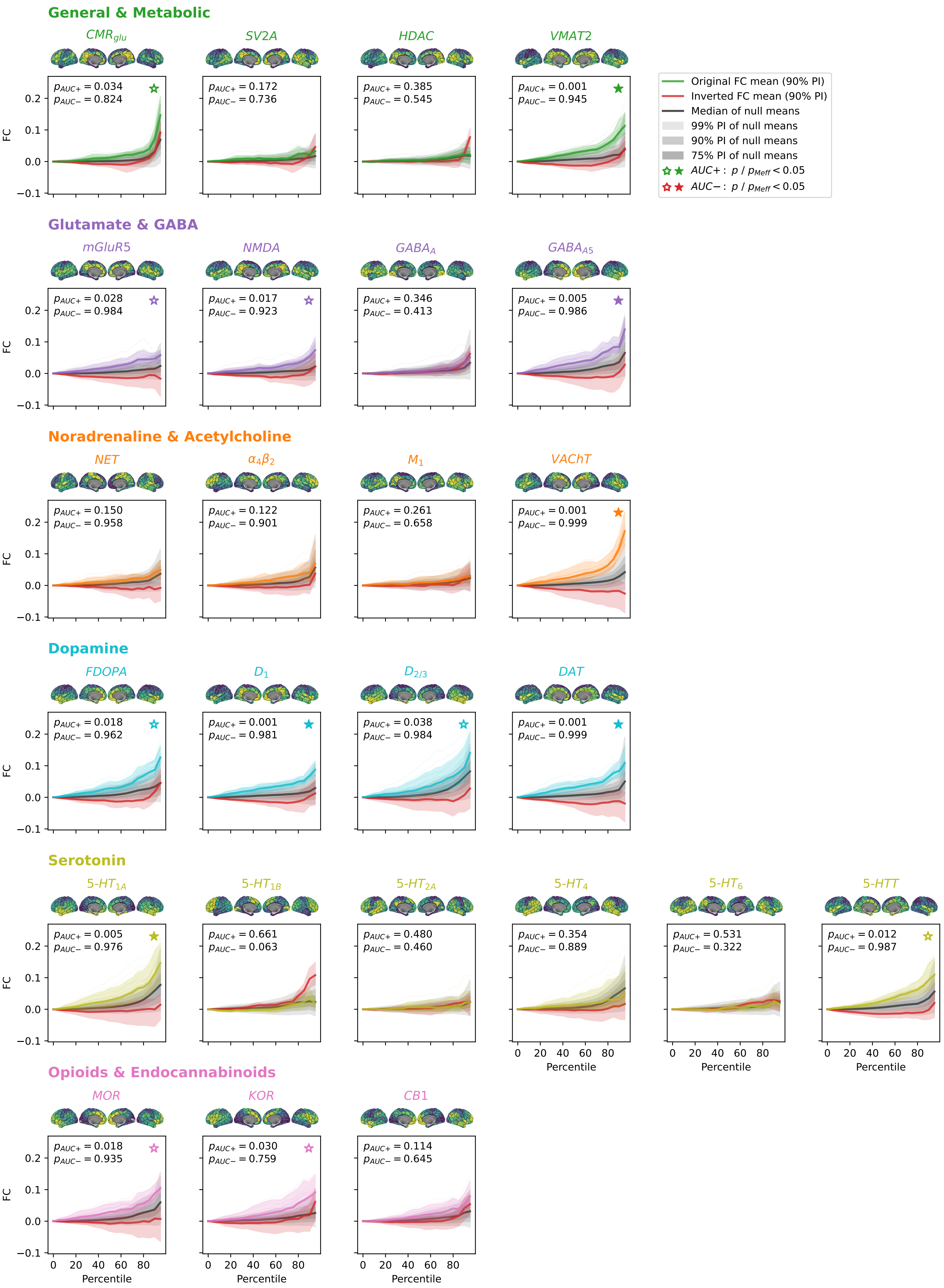

#### **Fig. S8**: NEOFC curves across MEG frequency bands

NEOFC curves for all 25 nuclear imaging reference maps derived from MEG AEC FC matrices, shown for six frequency bands: **a** delta, **b** theta, **c** alpha, **d** beta, **e** low gamma, **f** high gamma. Displayed as in **Fig. S2** (Schaefer200 parcellation). Abbreviations: AEC: amplitude envelope correlation; FC: functional connectivity; PI: percentile interval; see **Fig. 2** for reference map abbreviations.

**Fig. S9a**: Schaefer100

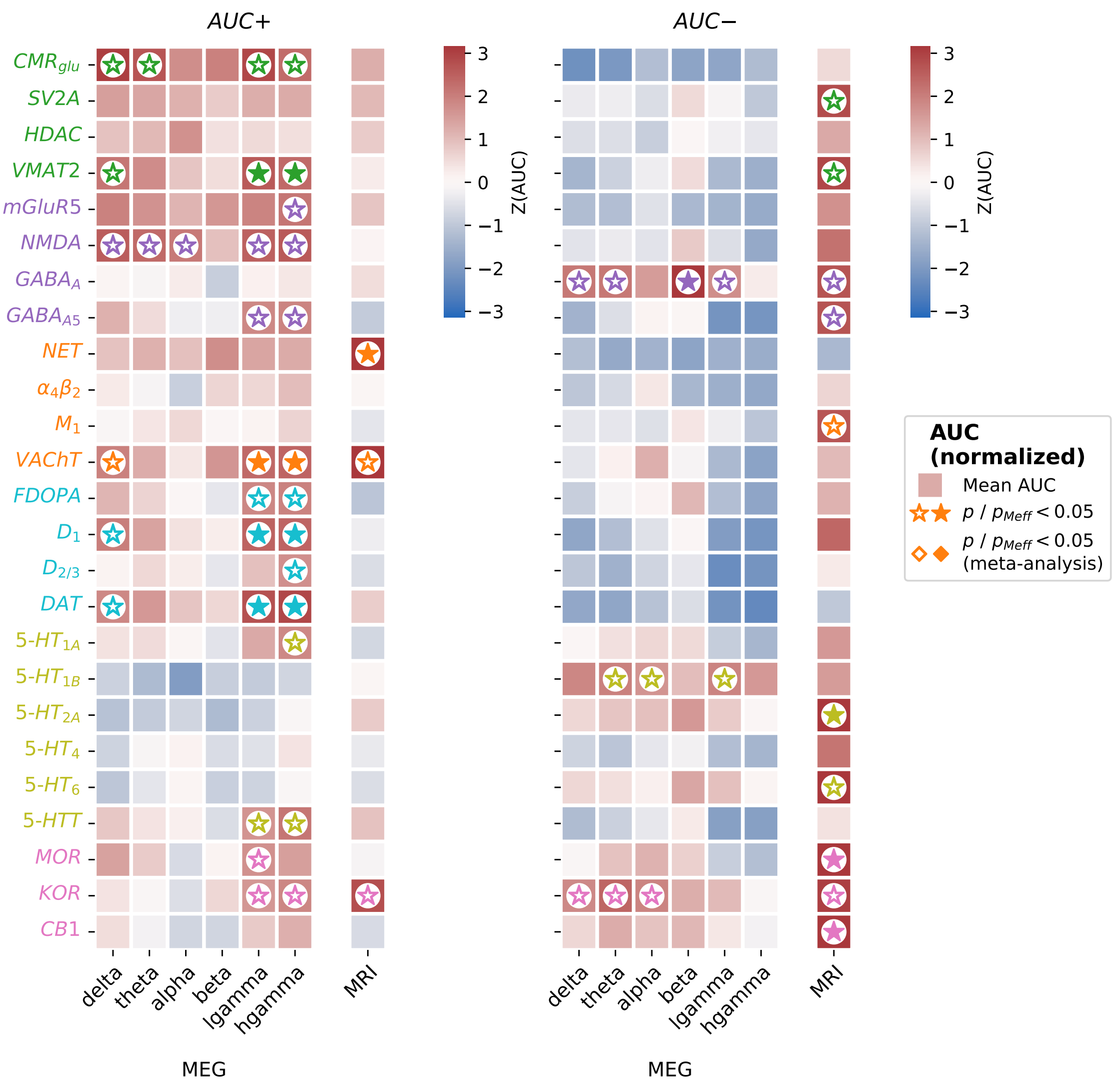

**Fig. S9b**: Schaefer400

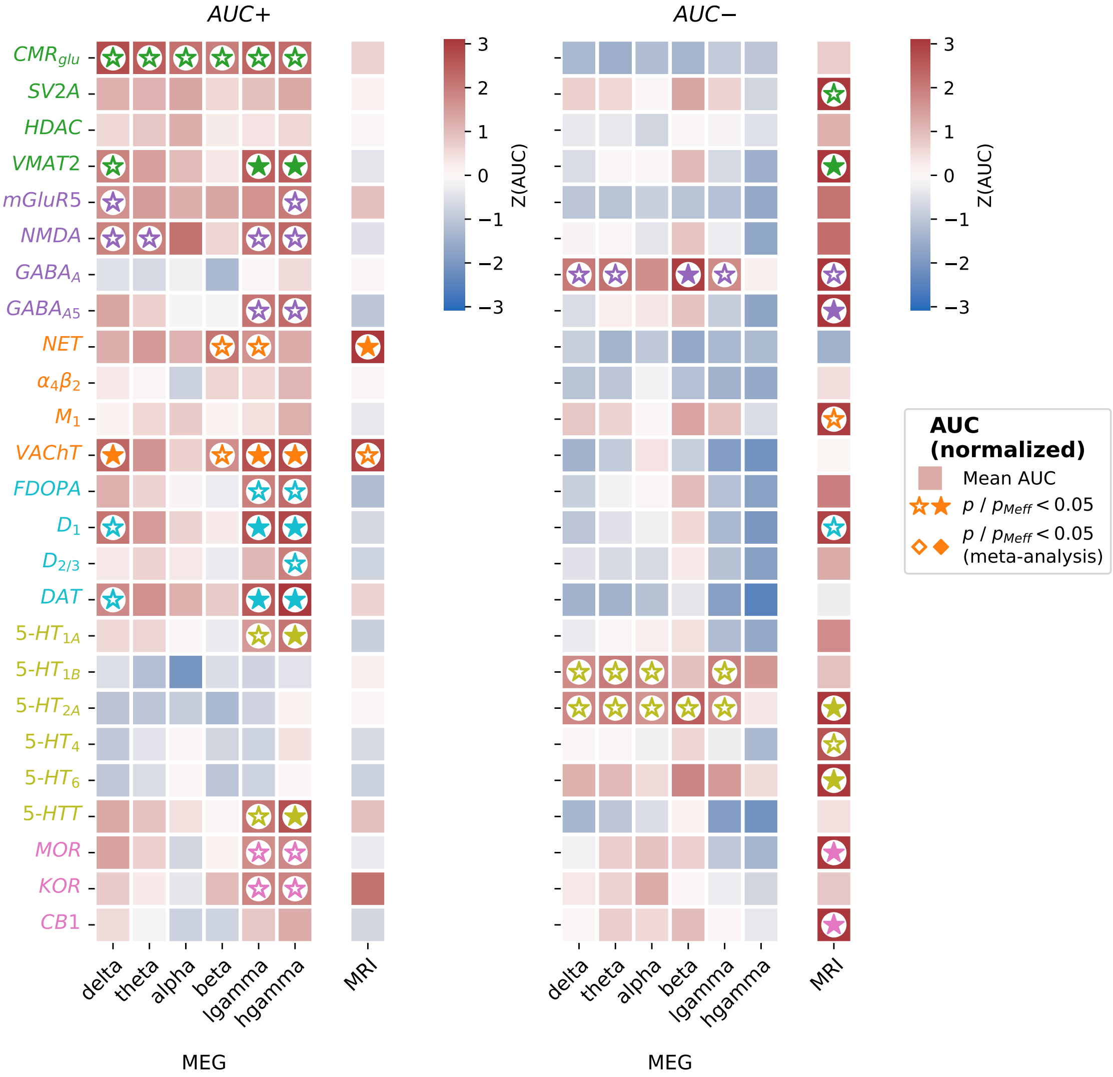

**Fig. S9**: MEG AUC results with different parcellation resolutions

AUC+ and AUC– scores for 25 nuclear imaging reference maps derived from MEG AEC FC matrices, shown for two alternative parcellation resolutions: **a** Schaefer100, **b** Schaefer400. Displayed as in **Fig. 3b**. Abbreviations: AUC: area under the curve; AEC: amplitude envelope correlation; FC: functional connectivity; see **Fig. 2** for reference map abbreviations.

**Fig. S10a**: AUC results

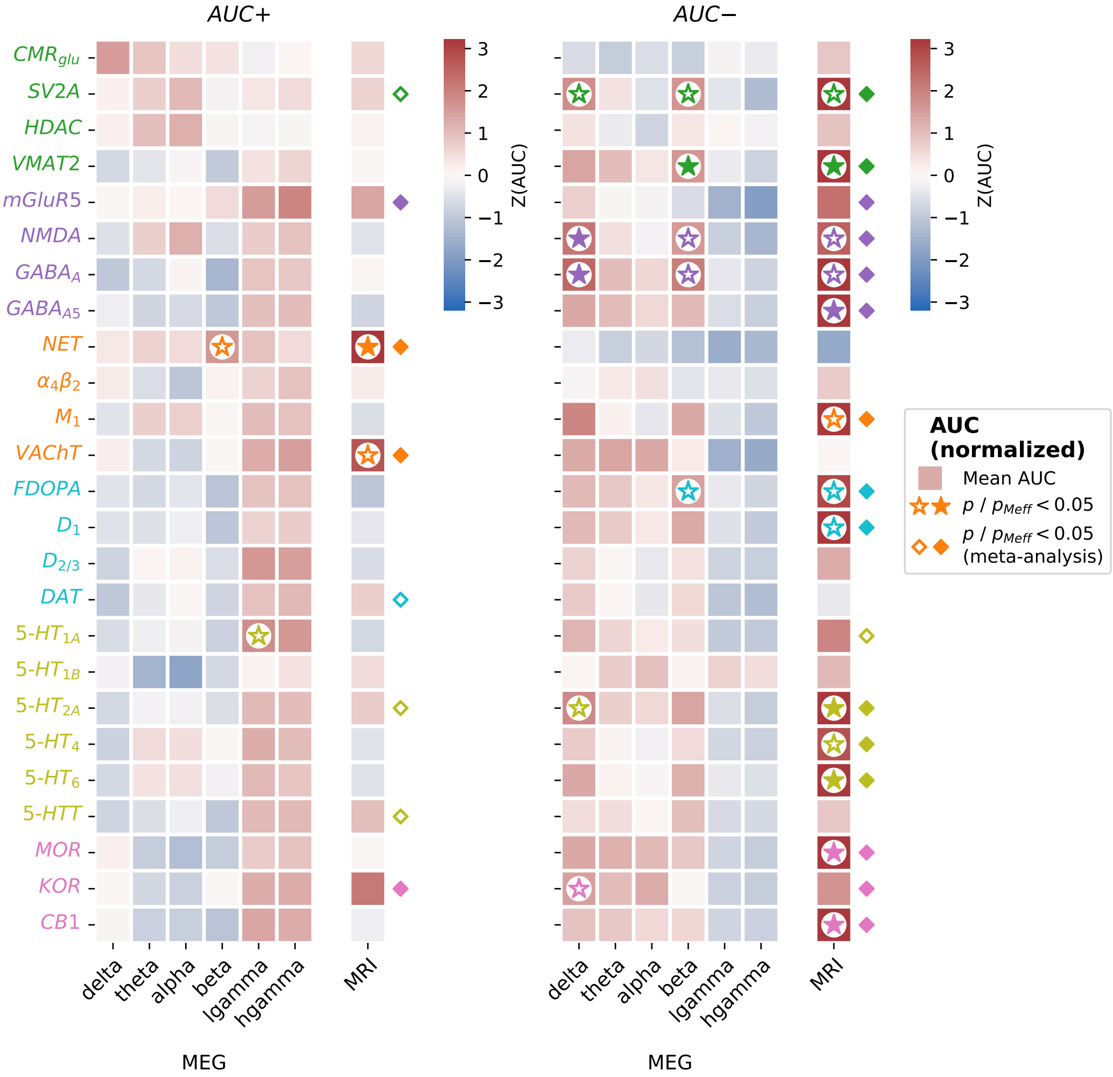

**Fig. S10b**: Correspondence of AUC profiles

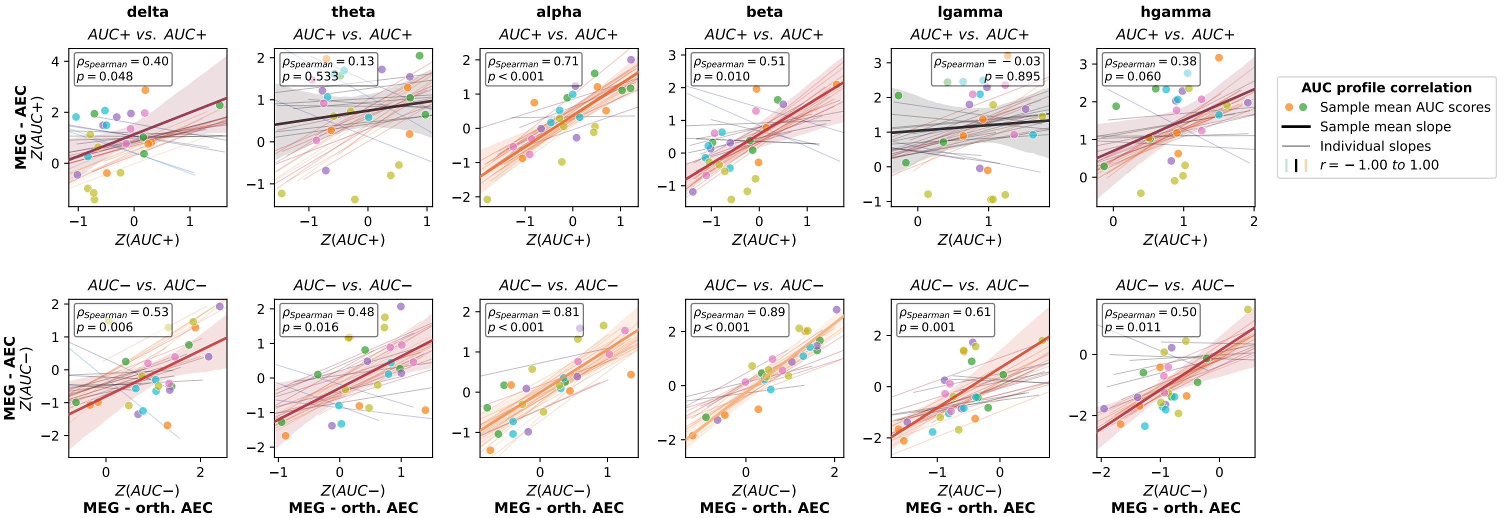

**Fig. S10**: MEG AUC results after orthogonalization

**a** AUC+ and AUC– scores for 25 nuclear imaging reference maps derived from orthogonalized MEG AEC FC matrices (Schaefer200 parcellation), directly comparable to the main analysis in **Fig. 3b**, which used non-orthogonalized AEC matrices. **b** Correspondence between MEG AUC profiles (z-scored against null) calculated with and without orthogonalization. A Spearman correlation was computed across normalized AUC profiles (25 reference maps = 25 observations), separately for AUC+, AUC–, and each frequency band. Compare also to Fig. 3c and S11. Abbreviations: AUC: area under the curve; AEC: amplitude envelope correlation; orth: orthogonalized; FC: functional connectivity; see **Fig. 2** for reference map abbreviations.

**Fig. S11**

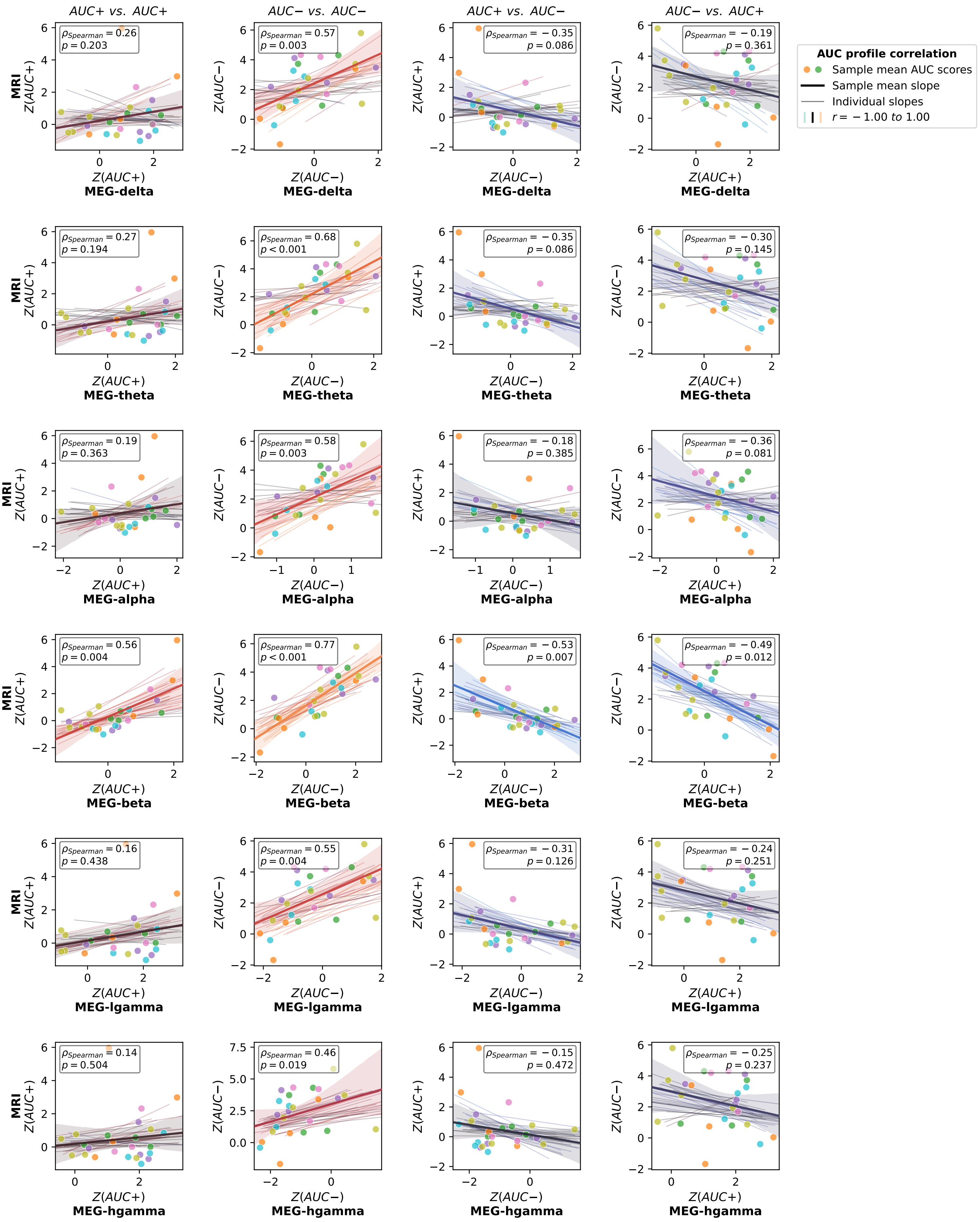
AUC+ vs. AUC+
AUC− vs. AUC−
AUC+ vs. AUC−
AUC− vs. AUC+
AUC profile correlation
Sample mean AUC scores
Sample mean slope
Individual slopes
r = −1.00 to 1.00
MRI
Z(AUC+)
Z(AUC−)
MEG-delta
ρSpearman = 0.26 p = 0.203
ρSpearman = 0.57 p = 0.003
ρSpearman = −0.35 p = 0.086
ρSpearman = −0.19 p = 0.361
MEG-theta
ρSpearman = 0.27 p = 0.194
ρSpearman = 0.68 p < 0.001
ρSpearman = −0.35 p = 0.086
ρSpearman = −0.30 p = 0.145
MEG-alpha
ρSpearman = 0.19 p = 0.363
ρSpearman = 0.58 p = 0.003
ρSpearman = −0.18 p = 0.385
ρSpearman = −0.36 p = 0.081
MEG-beta
ρSpearman = 0.56 p = 0.004
ρSpearman = 0.77 p < 0.001
ρSpearman = −0.53 p = 0.007
ρSpearman = −0.49 p = 0.012
MEG-lgamma
ρSpearman = 0.16 p = 0.438
ρSpearman = 0.55 p = 0.004
ρSpearman = −0.31 p = 0.126
ρSpearman = −0.24 p = 0.251
MEG-hgamma
ρSpearman = 0.14 p = 0.504
ρSpearman = 0.46 p = 0.019
ρSpearman = −0.15 p = 0.472
ρSpearman = −0.25 p = 0.237

#### **Fig. S11**: Correspondence of fMRI and MEG AUC profiles across all frequency bands

Correspondence between fMRI and MEG AUC profiles (z-scored against null) across 25 reference maps, shown for all six MEG frequency bands (rows) and four AUC combinations (columns): fMRI AUC+ vs. MEG AUC+, fMRI AUC– vs. MEG AUC–, fMRI AUC+ vs. MEG AUC–, fMRI AUC– vs. MEG AUC+. Displayed as in **Fig. 3c** (Schaefer200 parcellation). Compare also to Fig. S10b. Abbreviations: AUC: area under the curve; fMRI: functional magnetic resonance imaging; MEG: magnetoencephalography; CI: confidence interval; see **Fig. 2** for reference map abbreviations.

**Fig. S12**

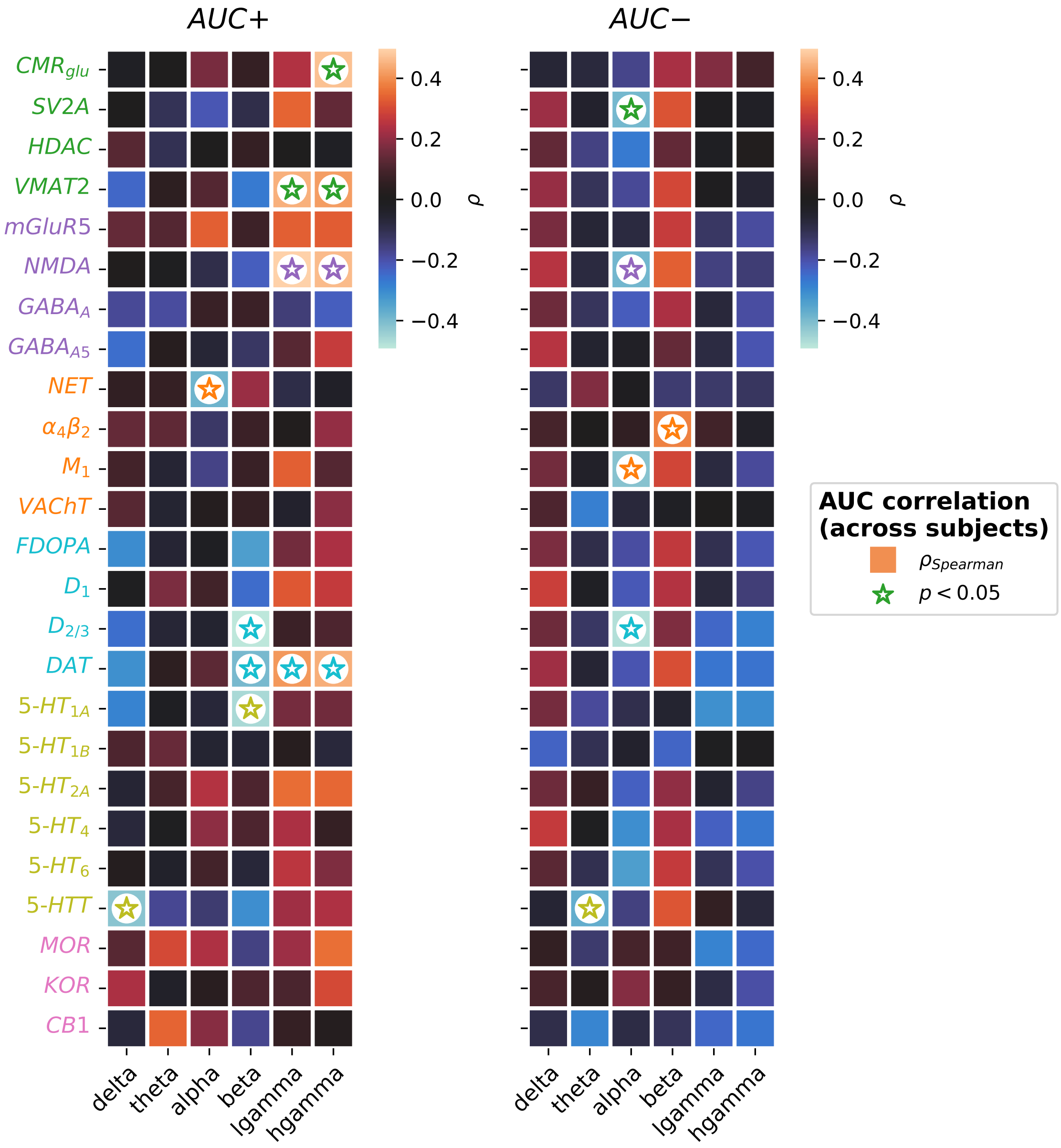

AUC+
AUC−
CMR_glu
SV2A
HDAC
VMAT2
mGluR5
NMDA
GABA_A
GABA_A5
NET
α4β2
M1
VAChT
FDOPA
D1
D2/3
DAT
5-HT1A
5-HT1B
5-HT2A
5-HT4
5-HT6
5-HTT
MOR
KOR
CB1
delta
theta
alpha
beta
lgamma
hgamma
0.4
0.2
0.0
−0.2
−0.4
ρ
AUC correlation (across subjects)
ρ Spearman
p < 0.05

#### **Fig. S12**: Correlation of atlas-wise AUC scores between MRI and MEG

Spearman's rho between individual fMRI and MEG AUC profiles for 25 nuclear imaging reference maps and six MEG frequency bands, shown for AUC+ (left) and AUC– (right) (Schaefer200 parcellation). Color: Spearman's rho; markers indicate $p < 0.05$. Abbreviations: AUC: area under the curve; fMRI: functional magnetic resonance imaging; MEG: magnetoencephalography; see **Fig. 2** for reference map abbreviations.

**Fig. S13a**: AUC+

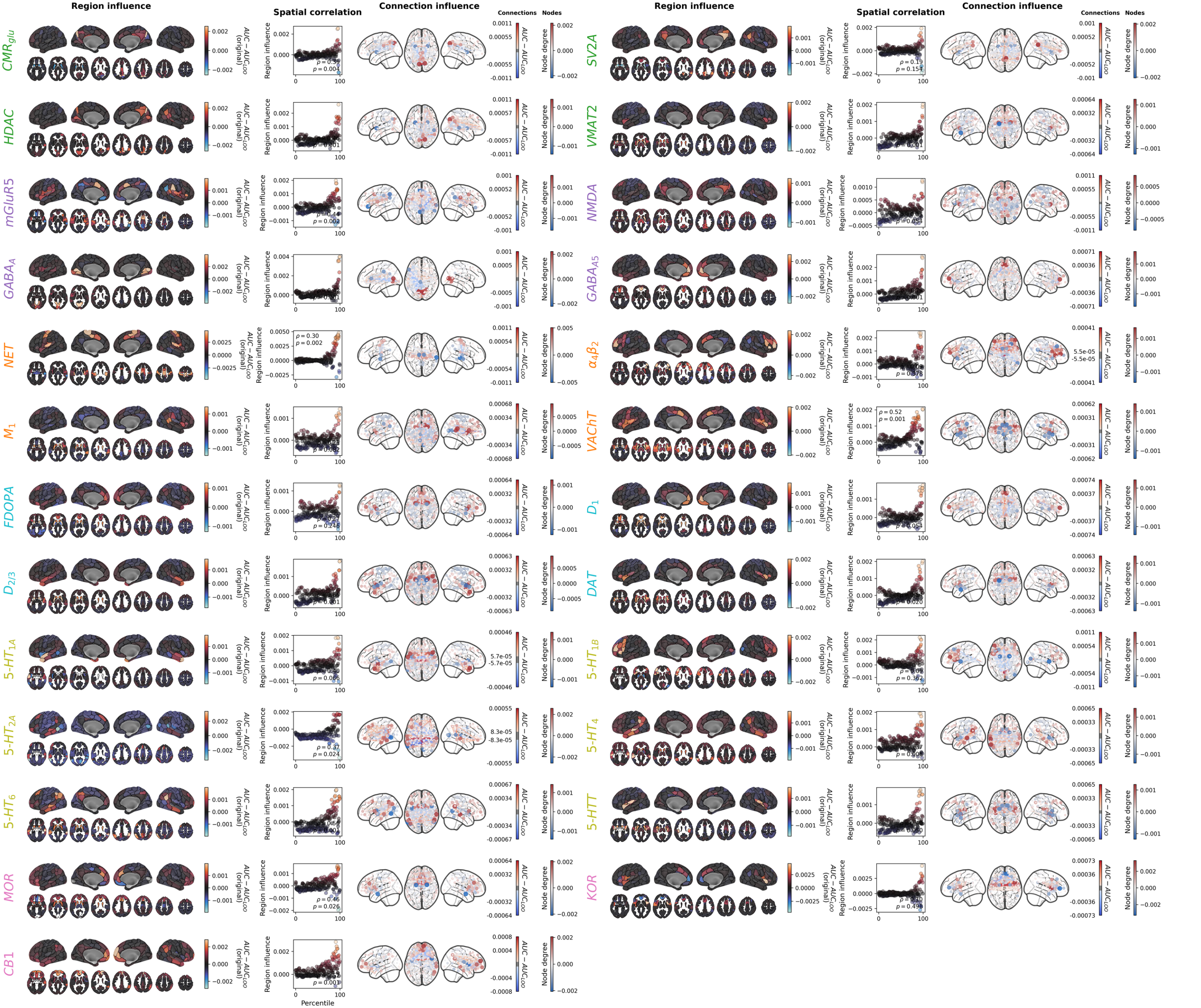
Region influence
Spatial correlation
Connection influence
Connections
Nodes
Region influence
Spatial correlation
Connection influence
Connections
Nodes
CMRglu
SV2A
HDAC
VMAT2
mGluR5
NMDA
GABAA
GABAA5
NET
α4β2
M1
VAChT
FDOPA
D1
D2/3
DAT
5-HT1A
5-HT1B
5-HT2A
5-HT4
5-HT6
5-HTT
MOR
KOR
CB1
AUC − AUCLOO (original)
Region influence
AUC − AUCLOO
Node degree
Percentile

**Fig. S13b**: AUC–

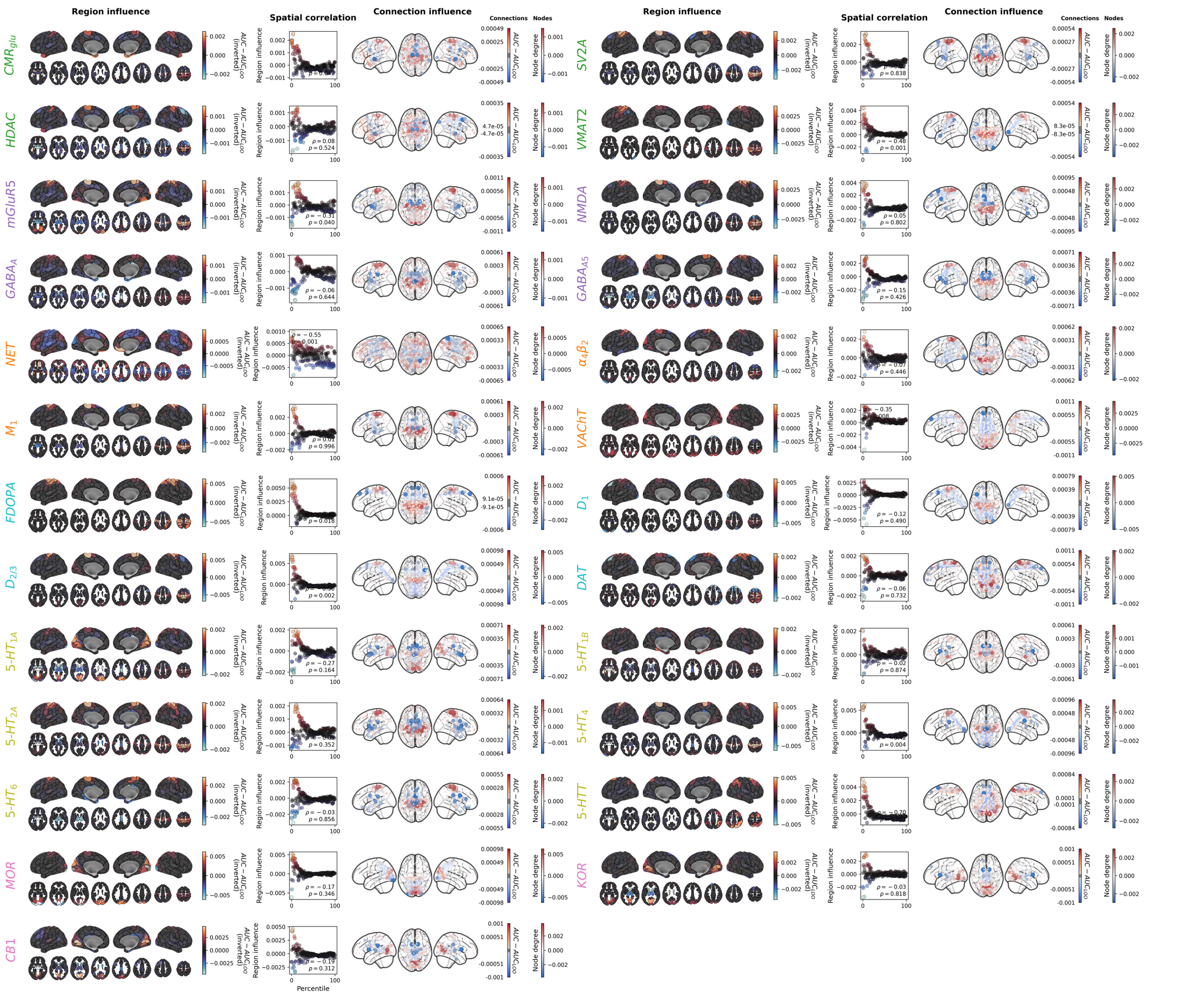
Region influence
Spatial correlation
Connection influence
Connections
Nodes
Region influence
Spatial correlation
Connection influence
Connections
Nodes
CMRglu
HDAC
mGluR5
GABAA
NET
M1
FDOPA
D2/3
5-HT1A
5-HT2A
5-HT6
MOR
CB1
SV2A
VMAT2
NMDA
GABAA5
α4β2
VAChT
D1
DAT
5-HT1B
5-HT4
5-HTT
KOR
AUC − AUCLOO (inverted)
Region influence
Node degree
Percentile

**Fig. S13**: Leave-one-out analysis to identify influential regions and connections

Regional (left, leave-one-region-out) and connection (right, leave-one-connection-out) influence on AUC for all 25 nuclear imaging reference maps (Schaefer200 + subcortex parcellation). **a** AUC+ maps; **b** AUC– maps. Color indicates direction and magnitude of influence. Displayed as in **Fig. 4a**. Abbreviations: AUC: area under the curve; see **Fig. 2** for reference map abbreviations.

**Fig. S14a**: Schaefer200

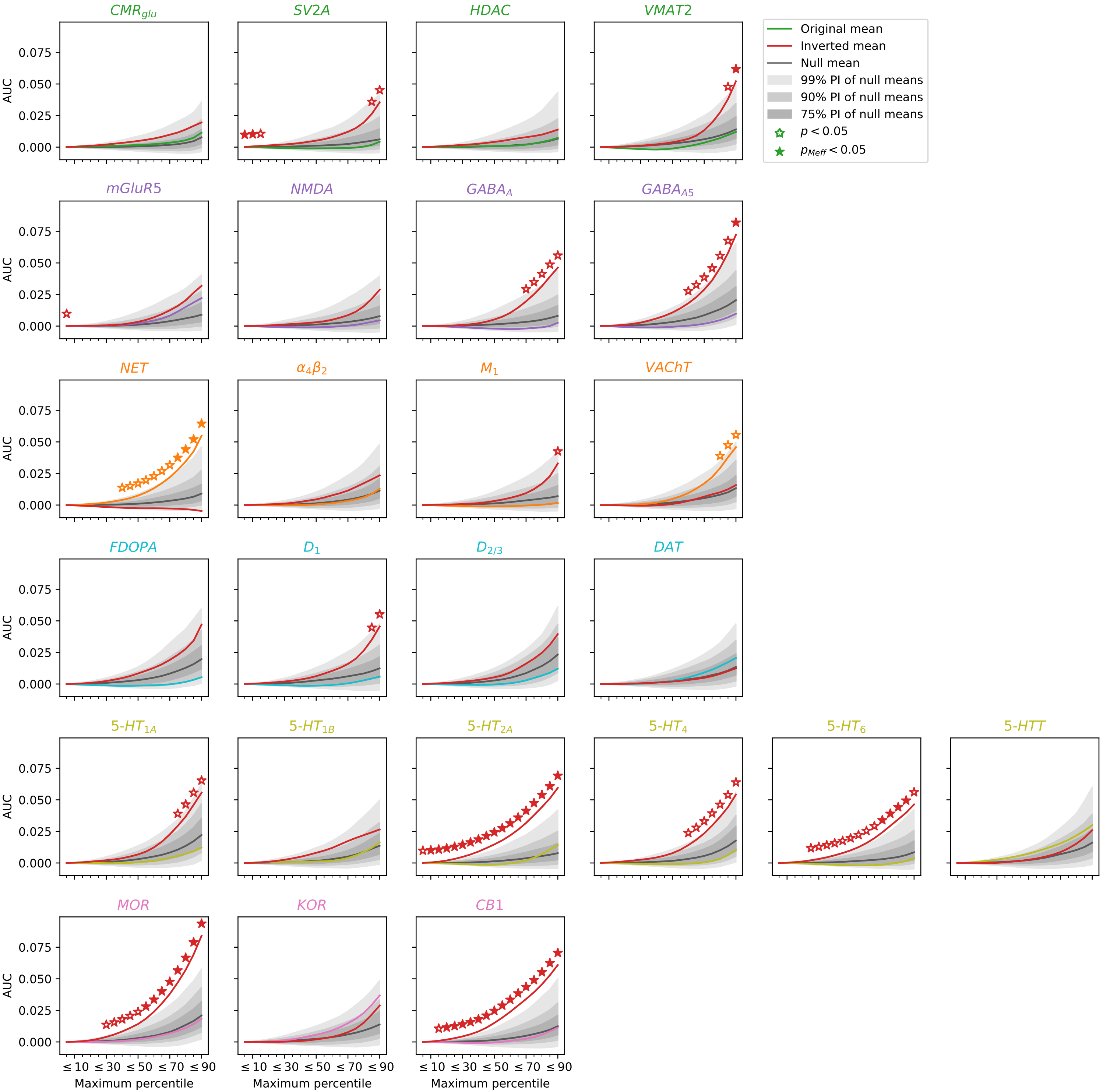

**Fig. S14b**: Schaefer200+Subcortical

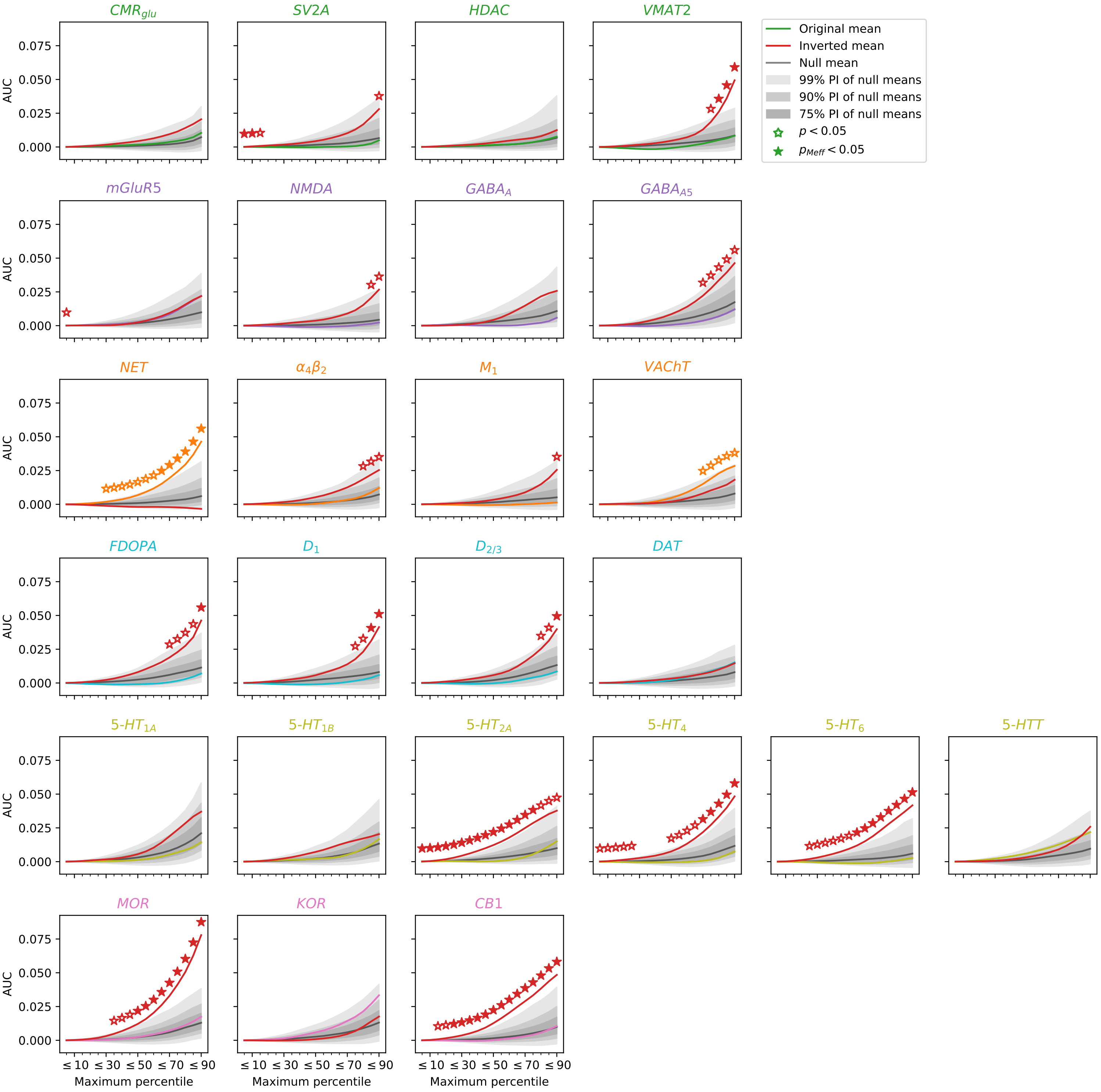

**Fig. S14**: NEOFC curves with AUC estimation restricted to lowered percentiles

Sensitivity analysis assessing whether AUC effects are driven by the highest-density atlas regions. The maximum percentile threshold included in AUC calculation was systematically varied from 5 to 90, and significance reported as a function of this threshold for all 25 nuclear imaging reference maps. **a** Schaefer200 parcellation; **b** Schaefer200 + subcortex parcellation. Abbreviations: AUC: area under the curve; pMeff: p-value corrected for effective number of comparisons; see **Fig. 2** for reference map abbreviations.

**Fig. S15**

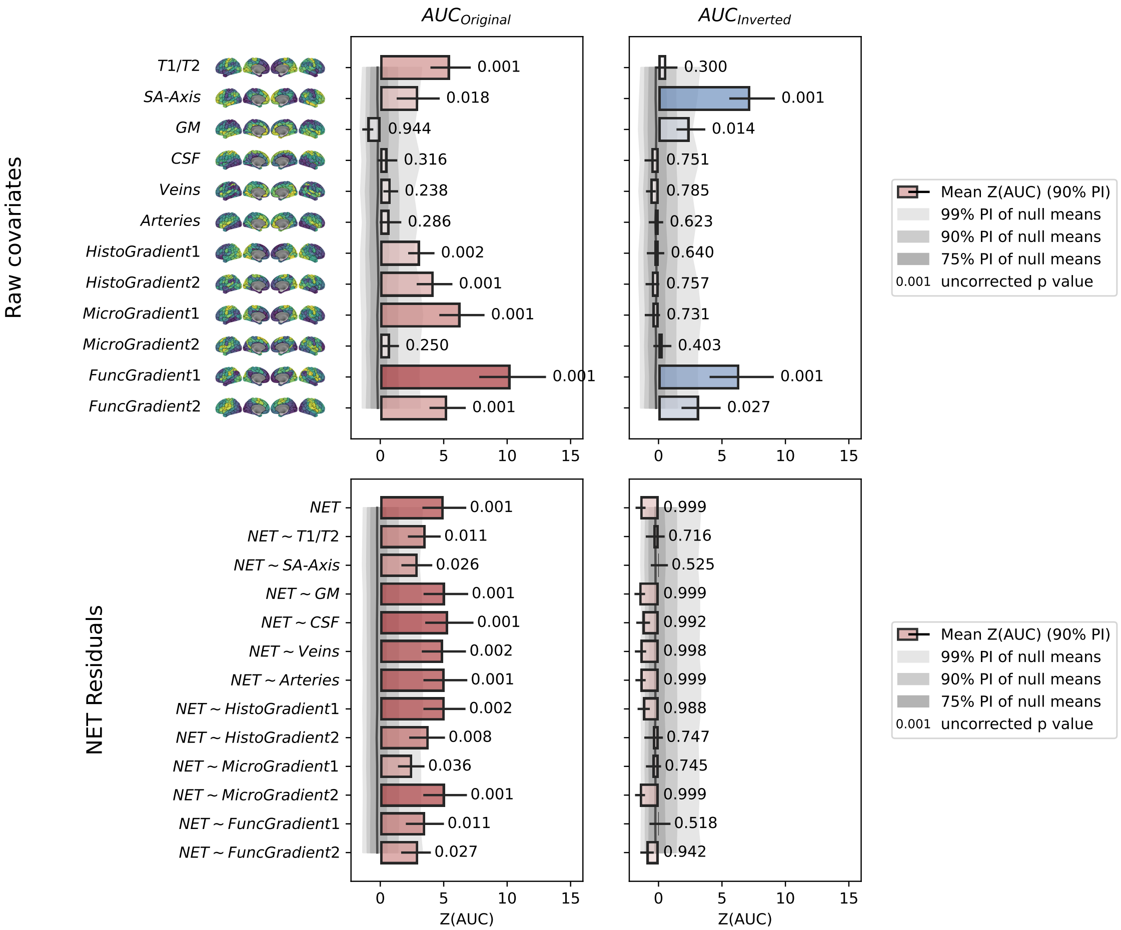

$AUC_{Original}$
$AUC_{Inverted}$
Raw covariates
T1/T2
SA-Axis
GM
CSF
Veins
Arteries
HistoGradient1
HistoGradient2
MicroGradient1
MicroGradient2
FuncGradient1
FuncGradient2
0.001
0.018
0.944
0.316
0.238
0.286
0.002
0.001
0.001
0.250
0.001
0.001
0.300
0.001
0.014
0.751
0.785
0.623
0.640
0.757
0.731
0.403
0.001
0.027
NET Residuals
NET
NET ~ T1/T2
NET ~ SA-Axis
NET ~ GM
NET ~ CSF
NET ~ Veins
NET ~ Arteries
NET ~ HistoGradient1
NET ~ HistoGradient2
NET ~ MicroGradient1
NET ~ MicroGradient2
NET ~ FuncGradient1
NET ~ FuncGradient2
0.001
0.011
0.026
0.001
0.001
0.002
0.001
0.002
0.008
0.036
0.001
0.011
0.027
0.999
0.716
0.525
0.999
0.992
0.998
0.999
0.988
0.747
0.745
0.999
0.518
0.942
0
5
10
15
Z(AUC)
Mean Z(AUC) (90% PI)
99% PI of null means
90% PI of null means
75% PI of null means
0.001 uncorrected p value

**Fig. S15**: Effects of covariate regression on NET AUC results

Sensitivity analysis testing whether the NET AUC+ effect is driven by spatially confounding factors. Each spatial covariate map was individually regressed from the NET atlas and NEOFC AUC+ estimation was rerun on the residuals (Schaefer200 parcellation). Covariate maps included T1/T2 ratio, sensory-association axis, BigBrain gradient maps, gray matter and cerebrospinal fluid probability maps, and probability maps of cerebral veins and arteries. AUC+ results after regression are shown in comparison to the unregressed baseline. Displayed as in **Fig. 2c**. Abbreviations: AUC: area under the curve; PI: percentile interval; NET: norepinephrine/noradrenaline transporter; SA: sensory-association; GM: gray matter; CSF: cerebrospinal fluid.

**Fig. S16**

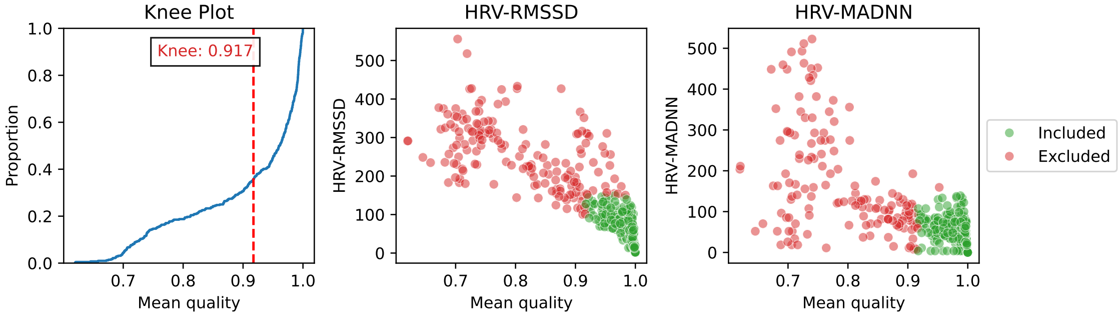
Knee Plot
Knee: 0.917
Proportion
1.0
0.8
0.6
0.4
0.2
0.0
0.7
0.8
0.9
1.0
Mean quality
HRV-RMSSD
500
400
300
200
100
0
HRV-MADNN
Included
Excluded

**Fig. S16**: Quality control of HCP-YA physiological data (heart-rate variability)

Quality control of HRV estimates derived from PPG recordings (HCP-YA dataset). Left: empirical cumulative distribution of mean PPG signal quality index; red dashed line indicates the data-driven knee-point threshold used for session exclusion. Center and right: RMSSD and MadNN as a function of mean PPG quality index; color indicates included (green) and excluded (red) sessions based on quality threshold and upper HRV bound (150 ms). Abbreviations: HRV: heart rate variability; PPG: photoplethysmography; RMSSD: root mean square of successive differences; MadNN: median absolute deviation of NN intervals.

### Fig. S17

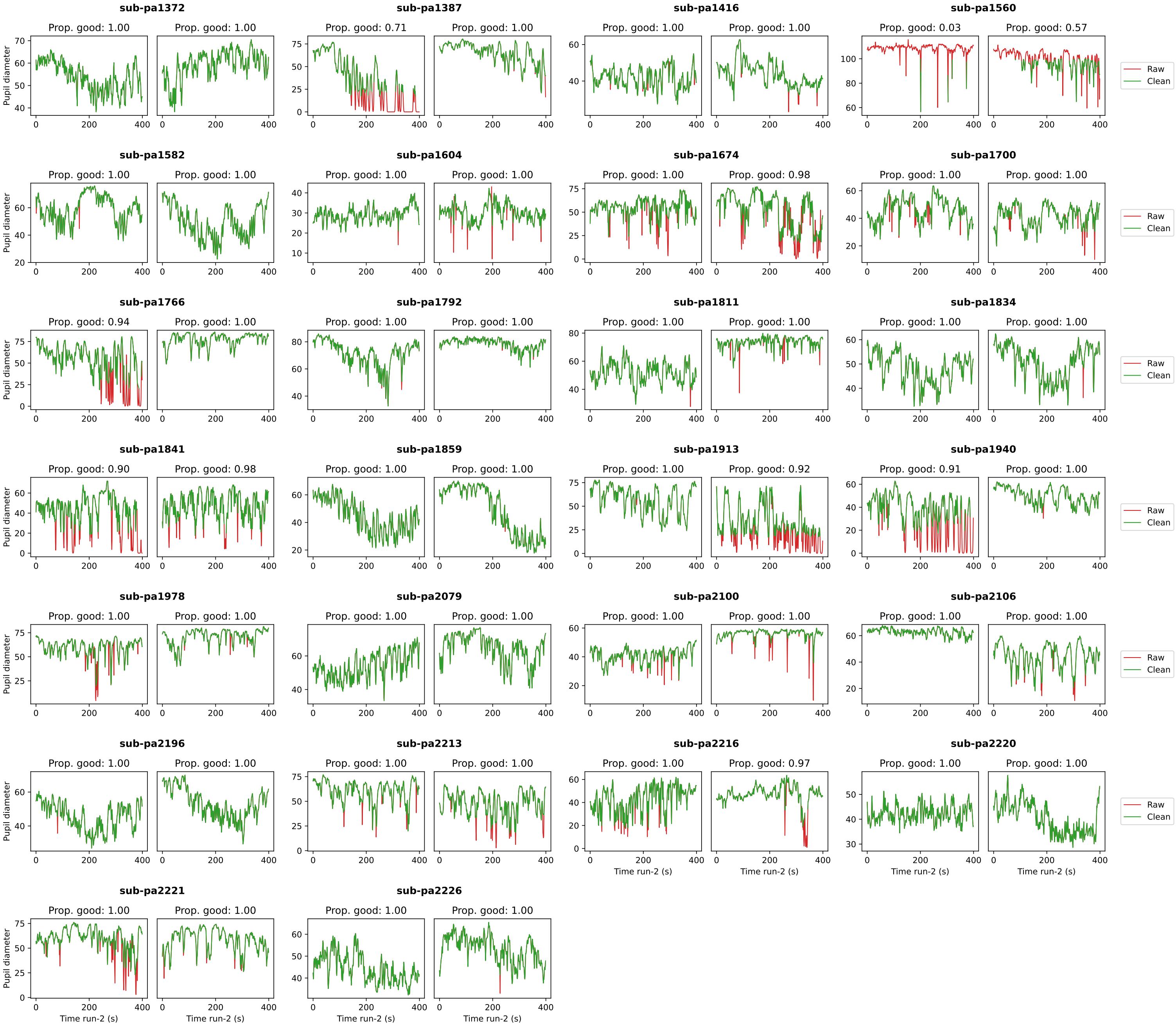

sub-pa1372
Prop. good: 1.00
Prop. good: 1.00
sub-pa1387
Prop. good: 0.71
Prop. good: 1.00
sub-pa1416
Prop. good: 1.00
Prop. good: 1.00
sub-pa1560
Prop. good: 0.03
Prop. good: 0.57
sub-pa1582
Prop. good: 1.00
Prop. good: 1.00
sub-pa1604
Prop. good: 1.00
Prop. good: 1.00
sub-pa1674
Prop. good: 1.00
Prop. good: 0.98
sub-pa1700
Prop. good: 1.00
Prop. good: 1.00
sub-pa1766
Prop. good: 0.94
Prop. good: 1.00
sub-pa1792
Prop. good: 1.00
Prop. good: 1.00
sub-pa1811
Prop. good: 1.00
Prop. good: 1.00
sub-pa1834
Prop. good: 1.00
Prop. good: 1.00
sub-pa1841
Prop. good: 0.90
Prop. good: 0.98
sub-pa1859
Prop. good: 1.00
Prop. good: 1.00
sub-pa1913
Prop. good: 1.00
Prop. good: 0.92
sub-pa1940
Prop. good: 0.91
Prop. good: 1.00
sub-pa1978
Prop. good: 1.00
Prop. good: 1.00
sub-pa2079
Prop. good: 1.00
Prop. good: 1.00
sub-pa2100
Prop. good: 1.00
Prop. good: 1.00
sub-pa2106
Prop. good: 1.00
Prop. good: 1.00
sub-pa2196
Prop. good: 1.00
Prop. good: 1.00
sub-pa2213
Prop. good: 1.00
Prop. good: 1.00
sub-pa2216
Prop. good: 1.00
Prop. good: 0.97
sub-pa2220
Prop. good: 1.00
Prop. good: 1.00
sub-pa2221
Prop. good: 1.00
Prop. good: 1.00
sub-pa2226
Prop. good: 1.00
Prop. good: 1.00
Pupil diameter
Time run-2 (s)
Raw
Clean

#### **Fig. S17**: Quality control of YRSP physiological data (pupil diameter over time)

Raw (red) and cleaned (green) pupil diameter time series for both runs of each participant in the YRSP dataset. Panel titles indicate the proportion of valid data points after artifact removal. Sessions with fewer than 80% valid data points were excluded from analyses. Abbreviations: YRSP: Yale Resting-State Pupillometry dataset.

**Fig. S18**

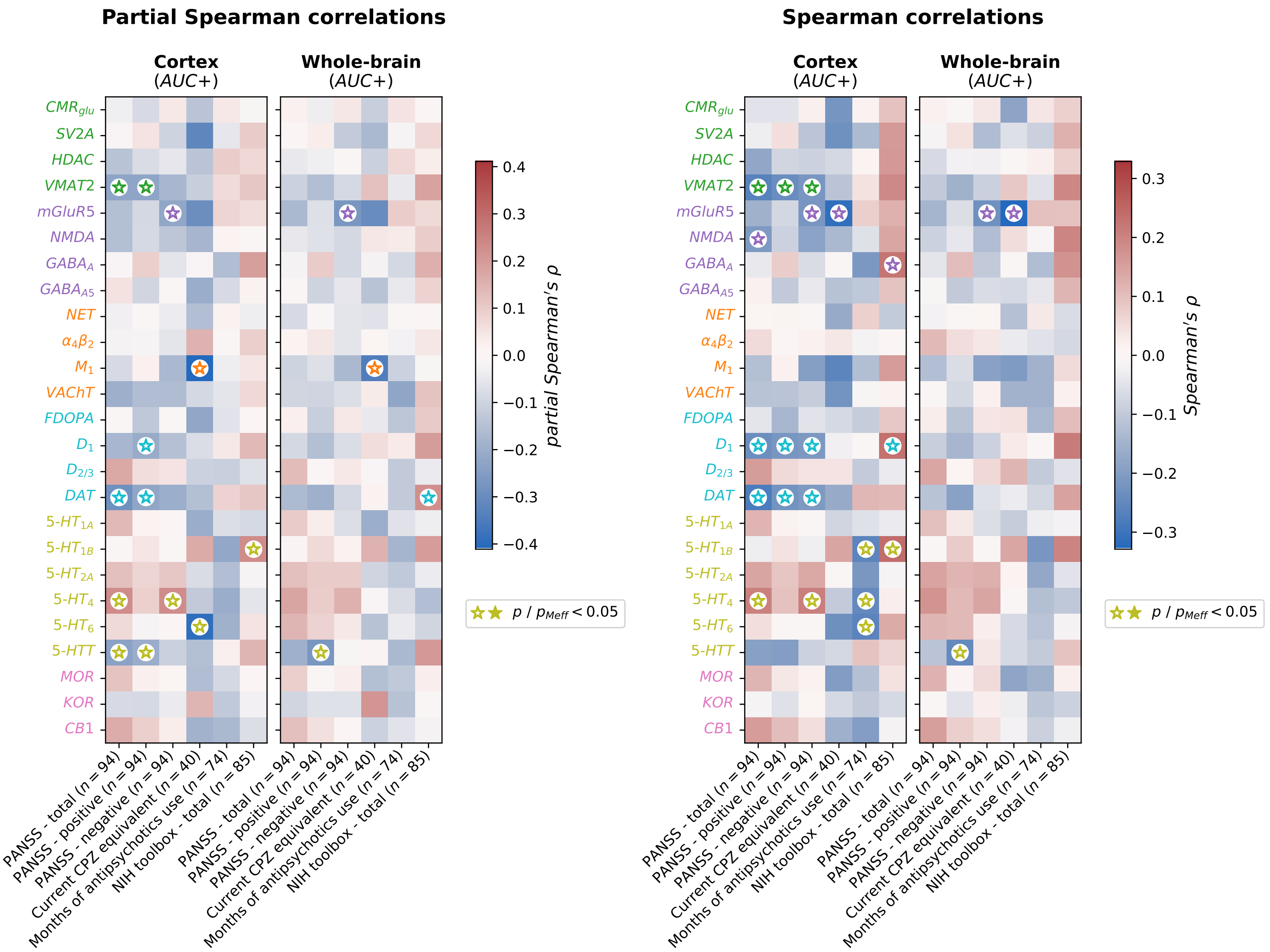

Partial Spearman correlations
Spearman correlations
Cortex (AUC+)
Whole-brain (AUC+)
CMR_glu
SV2A
HDAC
VMAT2
mGluR5
NMDA
GABA_A
GABA_A5
NET
α4β2
M1
VAChT
FDOPA
D1
D2/3
DAT
5-HT1A
5-HT1B
5-HT2A
5-HT4
5-HT6
5-HTT
MOR
KOR
CB1
PANSS - total (n = 94)
PANSS - positive (n = 94)
PANSS - negative (n = 94)
Current CPZ equivalent (n = 40)
Months of antipsychotics use (n = 74)
NIH toolbox - total (n = 85)
partial Spearman's ρ
Spearman's ρ
p / p_Meff < 0.05

#### **Fig. S18**: Associations of AUC scores with clinical variables

Associations of NEOFC AUC+ and AUC– scores (Schaefer200 and Schaefer200 + subcortex parcellations) with six clinical and cognitive outcomes in the psychosis group. Both Spearman and partial Spearman correlations were computed, adjusting for sex, age, mean FD, global connectivity, and current antipsychotic dose. Correlations with antipsychotic outcomes were calculated without the antipsychotic covariate. Abbreviations: AUC: area under the curve; PANSS: Positive and Negative Syndrome Scale; CPZ: chlorpromazine; FD: framewise displacement; Meff: effective number of tests.
