## Supplementary figures and images for "Linking human brain functional connectivity to underlying neurotransmitter systems"

### Supplementary Animation S1

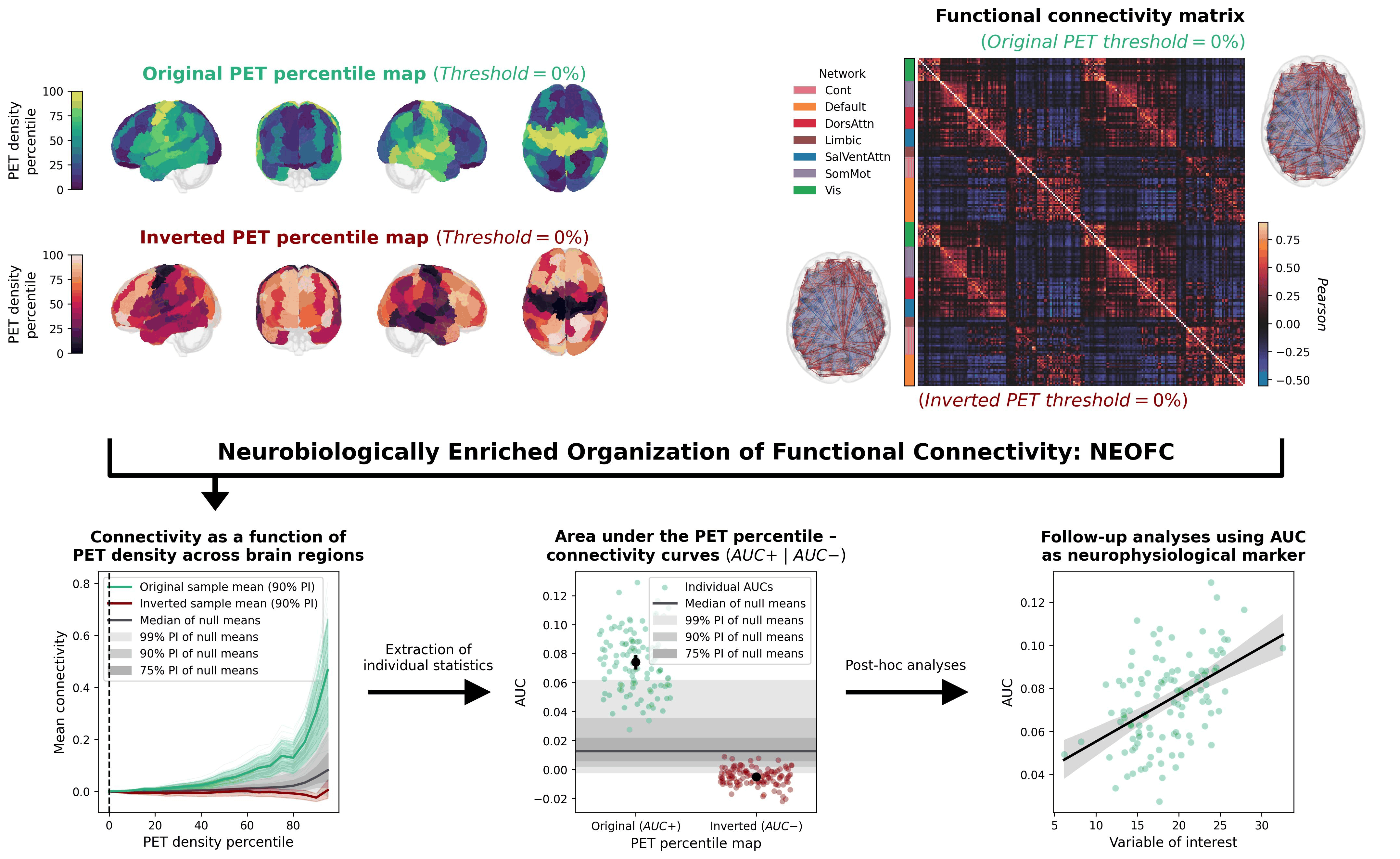
